# Reduction of forebrain interneurons alters cellular organization and cortical neurogenesis in a mouse model of X-linked intellectual disability

**DOI:** 10.64898/2026.09.08.749487

**Authors:** Jianing Li, Vedant Garg, Ane Martiartu, Anthony Tanzillo, Yajun Zhang, Timothy J. Petros

## Abstract

Arx is an X-linked transcription factor that is expressed in inhibitory neurons within the ganglionic eminences during embryogenesis. In humans, *ARX* variants are one of the leading causes of X-linked intellectual disabilities (XLID) and related syndromes. While most often studied in males, recent evidence indicates that >50% of heterozygous female carriers of *ARX* variants also display mild to severe phenotypes. The mechanisms by which *Arx* mutations lead to a wide range of disease pathologies remains poorly understood. Here we removed Arx in the medial ganglionic eminence (MGE) to generate conditional knockout male and heterozygous female mice. MGE-derived interneurons were reduced in a gene dosage, subtype and brain region dependent manner in Arx mutant mice. Knockout male mice displayed severe phenotypes with infantile spasms and epilepsy, while Het female mice displayed moderate anxiety and locomotor defect mimicking patient symptoms. Single cell sequencing and spatial transcriptomics revealed striking cell autonomous and non-cell autonomous changes throughout development. Notably, loss of MGE-derived interneurons alters dorsal cortical neurogenesis and cell-cell communication resulting cortical malformations. This study reveals mechanistic insights into genetic and cellular changes that occur when Arx is removed from MGE-derived interneurons that advances our understanding of Arx-related neurodevelopmental disorders.

## INTRODUCTION

The X-chromosome contains ∼5% of the human genome, yet ∼15% of genes (> 100 genes) associated with intellectual disability (ID) are located on the X-chromosome^1,2^. This X-linked recessive inheritance of ID-related genes in large part explains why the prevalence of ID is ∼50% higher in males^3^. X-linked intellectual disabilities (XLIDs) are a complex and heterogeneous set of syndromes, with ∼50% of these disorders being comorbid with seizures or epilepsy^4^. A better understanding of the genetic mechanisms underlying XLIDs is necessary to understand these disorders and develop new therapeutic treatments.

One of the most frequently mutated genes in XLIDs is the transcription factor Aristaless-related homeobox, Arx. Males with *ARX* variants display wide ranging symptoms including ID, epilepsy, lissencephaly, and autism-like phenotypes. The most severe syndromes involving brain and genital malformations, (X-linked lissencephaly associate with abnormal genitalia, XLAG, and Proud Syndrome) often arise from point mutations or deletions near the homeodomain while non-malformation phenotypes (Partington Syndrome and West Syndrome) arise from polyalanine expansions and duplications outside of the homeodomain region^5–8^. While most clinical and scientific studies on Arx function focus on males, recent evidence indicates that ∼60% of female heterozygous carriers of *ARX* variants display mild to severe XLID-like symptoms^9^. Some of this variability is likely due to X chromosome inactivation (XCI). There is a bias of paternal XCI in the brain, and evidence that local XCI mosaicism of WT vs. mutant alleles can correlate with disease penetrance^10^.

The comorbidity of ID and epilepsy implicates abnormal function of GABAergic interneurons in XLIDs and *ARX* variants. Indeed, *Arx* is strongly upregulated in postmitotic neurons within the medial, lateral and caudal ganglionic eminences (MGE, LGE & CGE, respectively)^11,12^, the source of nearly all GABAergic interneurons and projection neurons in the forebrain. Several studies found that loss of *Arx* in the GEs disrupts normal migration of interneuron precursors^11,13,14^, leading to reduced cortical interneurons^14–16^ and increased seizures^17^. Notably, postnatal removal of *Arx* from parvalbumin-expressing (PV+) interneurons altered network excitability and induced spontaneous seizures^18^, indicating that *Arx* plays an important role in interneuron function in adult mice as well. Other studies generated mice mimicking human *ARX* variants that display variations in symptoms and phenotype severity^19–24^. Additionally, perturbation of *ARX* in organoids and human primary fetal neuronal cultures recapitulate some of these migration defects and cell fate changes^25,26^. Despite this progress, the mechanisms by which Arx mutations lead to disease pathology remain poorly understood.

To characterize how loss of Arx induces cell autonomous and non-cell autonomous changes in interneurons throughout development, we generated conditional knockout male and heterozygous female mice where *Arx* is specifically removed from the MGE. While male KO mice have a near complete loss in MGE-derived interneurons in the cortex and hippocampus, female Het mice show a ∼50% reduction. Arx cKO male mice die by 6 weeks, but Arx Het females display increased in seizure susceptibility and mild impairment in locomotor activity and sensorimotor gating, consistent with human female heterozygous carriers of *ARX* variants^9^. We combined single nuclei Multiome analyses and spatial transcriptomics to characterize striking cell autonomous and non-cell autonomous changes in the embryonic and adult brains of Arx cKO males and het female mice. Loss of Arx in the MGE produces a distinct, transcriptionally defined cohort of cells in male cKO, while females contain both normal and aberrant neuronal populations. Arx-deficient MGE-derived interneurons also exert non-cell-autonomous effects on cortical progenitor cells and Cajal-Retzius cells, resulting in a thinner cortex and abnormal cortical lamination. In the adult, we observe cellular and transcriptional changes associated with epilepsy, especially in the hippocampus. This comprehensive dataset provides a valuable resource for understanding how loss of Arx effects gene expression, cell fate and behavior in relation to human XLIDs.

## RESULTS

### MGE-derived interneuron numbers decline in a Arx dosage-dependent manner

We generated conditional *Nkx2.1-Cre^C/+^;Arx^F/Y^;Ai9^F/+^*knockout males (cKO) and *Nkx2.1-Cre^C/+^;Arx^F/+^;Ai9^F/+^*heterozygous females (Het) to remove *Arx* from the MGE, with Nkx2.1-lineage cells expressing the red fluorescent reporter tdTomato. We confirmed that Arx protein is strongly downregulated in the MGE (Figure S1A). Most cortical MGE-derived interneurons express Arx at E13.5 and P0, but Tom+ cells in the cortex of cKO male mice were Arx-negative (Figure S1A-B). Arx cKO male mice exhibited reduced body and brain weights, with most dying before weaning, whereas Het females had normal body weight, brain weight and viability (Figure S2A). The MGE gives rise to nearly all parvalbumin– and somatostatin-expressing interneurons (PV+and SST+, respectively) in the cortex, as well as a population of neuronal nitric oxide synthase-expressing (nNos+) neurogliaform and ivy cells in the hippocampus. At P21, we observed a near complete loss of MGE-derived interneurons in the cortex, with significantly decreased density and percentage of PV+ and SST+ interneurons in Arx cKO males (Figure 1A-B). Note that most Tom+ cells in the cortex of Arx cKO males do not express PV or SST, and conversely most PV+ and SST+ cells are Tom-negative. In the *Nkx2.1-Cre* mouse, Cre is not efficiently expressed in the most dorsal portion of the MGE^27^, so these PV+/Tom– and SST+/Tom-cells likely originate from this region where Cre is not expressed, and thus Arx expression is presumably normal in these cells. Arx Het females exhibited a ∼50% reduction in PV+ and SST+ interneuron numbers, but their relative proportions of these subtypes remained unchanged compared to WT (Figure 1A-B). We observe a ∼2:1 ratio of PV+:SST+ cells in WT and Het females, but this ratio is reversed in cKO male mice (Figure 1C). This decreased tendency and altered proportion in MGE-derived cells was present in both the deep and superficial cortical layers (Figure S3).

**Figure 1.**
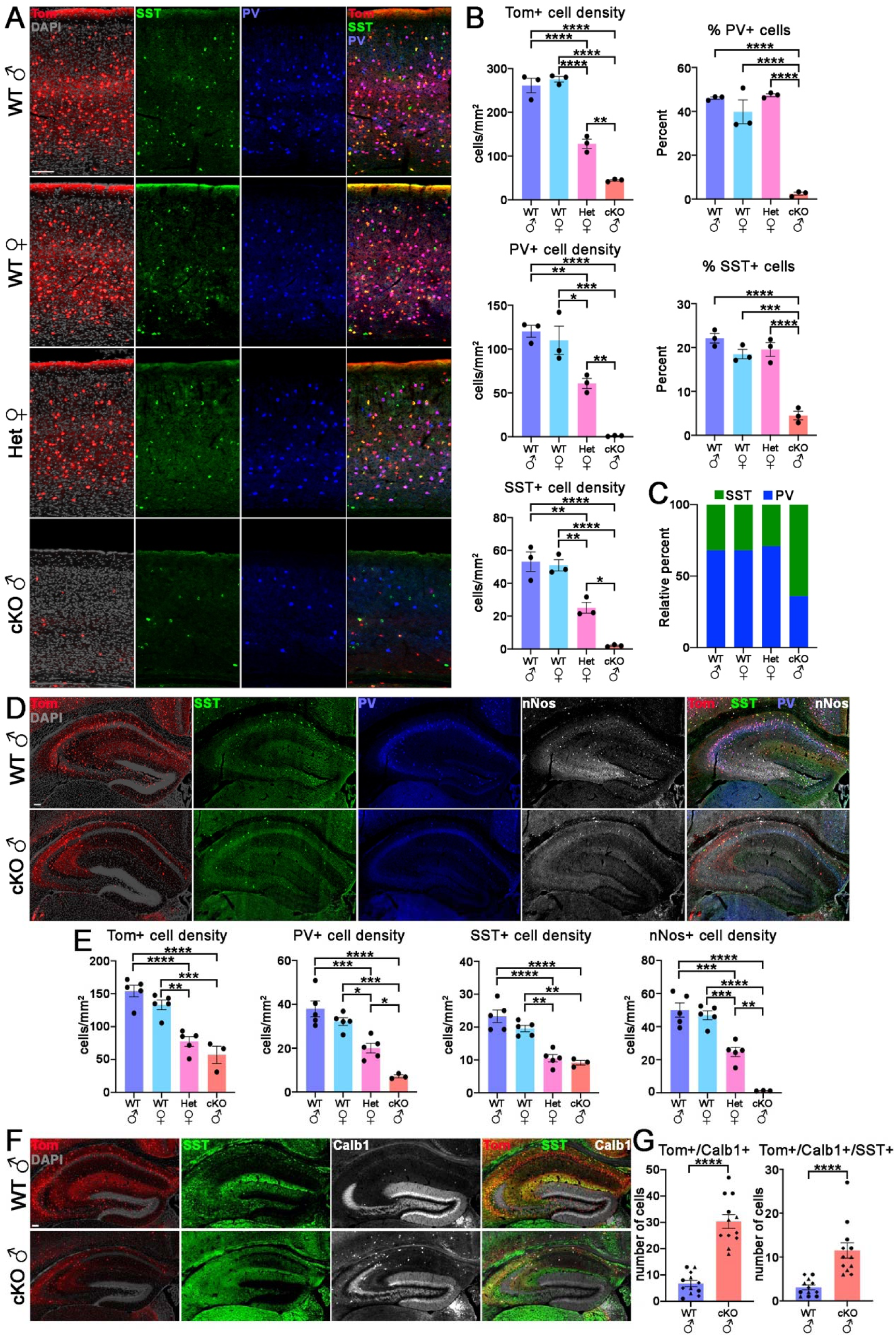
MGE-derived interneuron loss in a Arx dosage-dependent manner. **A.** P21 somatosensory cortex from *Nkx2.1-Cre*;*Arx*;*Ai9* WT male and female, Het female and cKO male mice stained for tdTomato, PV, SST and DAPI. Scale bar = 50 μm. **B.** Quantification of cell density (left) and percent of PV+ and SST+ cells (right) in the somatosensory cortex. n = 3 for each condition. **C.** Average relative percent of PV+ and SST+ cells in the somatosensory cortex. **D.** P21 hippocampus in WT and cKO male mice stained for PV, SST and nNos. Scale bar = 100 μm. **E.** Quantification of cell density of PV+, SST+ and nNos+ cells in the hippocampus. n = 5 WT male, 5 WT female, 5 Het female and 3 cKO male mice. **F.** P21 hippocampus in WT and cKO male mice stained for SST and Calb1. Scale bar = 100 μm. **G.** Quantification of cell density of Tom+/Calb1+ and Tom+/Calb1+/SST+ cells in the hippocampus. n = 3 WT and cKO brains, 12 sections/brain, each brain labeled with a different shape. All stats are one-way ANOVA followed by Tukey’s multiple comparison tests (B, E), or Welch’s 2-tailed t-test (G: left) and Mann-Whitney 2-tailed test (G: right) based on normal distribution when only WT and cKO mice tested: * = p ≤ .05, ** = p ≤ .01, *** = p ≤ .001, **** = p ≤ .0001.

There was also a strong reduction of MGE-derived interneurons in the hippocampus of Arx cKO males, but less extreme compared to the cortex. The overall number of Tom+ cells was similar between Arx Het females and cKO males, ∼50% reduction compared to WT mice (Figures 1D-E & S4A-B). nNos+ cells were nearly absent in the Arx cKO hippocampus whereas the number of SST+ interneurons was similar between Arx Het females and cKO males (Figures 1D-E & S4A-B). The PV+ presynaptic marker synaptotagmin-2 (Syt2)^28^ was strongly decreased throughout the hippocampus of Arx cKO males (Figure S2B). This distribution of interneuron loss was not uniform across hippocampal regions: Tom+ cell loss was most pronounced in the dentate gyrus (DG) relative to CA1 and CA2/3 regions of Arx cKO males (Figure S4C-E). We observed a significant increase in the number of Tom+/Calb1+ interneurons in the hippocampus of Arx cKO males, with many of these cells co-expressing SST (Figure 1F-G). The MGE gives rise to a population of oligodendrocytes in CA3 that perdure into adulthood^29,30^, and the density of these oligodendrocytes is unaffected in Arx cKO males (Figure S2C-D). In sum, forebrain MGE-derived interneurons were reduced in a gene dosage, subtype, and region dependent manner in Arx Het female and male cKO mice.

### Increased seizure sensitive and moderate impairment in locomotor and sensorimotor gating of Arx Het mice

Most Arx cKO male mice exhibited spontaneous seizures from P10-P21, characterized by severe forelimb clonus and rearing with hypoactivity. These seizures are likely a leading cause of their premature death, which prevented us from performing any behavior assays on these mice. Over half of human female carriers of *ARX* variants display seizures, ID and other phenotypes^9,17^, and we observed spontaneous seizures in a few Het female mice. To assess seizure susceptibility in Het female mice, we induced seizures with a single injection of Pentylenetetrazole (PTZ). Het female mice exhibited increased seizure scores and seizure severity compared to WT mice, indicating increased seizure susceptibility in Het females (Figure 2A).

**Figure 2.**
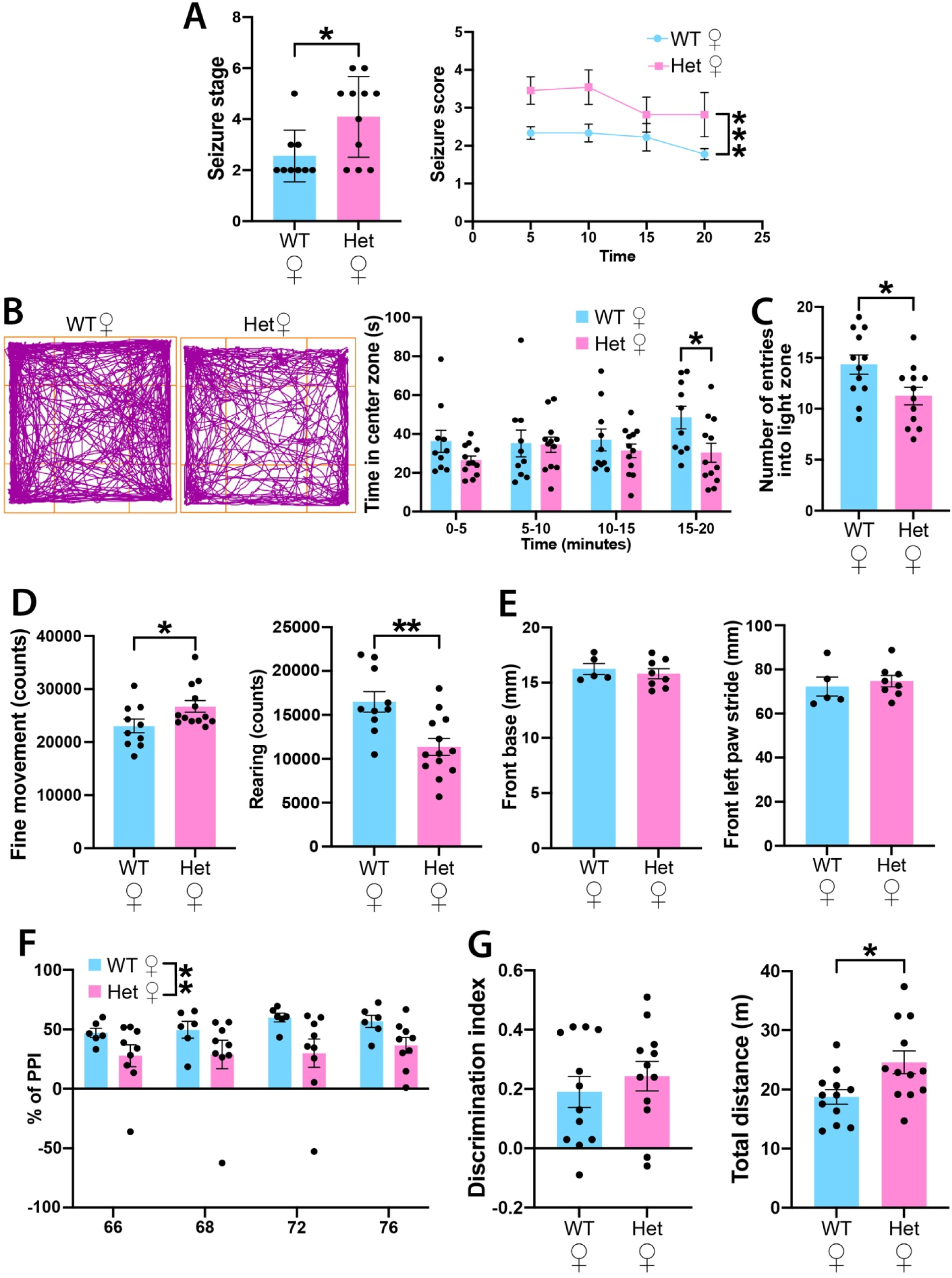
Increased seizure sensitivity and impaired locomotor and sensorimotor gating of Arx Het mice. **A.** Maximum seizure scores for WT (n = 9) and Het (n = 11) females following PTZ injection (left), and during the 20-minute observation period (right). **B.** Representative trajectory plots (left) and time in the center zone (right) of female WT (n = 10) and Het (n = 12) mice during the open field test. **C.** Number of entries into light zone of WT (n = 12) and Het (n = 12) mice in the light-dark box. **D.** Beam break counts of fine movements (left) and rearing (right) of WT (n = 10) and Het (n = 13) in home cage test. **E.** Front base (left) and front left paw stride (right) of WT (n = 5) and Het (n = 8) mice in the free walk test. **F.** Percent of prepulse inhibition (PPI) in WT (n = 6) and Het (n = 9) mice. **G.** Discrimination index (left) and total distance traveled (right) in the 2-object novel object recognition test of WT (n = 12) and Het (n = 12) mice. Stats are ordinary two-way ANOVA (A: right, F) followed by Sidak’s multiple comparisons test (B), Mann-Whitney 2-tailed test (A: left) and unpaired 2-tailed t-test (C, D, E, G) based on normal distribution: * = p ≤ .05, ** = p ≤ .01, *** = p ≤ .001.

We performed a series of behavioral tests to assess phenotypes associated with neurodevelopmental disease in WT and Het female mice. To assess anxiety-like behaviors, we performed the open field test and light-dark box test. Het female mice spent less time in the center zone during the open field test, with this difference reaching significance during the final 5-minute interval (Figure 2B). In the light-dark box test, Het female mice exhibited a significant decrease in the number of entries of the light zone (Figure 2C). To assess general locomotor activity, we continuously recorded singly housed mice using the Photobeam Activity System over four days. Het female mice exhibited an increase in fine movements with a decrease in rearing movements compared to WT mice (Figure 2D), mimicking dystonic movements in human patients with *ARX* variants^7^. Gait and motor coordination was assessed in a free-walk test, but no significant differences were observed between WT and Het female mice in any parameter measured (Figure 2E).

We then assessed sensorimotor gating and memory behaviors in Het female mice. Sensorimotor gating prevents the brain from overwhelmed by environmental stimuli and reflect the ability to suppress a motor response^31^. Prepulse inhibition (PPI) of the startle response assesses sensorimotor gating, and deficits in PPI are associated with schizophrenia and ADHD^31,32^. Het female mice exhibited a significantly reduced PPI percentage compared to WT mice, indicating deficits in sensorimotor processing (Figure 2F). To assess memory deficits, we employed the 2-object novel object recognition test. Although no significant difference in discrimination index was observed between groups, Het female mice traveled a greater total distance during the task compared to WT controls (Figure 2G), potentially reflecting hyperactivity and/or heightened anxiety, consistent with results from the open field and light-dark box tests. In summary, Het female mice display behavioral deficits including increased seizure susceptibility, mild anxiety-like behavior, and impairments in locomotor activity and sensorimotor gating, as observed in human female carriers.

### Building a spatiotemporal transcriptome atlas in Arx mutant mice

The above findings reveal that Arx cKO male and Het female mice represent two levels of disease phenotypes. Arx cKO mice recapitulate severe clinical features of human *ARX* variants including postnatal microcephaly, intractable epilepsy, and early lethality^7^. Het female mice, by contrast, display a moderate phenotype consistent with human female carriers of *ARX* variants, characterized by mild to moderate intellectual disability, stereotyped hand movements, and dystonia, deficits that are associated with dysfunction of the cerebral cortex, hippocampus, and basal ganglia^7,17^. To elucidate the genetic mechanisms underlying these two distinct phenotypes, we constructed a transcriptomic atlas spanning developmental stages through adulthood using single nuclei Multiome sequencing (snRNA-seq and snATAC-seq) and spatial transcriptomics (Figure 3A).

**Figure 3.**
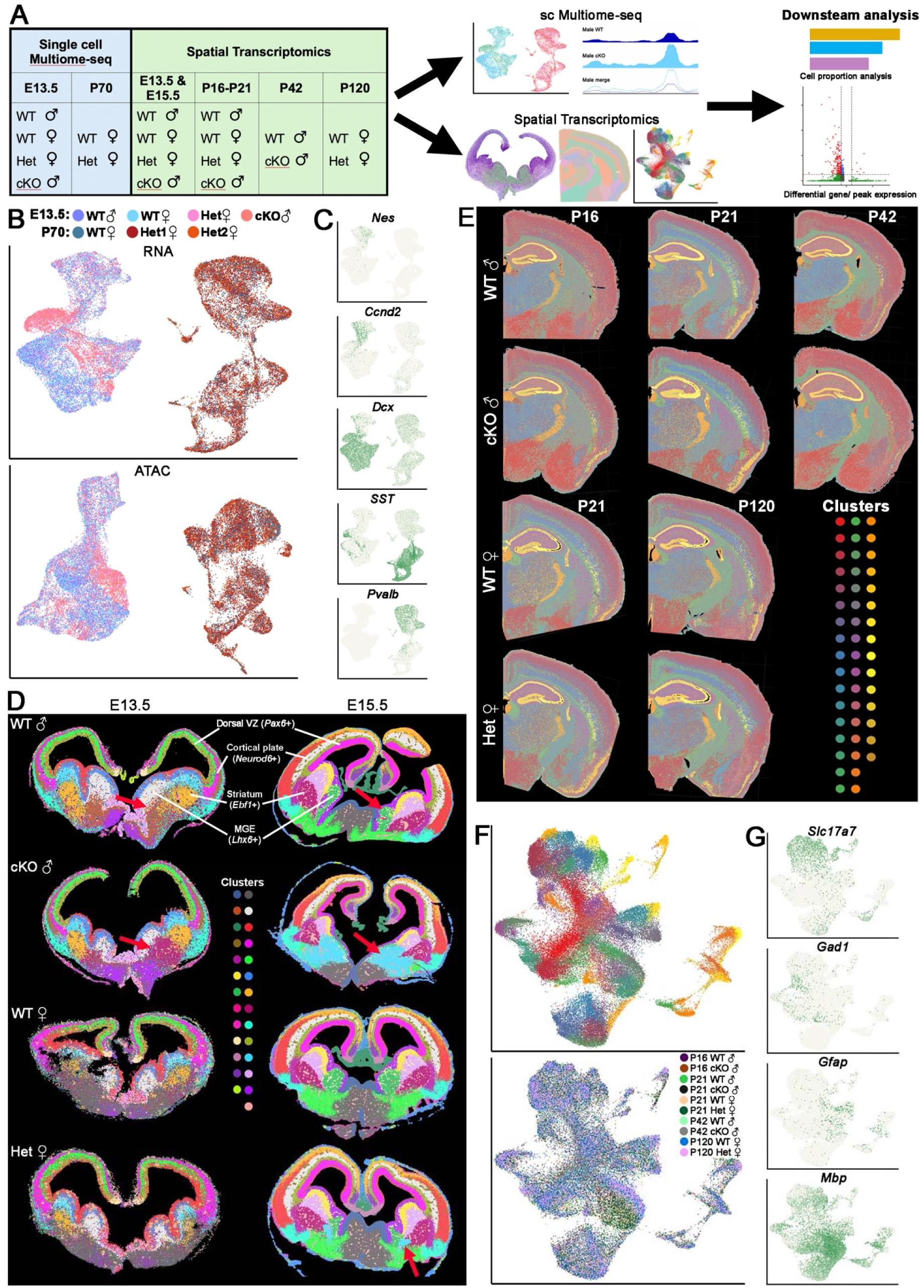
Spatiotemporal transcriptome atlas in Arx mutant mice. **A.** Summary of ages & genotypes for snMultiome-seq and spatial transcriptomics workflow. **B.** UMAP plots of snRNA (top) and snATAC (bottom) data of E13.5 and P70 Nkx2.1-lineage cells annotated by age, gender and genotype. **C.** UMAP plots of snRNA annotated by marker genes for apical progenitors (*Nestin*), basal progenitors (*Ccnd2*), post-mitotic immature neurons (*Dcx*), and mature SST+ (*Sst*) and PV+ (*Pvalb*) interneurons. **D.** E13.5 and E15.5 spatial transcriptomics data clustered by gene expression profiles that correspond to defined anatomical regions. E13.5 and E15.5 datasets were individually analyzed, and colors only matched in individual timepoint across genotypes. **E.** Postnatal spatial transcriptomics brain slices colored by molecular clusters based on gene expression profiles. **F.** UMAP plots (downsampled to 5,000 spots per tissue) of postnatal spatial transcriptome slices annotated by clusters (top) and samples (bottom). Cluster colors in UMAP are same as in panel E. **G.** UMAP plots annotated by marker genes for excitatory neurons (*Slc17a7+*), inhibitory neurons (*Gad1+*), astrocytes (*Gfap+*) and oligodendrocytes (*Mbp+*).

We generated WT, Het female and cKO male *Nkx2.1-Cre;Arx^F^;Sun1-sfGFP* mice to harvest Nkx2.1+ lineage GFP+ nuclei via flow cytometry. We performed single nuclei Multiome sequencing on MGE from E13.5 WT male, cKO male, WT female and Het female mice. We also harvested Nkx2.1+ lineage nuclei from the cortex of P70 WT and Het females for Multiome sequencing, but the lack of MGE-derived cortical interneurons in adult cKO male mice prevented similar analyses. 18,758 E13.5 nuclei and 16,214 P70 nuclei passed quality control and were used for analysis (Supplementary Table 1). Visualization of RNA-only and ATAC-only data revealed clear segregation between embryonic MGE and adult cortex datasets (Figure 3B). Distinct neurogenic cell populations such as apical progenitors (*Nes*+), basal progenitors (*Ccnd2*+), and postmitotic neurons (*Dcx*+) were visible in the E13.5 dataset, whereas the two principal MGE-derived interneuron subtypes, *Pvalb*+ and *Sst*+, were clearly segregated in the P70 dataset (Figure 3C).

To characterize how loss of *Arx* in MGE-derived cells alters gene expression in all cell types throughout development, we generated spatial transcriptomics datasets spanning embryonic development through adulthood (Figure 3A & Supplementary Table 1). Spatial transcriptomics was performed on all sexes and genotypes at E13.5, E15.5 and postnatal days 16-21. We also performed spatial transcriptomics in male cKO mice at P42, the oldest age we obtained, and female Het mice at P120, following the behavior assays described above. In the E13.5 and E15.5 brain, we identified 27 cell clusters based on gene expression profiles that correspond to defined anatomical regions, such as the dorsal cortical ventricular zone (*Pax6*+), cortical plate (*Neurod6*+), future striatum (*Ebf1*+) and MGE (*Lhx6*+) (Figure 3D). We observed a striking difference in cluster composition in the postmitotic mantle region of the MGE of male cKO mice compared to WT in both ages (Figure 3D).

For the P16-P120 brain sections, 49 cell clusters were generated based on gene expression profiles with a clear correspondence to anatomical regions and cell types (Figure 3E). We then downsampled each section to 5,000 spots to facilitate UMAP visualization (Figure 3F). Within these datasets, we identified several major cell types, including excitatory neurons (*Slc17a7*+), inhibitory neurons (*Gad1*+), astrocytes (*Gfap*+) and oligodendrocytes (*Mbp*+) (Figure 3G). Thus, we could identify the expected developmental trajectory and cell types in the single nuclei Multiome and spatial transcriptomics dataset from embryogenesis through adulthood.

### Ectopic cell population with distinct genetic signatures in the MGE of Arx mutant mice revealed by spatiotemporal transcriptomics

We subsetted the E13.5 MGE Multiome data and integrated the snRNA and snATAC datasets using weighted nearest neighbor (WNN). Postmitotic cells in male cKO mice were completely segregated from postmitotic cells in male WT mice, with female Het mice having cells in both the endogenous and ectopic clusters (Figures 4A-B & S5A-B). We defined cell clusters based on their gene expression profiles, revealing 4 progenitor cell types, 14 postmitotic cell types, and 1 cluster of oligodendrocyte precursors (Figures 4B & S5A). There were no changes in the proportion of progenitor cell types in male or female Arx mutant mice, likely due to *Arx* expression being enriched in postmitotic MGE neurons^11^ (Figure 4C).

**Figure 4.**
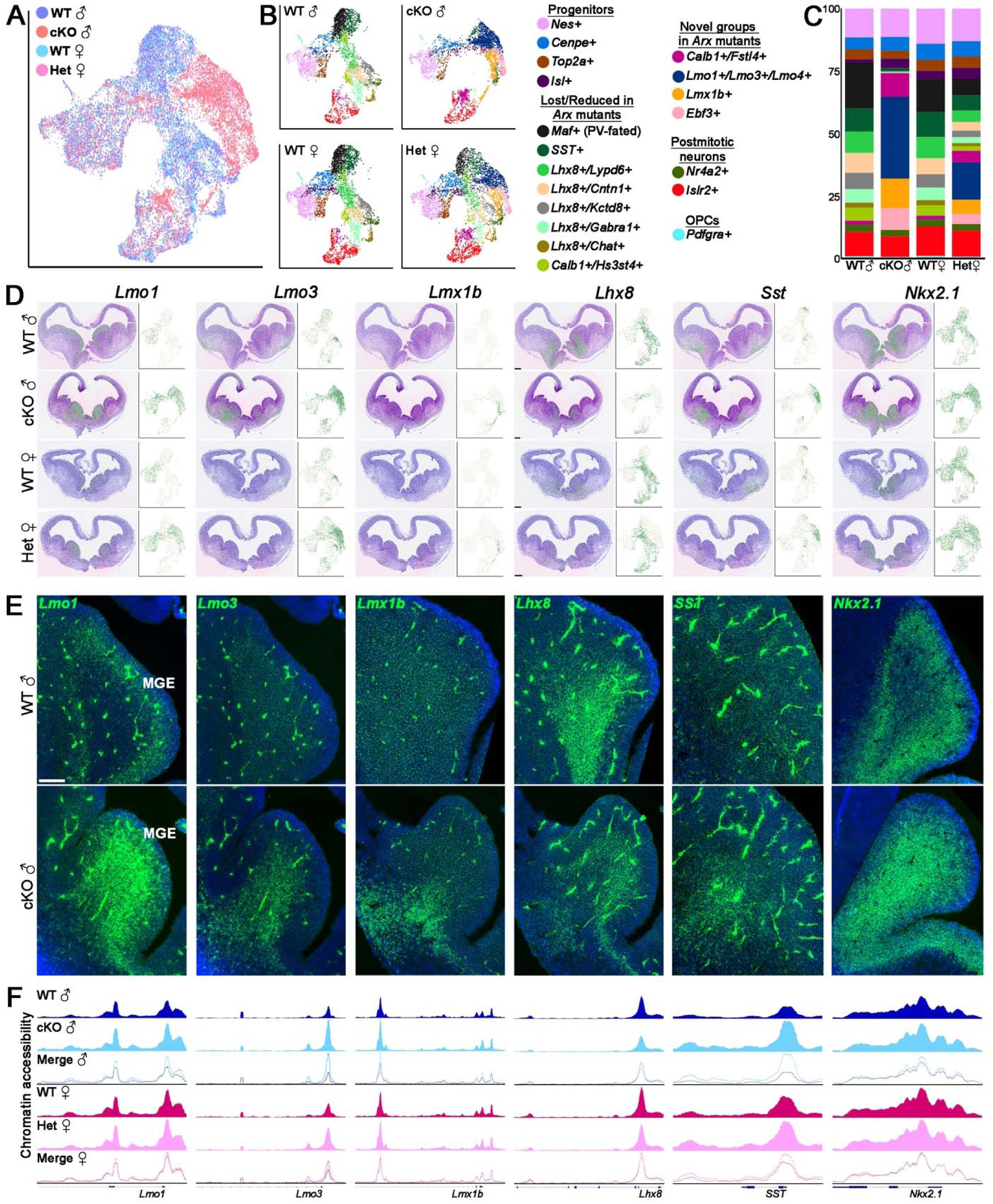
Arx dose-dependent alteration of MGE cell fate at E13.5. **A-B**. UMAP plots of integrated snRNA and snATAC data via weighted nearest neighbor (WNN) of Nkx2.1-lineage cells from E13.5 MGE, annotated by genotype (A) and inferred cell clusters based on gene expression profiles (B). **C.** Relative proportions of E13.5 MGE cell clusters of WT, Het and cKO mice. **D-E.** mRNA expression of *Lmo1, Lmo3, Lmx1b, Lhx8, Sst,* and *Nkx2.1* in E13.5 brains visualized in spatial transcriptomics data (D, left), UMAP plot from snMultiome data (D, right) and FISH (E). Scale bar = 200 μm (D) or 100 μm (E). **F.** Tracks showing accessible chromatin peaks from snATAC data (pseudobulk) for individual genotypes at the *Lmo1, Lmo3, Lmx1b, Lhx8, Sst* and *Nkx2-1* genomic loci.

In male cKO mice, we observe a near complete loss of *Maf*+ cells, which are predictive of PV-fated interneurons^33^, and a strong reduction of normal SST-fated interneuron cluster (Figure 4B-C); this is consistent with the downregulation of these interneurons in cKO male mice. Nearly all *Lhx8*+ clusters, which predominantly give rise to MGE-derived striatal interneurons and projection neurons comprising the globus pallidus and other basal ganglion nuclei^34–36^, are lost in male cKO mice (Figure 4B-C). These *Lhx8*+ cells are replaced by cell clusters identified by strong upregulation of several Lim homeodomain transcription factors (*Lmo1*, *Lmo3*, *Lmo4*, *Lmx1b*) and the transcription factor *Ebf3* (Figure 4B-C). We identified two clusters expressing calbindin (*Calb1*+), one of which is completely lost (*Hs3st4*+) while the other is expanded (*Fstl4*+) in male cKO mice (Figure 4B-C). Whether this expanded *Calb1*+ population in the MGE gives rise to the increased Calb1+ interneuron population in the hippocampus of adult male cKO mice (Figure 1F-G) remains unknown. The MGE of female Het mice contained both the normal and ectopic clusters, with ∼50% reduction of endogenous *Maf*+, *Sst*+ and *Lhx8*+ clusters compared to female WT mice, and the proportion of mutant-specific ectopic clusters is ∼50% compared to male cKO mice (Figure 4A-C). These results are consistent with the ∼50% reduction of PV+ and SST+ interneurons in adult female Het mice.

Most differentially expressed genes (DEGs) were increased in the MGE of both male cKO and female Het MGE (Figure S6A), consistent with Arx functioning primarily as a transcriptional repressor^37–39^. Markers of the ectopic cell clusters (*Lmo1*, *Lmo3*, *Lmo4*, *Lmx1b*, *Ebf3*) were some of the strongest upregulated genes in male cKO mice (Figure S6A). Several genes associated with cell differentiation and fate commitment (*Etv1, Pbx1, Pbx3*)^40–43^, alternative splicing (*Rbfox1, Nova1*), and epilepsy and neurodevelopmental delay (*Kcnq3*, *Glyctk, Magel2*)^44–46^ were also significantly upregulated in the MGE of male cKO and female Het mice (Figure S6A-B). In contrast, genes associated with GABAergic neuron identity (*Lhx8*) and PV+ interneuron specification (*Maf*) were significantly downregulated in *Arx* mutant mice (Figure S6A). Many of these genes also had corresponding differentially accessible peaks (DAPs) (Figure S6C), indicating a clear correlation between chromatin accessibility and gene expression in these mice.

Several genes involved in interneuron migration were altered in Arx mutant mice. Neuregulin 1 (Nrg1), Ackr3 (Cxcr7) and Cxcr4 are critical regulators of tangential interneuron migration to the cortex^41,47,48^. *Nrg1* was one of the strongest upregulated genes in Arx mutant MGE while *Ackr3* and *Cxcr4* were both downregulated in Arx mutants (Figure S6A). Dysregulation of these guidance factors are likely causes for abnormal interneuron migration in the ventral forebrain.

Complementing this Multiome data with spatial transcriptomics provided additional insight into gene expression changes in *Arx* mutant mice. While *Lmo1* is expressed in cycling MGE progenitors in WT mice, *Lmo1* is strongly upregulated in postmitotic cells in the MGE mantle in male cKO mice and appears to be downregulated as these cells migrate out of the MGE (Figure 4D). *Lmo3* is upregulated in cells slightly deeper in the mantle and is maintained in the *Islr2*+ and *Calb1*+*/Fstl4*+ postmitotic neuron clusters (Figure 4D). *Lmx1b*+ is specifically upregulated in the deepest portion of the mantle, almost complementary to *Lmo1* (Figure 4D). These observations were confirmed with FISH (Figures 4E & S7). We also observed a concurrent increase in chromatin accessibility at the promoters of *Lmo1*, *Lmo3* and *Lmx1b* in male cKO mice compared to WT, with the largest difference at *Lmo3* (Figures 4F). Female Het mice displaying a more moderate increase in accessibility at these loci. Conversely, *Lhx8* was strongly reduced in the MGE mantle of male cKO mice, concomitant with reduced accessibility at the *Lhx8* promoter, but *Lhx8* expression was retained in the subventricular zone (SVZ) (Figure 4D-F).

Despite the reduction of MGE-derived interneurons, we observed a significant increase in *Nkx2.1* and *Sst* expression in the MGE of Arx mutant mice (Figure S6A). Although the normal Sst+ cell cluster was nearly absent in male cKO MGE, many cells in the ectopic clusters express *Sst* (Figure 4D-E). Notably, the *Sst* locus had the largest increase in accessibility in male cKO mice in terms of fold change (Figures 4F & S6C). *Nkx2.1* is normally restricted to cycling MGE progenitors, but *Nkx2.1* expression is maintained in postmitotic cells in both male cKO and female Het mice (Figures 4D-E & S7).

The transcriptional changes were even more extreme in E15.5 Arx mutant mice. We extracted the entire E15.5 subpallium to ensure that all postmitotic migratory cells were included in this analysis. There were more DEGs at E15.5 compared to E13.5 in both cKO male and Het female mice (Figure S6D). Consistent with E13.5 results, markers of the ectopic cell population (*Lmo1*, *Lmo3*, *Lmo4*, *Lmx1b*, *Ebf3*) are significantly increased in Arx mutant mice (Figures S6D & S8). *Nkx2.1, Sst, Nrg1,* and *Kcnq3* were also significantly increased while *Lhx8, Maf, Ackr3,* and *Cxcr4* were decreased in E15.5 cKO mice (Figures S6D & S9A-B). Het female mice exhibited fewer DEGs compared to cKO male mice, but still displayed enrichment of *Lmo1*, *Lmo3*, *Lmo4, Sst* and reduction of *Lhx8* and *Ackr3* at E15.5 (Figure S6D). We confirmed many of these transcriptome changes via FISH (Figure S8 & S9). Notably, we observe a bolus of Nkx2.1+ MGE-derived interneurons in E13.5 and E15.5 cKO male mice stuck in the GE mantle, specifically the MZ migration stream (Figure S10A-B). These Nkx2.1+ MGE-derived cells were still confined to its periventricular zone in the ventral forebrain at P0 (Figure S10C), consistent with previous study^13^. We did observe some differences between E13.5 and E15.5. For example, *Fgf17* and *Fgf15* are expressed in the Nkx2.1+ septum at E13.5, in agreement with a previous study^49^, but both genes are strongly reduced in E15.5 cKO mice (Figure S9C).

A recent study using primary human neural progenitor cultures demonstrated that *ARX* promotes interneuron formation by repressing *LMO1*, and *ARX* knockdown in these cells produced an ectopic cell cluster containing over 700 DEGs^26^. 106 of these human DEGs were also dysregulated in our E13.5 MGE dataset. Among these, *LMO1*, *LMO3*, *LMO4* and *EBF3* were top genes upregulated in both datasets (Figure S10D), indicating strong similarities in transcriptome changes when *ARX* is removed from human and mouse neural progenitors.

In sum, integration of single nuclei Multiome-seq, spatial transcriptomics and FISH in Het female and cKO male mice revealed how reduction of *Arx* disrupts postmitotic cell fate specification, cell migration and developmental trajectories in a dosage-dependent manner via alteration in gene expression and chromatin accessibility of critical fate determining genes.

### Reduced cell activity and enhanced neuroprotective responses in the cortex of Arx mutant mice

To analyze P16-P120 spatial transcriptomics datasets, cortical regions were segmented using BANKSY^50^, which integrates a cell’s transcriptome with its local microenvironment. Cell type composition was determined using Robust Cell Type Decomposition (RCTD)^51^ in combination with the Allen Brain Institute reference scRNA-seq dataset^52^. Only singlet-assigned spots were retained for downstream analysis. Cells from P16, P21 and P42 male WT and cKO datasets largely co-clustered regardless of genotype (Figure 5A). Unsupervised clustering identified 23 hierarchically organized cell subtypes: 2 glia subtypes, 3 MGE-derived interneuron subtypes, 3 CGE-derived interneuron subtypes, and 15 excitatory neuron subtypes spanning superficial to deep cortical layers comprising intratelencephalic (IT), pyramidal tract (PT), near-projecting (NP), and corticothalamic (CT) neuron subtypes (Figures 5B-C & S11).

**Figure 5.**
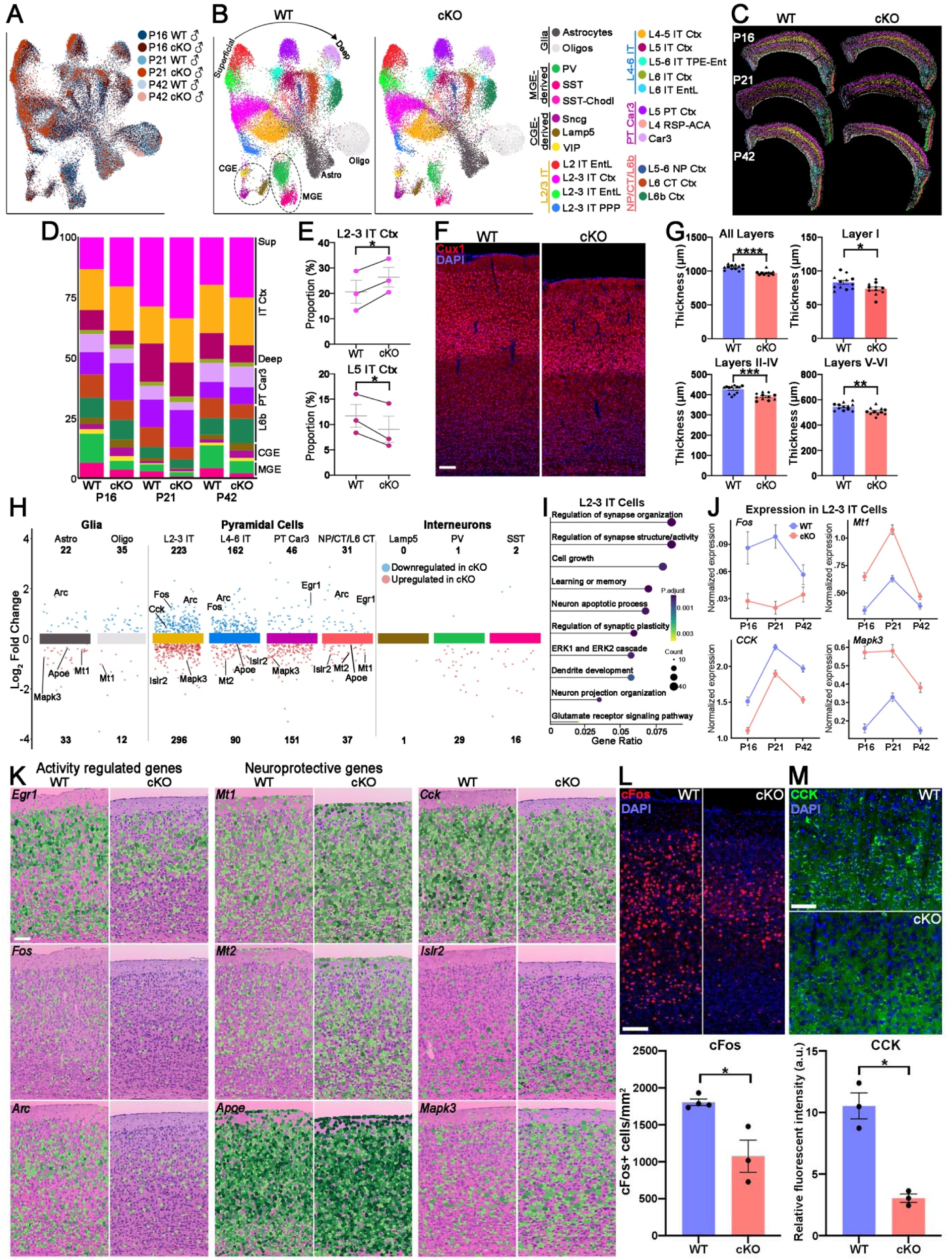
Non-cell autonomous genetic and cellular changes in the cortex of Arx cKO mice. **A-B**. UMAP plots of spatial transcripts labeled by age and genotype (A), and cell types based on gene expression (B). **C.** Spatial localization of major cortex cell types with singlet-assigned spots. **D.** Relative proportion of major cortical neuron types in WT and cKO mice. **E.** Increased proportion of L2-3 IT neurons and reduced proportion of L5 IT Ctx neurons in cKO mice. n = 3 for each genotype. **F-G.** P21 cortical sections through S1 stained with Cux1 to label layers II-IV (F), with reduced thickness across all cortical layers in cKO mice (G). Scale bar = 100 μm. n = 3 brains/genotype, 12 WT slices and 11 cKO slices counted/genotype, each brain labeled with a different shape. **H.** Scatter chart showing DEGs downregulated (blue) or upregulated (red) in cKO mice in distinct cortical cell types. **I.** clusterProfiler GO enrichment of top biological processes for DEGs of L2-3 IT cells between WT and cKO mice. **J.** Normalized expression of *Fos, Mt1, Cck* and *Mapk3* in L2-3 IT cells in WT and cKO mice at P16, P21 and P42. **K.** Spatial distribution of mRNA in P16 S1 cortex visualized in cell segmented spatial transcriptomics data. Scale bar = 100 μm. **L.** P21 S1 cortex from WT and cKO mice stained for cFos (top) and quantification of cFos+ cell density (bottom). Scale bar = 100 μm. n = 4 WT and 3 cKO mice. **M.** P21 layer 2-3 S1 cortex stained for CCK (top) and quantification of relative fluorescent intensity (bottom). Scale bar = 100 μm. n = 3 mice/genotype. Paired 2-tailed t-test (E), unpaired 2-tailed t-test (L) and Welch’s 2-tailed t-test (G, M) was performed: * = p ≤ .05, ** = p ≤ .01, *** = p ≤ .001, **** = p ≤ .0001.

We quantified the proportions of major cortical neuron types across the three postnatal time points. As expected, MGE-derived interneurons were markedly reduced in the cortex of male cKO mice at all 3 ages (Figure 5D). Removing glia from the analyses, we observed a proportional increase in cortical L2–3 IT neurons and a corresponding decrease in L5 IT neurons in male cKO mice compared to WT (Figure 5D-E). Despite this proportional difference in thickness between superficial and deep cortical layers, all cortical layers were significantly reduced in male cKO mice (Figure 5F-G).

To identify cell autonomous and non-cell autonomous DEGs in cortical cell types across postnatal development, we integrated the P16, P21 and P42 datasets and combined the 15 excitatory neuron subtypes into four broad subgroups: L2–3 IT, L4–6 IT, PT/Car3 and NP/CT/L6b (Figure 5B, H). Among these populations, L2–3 IT neurons exhibited the greatest number of DEGs (296 genes upregulated and 223 genes downregulated in cKO), which were enriched for processes including synaptic function, cell growth, neural development, and ERK signaling (Figure 5H-I).

Activity-regulated genes (ARGs) such as *Fos*, *Arc*, *Egr1* and *Nr4a1* were strongly downregulated across most excitatory neuron subgroups in male cKO mice (Figures 5H, J-K & S12). We confirmed a significant decrease in cFos protein levels in cKO males, with the strongest reduction in the most superficial and deep layers (Figure 5L). In contrast, metallothionein family genes *Mt1* and *Mt2*, which confer protection against oxidative stress and immune challenge^53^, were significantly upregulated in male cKO cortex (Figure 5H, J-K). The *Apoe* gene encodes Apolipoprotein E (ApoE), which mediates lipid transportation between cells. APOE4 variants are a major risk for Alzheimer’s disease while ApoE2 and ApoE3Ch variants confer resistance from excitotoxicity and protect neurons from ferroptosis^54^. *Apoe* expression was significantly increased in cKO cortex (Figure 5H, K). CCK+ neurons co-express neurotransmitters (GABA or glutamate), mediate synaptic transmission, and are integral components of cortical circuits^55,56^. *Cck* expression was strongly downregulated in cKO pyramidal cells, particularly in the L2–3 IT population, which we confirmed via immunostaining (Figure 5H-K, M). *Islr2*, a marker for L6a excitatory neurons that is associated with neurite extension^57^, was strongly upregulated in deep cortical layers in cKO male mice (Figure 5K). *Mapk3*, a core component of the MAPK/ERK signaling cascade that is associated with neurodevelopmental disorders in patients with 16p11.2 deletions^58^, was broadly increased in cKO cortex (Figure 5H-K).

In sum, we identified both broad and layer specific non-cell autonomous DEGs in cortical excitatory neurons in Arx cKO male mice, likely arising from the significant loss of MGE-derived interneurons in these mice. DEGs span a broad range of cellular activity including neuropeptide signaling, synaptic function, and neuroprotective responses across multiple cell types, highlighting the extent of non-cell-autonomous transcriptional consequences of interneuron loss.

### Hippocampal sclerosis and dysregulation CA3 gene network induced by BDNF in Arx cKO hippocampus

Temporal lobe epilepsy (TLE) is the most common form of focal epilepsy, and mesial temporal lobe epilepsy with hippocampal sclerosis (MTLE-HS) is a well characterized disorder, most commonly occurring in the hippocampus^59^. Since Arx mutant mice display seizures, we extracted the hippocampus from our spatial transcriptomics dataset to identify DEGs in distinct hippocampal cell populations in male cKO mice. Cells could be cleanly divided into principal cells from distinct hippocampal subdomains (CA1, CA2, CA3, DG), MGE-derived (PV and SST) and CGE-derived interneurons (Lamp5 and Sncg), and astrocytes (Figures 6A-C & S13A-B). Hippocampal oligodendrocytes were excluded from the analysis because they could not be cleanly separated from the adjacent corpus callosum in these sections. Consistent with our immunostaining results (Figure 1D-E), PV+ and SST+ interneurons were markedly reduced in the cKO hippocampus (Figure 6B). Notably, CA1 and CA3 cells were largely segregated based on genotype and age (Figure 6A-B), which is in stark contrast to the comingling of cortical cells from WT and cKO mice in a salt-and-pepper fashion (Figure 5A). This indicates greater transcriptional differences in hippocampal excitatory cells compared to the cortex.

**Figure 6.**
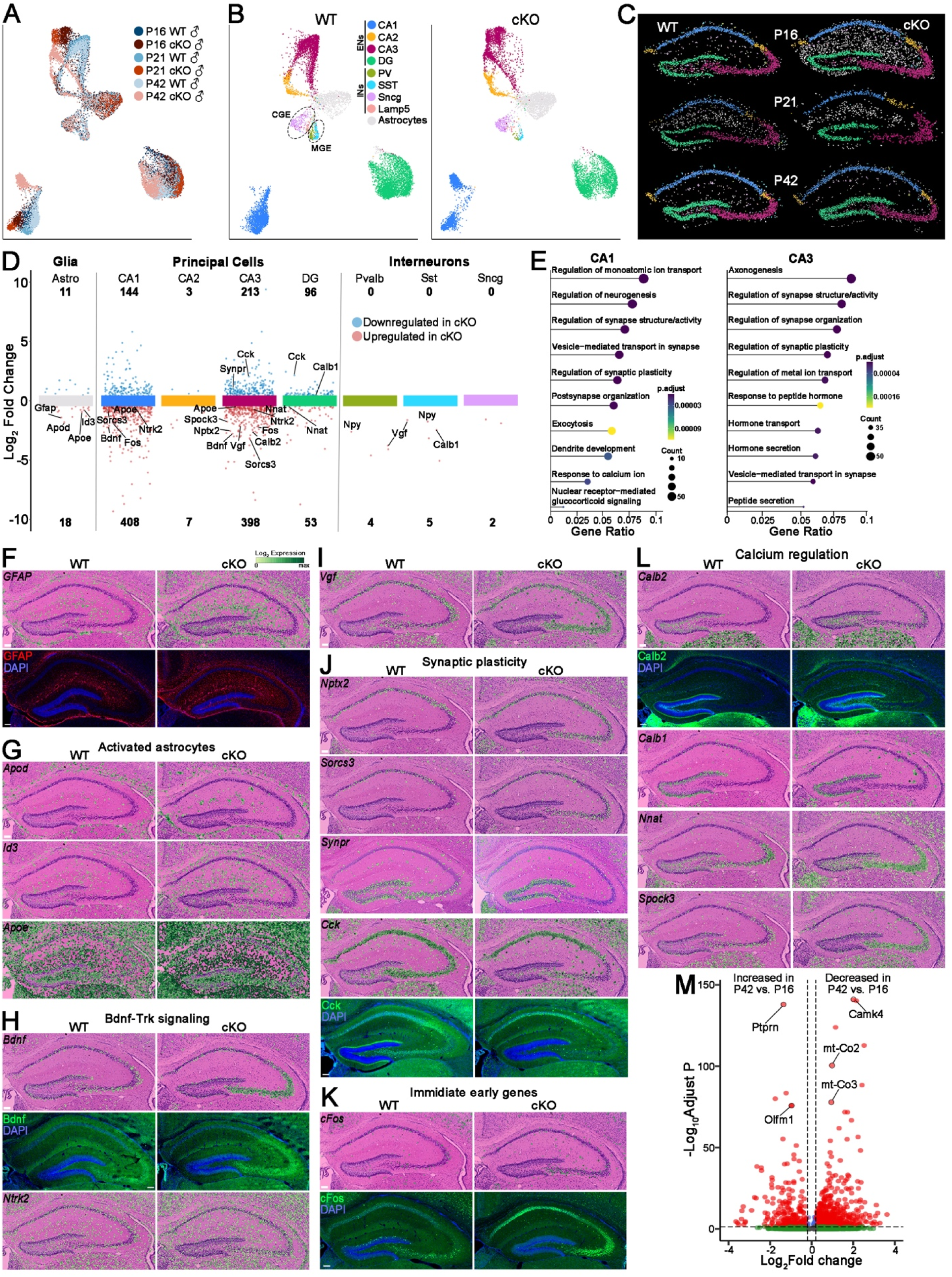
Indicators of epilepsy and genetic dysregulation in the hippocampus of Arx cKO hippocampus. **A-B**. UMAP plots of hippocampus spatial transcriptomics data labeled by age and genotype (A), and cell types (B). **C.** Spatial localization of major hippocampal cell types with singlet-assigned spots. **D.** Scatter chart showing DEGs downregulated (blue) or upregulated (red) in cKO mice in distinct hippocampus cell types **E.** clusterProfiler GO enrichment of top biological processes for DEGs of CA1 (left) and CA3 (right) between WT and cKO mice. **F-L.** Spatial distribution of mRNA in P16 hippocampus of WT and cKO mice visualized in cell segmented spatial transcriptomics data for astrocytes (F, with GFAP immunostaining), activated astrocytes (G), Bdnf-TrkB signaling (H, with BDNF immunostaining), *Vgf* (I), synaptic plasticity (J, with CCK immunostaining), immediate early genes (K, with cFos immunostaining), and calcium regulation (L, with Calb2 immunostaining). Synpr mRNA (J) was visualized in P42 hippocampus. **M.** Volcano plot depicting hippocampus DEGs between P16 and P42 cKO mice. Two-sided Wilcoxon rank sum test with Bonferroni correction (D, M) and one-sided Fisher’s exact with BH correction (E) were used. Scale bar = 100 μm.

Integrating the P16, P21 and P42 spatial transcriptomics datasets together revealed that CA1 and CA3 had the largest number of DEGs, with most DEGs being upregulated in cKO mice (Figure 6D). Many DEGs in both CA1 and CA3 were associated with synaptic structure/function, synaptic plasticity and ion transport (Figure 6E). DEGs related to hormone and peptide secretion were specifically found in CA3 whereas DEGs involved in neurogenesis and dendrite development were prominent in CA1 (Figure 6E), highlighting hippocampal subregion specificity of DEG families. Very few DEGs were identified in MGE-derived interneurons, likely due to very few SST+ and PV+ cells in these mice. Notably, *Calb1* was significantly increased in Sst+ interneurons (Figure 6D), consistent with our immunostaining results (Figure 1F). *Vgf*, which is required for activity-dependent scaling in PV+ interneurons^60^, is increased in both PV+ and SST+ interneurons (Figure 6D).

Many hippocampal DEGs are associated with epilepsy and seizures. TLE with hippocampal sclerosis has the hallmark of increased astrocyte activity^59^. There was an apparent increase in astrocytes in the P16 cKO hippocampus (Figure 6B-C), which was confirmed with increased GFAP mRNA and protein (Figure 6F), consistent of severe gliosis. Other gliosis-related genes such as *Apod*^61^ and *Id3*^62^ were significantly upregulated throughout the hippocampus of male cKO mice (Figure 6G). *Apoe* mediates lipid transfer from hyperactive neurons to reactive astrocytes in TLE^63^, and *Apoe* was upregulated in both astrocytes and principal neurons in cKO mice (Figure 6D,G). BDNF is strongly upregulated in TLE where it can paradoxically increase neuronal excitability through interaction with Trk receptors while also producing antiepileptic effects via upregulation of NPY that can suppress seizures^64,65^. In male cKO mice, *Npy* is upregulated in MGE-derived interneurons, *Bdnf* mRNA and protein are strongly upregulated in the CA3 of male cKO mice, and its receptor *Ntrak2* is also upregulated throughout the hippocampus (Figures 6D,H & S14A). BDNF stimulates Vgf expression to modulate hippocampal synaptic transmission^66^, and *Vgf* is also upregulated in CA3 pyramidal cells (Figures 6D,I & S14A).

We observed dysregulation of numerous genes involved in synaptic transmission and plasticity. Neuronal pentraxin 2 (*Nptx2*) and Sortilin-related VPS10 domain-containing receptor 3 (*Sorcs3*) are postsynaptic proteins normally restricted to CA1 where they play critical roles in synaptic plasticity^67,68^. Both *Nptx2* and *Sorcs3* are significantly upregulated in CA3 of male cKO mice (Figures 6D,J & S14A). Synaptoporin (*Synpr*) is a synaptic vesicle protein expressed in DG mossy cells and CA3 cells where it modulates homeostatic plasticity^69,70^, but *Synpr* expression was lost in the CA3 of cKO mice (Figure 6D,J). *Cck* is normally expressed throughout CA1-CA3 pyramidal cells where it regulates excitatory cell output^71^, but it is significantly downregulated CA3 of male cKO mice (Figures 6D,J & S14A). The dysregulation of genes associated with synaptic plasticity in the hippocampus likely reflect a hyperexcitable state in cKO mice, confirmed by increased cFos mRNA & protein in CA1 and CA3 (Figures 6D,K & S14A).

Hippocampal calcium plays a critical role in the generation and establishment of epilepsy. Calbindin (*Calb1*) and Calretinin (*Calb2*), two seizure sensitive calcium-binding proteins, are often dysregulated in epilepsy^72,73^. *Calb2* mRNA and calretinin protein are upregulated in the DG and aberrantly expressed in CA3 in cKO hippocampus, while *Calb1* mRNA and calbindin protein are downregulated in the DG (Figures 6D,L & 1F). Neuronatin (*Nnat*) can modify dendritic calcium levels in hippocampal neurons^74^, and *Nnat* was increased in the cKO DG and CA3 of hippocampus (Figure 6D,L & S14A). Testican-3 (*Spock3*), a calcium-binding proteoglycan associated in the extracellular matrix^75^, was significantly upregulated in cKO CA3 (Figures 6D,L & S14A).

In addition to common DEGs throughout all 3 ages, we also identified age-specific DEGs between P16 and P42. Genes associated with seizure susceptibility (*Ptprn*)^76^, spine and axon growth (*Camk4, Olfm1*)^77,78^ and mitochondrial organization (*mt-Co2, mt-Co3*)^79^ were significantly changed in the hippocampus between P16 and P42 cKO male mice (Figure 6M & S14B), suggesting dynamic changes in gene expression likely due to seizure progression and altered brain function in Arx cKO male mice over time. These data demonstrate that loss of MGE-derived hippocampal interneurons induce both broad and region-specific transcriptome changes in hippocampal principal neurons related to seizures and hippocampal sclerosis, and altered neuronal and synaptic function.

### Dysregulation of energy metabolism and initiation of transcription and translation in Arx Het cortex and hippocampus

Since Arx Het female mice & humans also display phenotypes, we analyzed cell autonomous and non-cell autonomous transcriptional changes in juvenile and adult Arx Het females. Single nuclei Multiome sequencing was performed on GFP+ Nkx2.1-lineage cortical interneurons from one WT and two Het P70 female mice (Figure 7A). MGE-derived interneurons were cleanly segregated into SST+ and PV+ interneurons, which could be further divided into 8 PV+ and 10 SST+ subtypes^30,80^ (Figure 7B). There were no significant differences in the cell clusters or proportional distribution of PV+ and SST+ interneuron subtypes between WT and Het female mice (Figure 7B). A small number of DEGs were identified in both PV+ and SST+ interneurons (Figure 7C), likely because surviving interneurons express WT Arx with the mutant Arx allele undergoing X-inactivation. *Stmn3*, which is involved in microtubule formation and dynamics^81^, was upregulated in both PV+ and SST+ cells in Het mice (Figure 7C). Dysregulation of Stmn3 protein was also reported in a different Arx mutant mouse line^82^, consistent with our findings.

**Figure 7.**
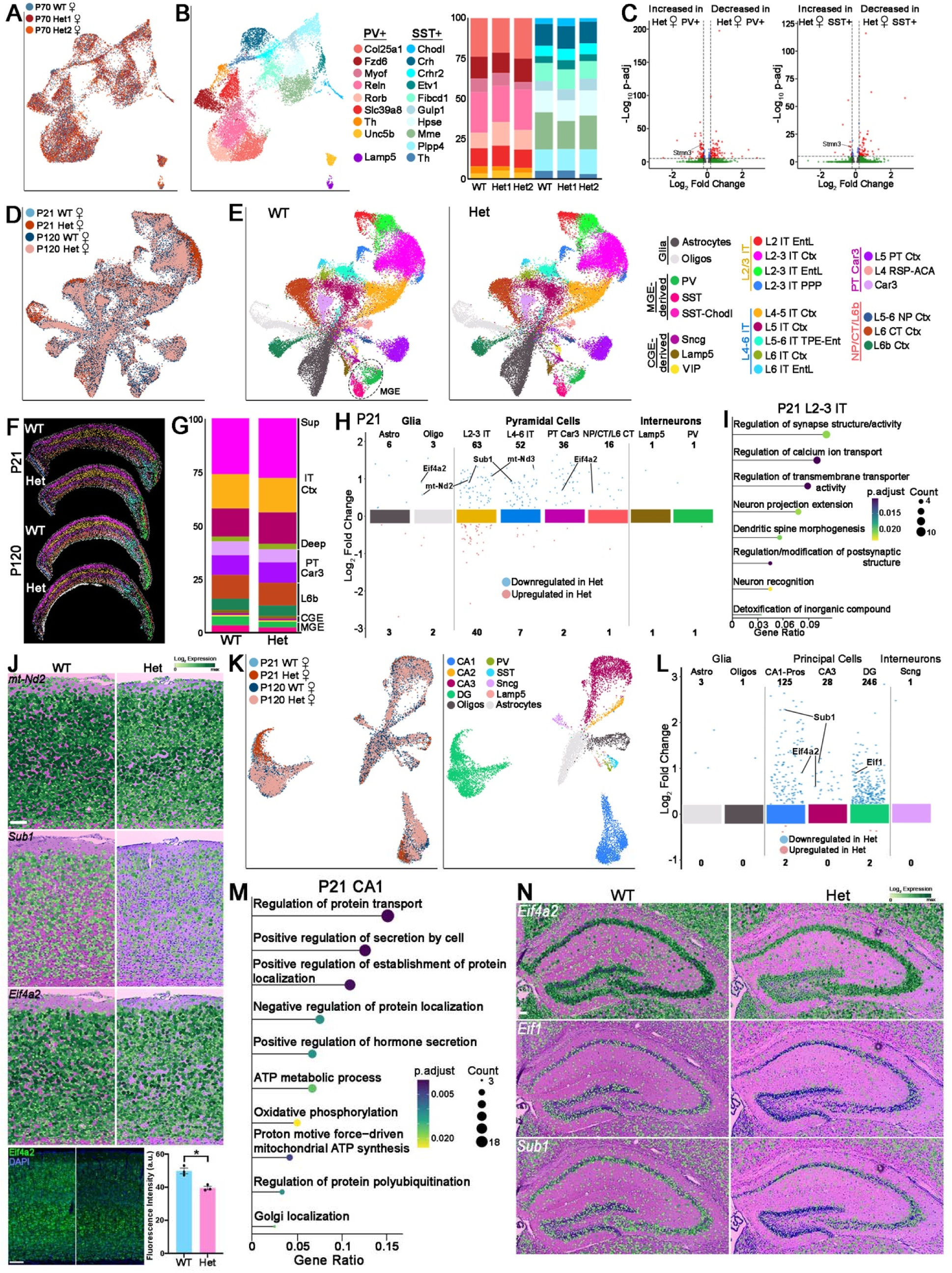
Downregulation of genes associated with energy metabolism, transcription and translation in Arx Het cortex and hippocampus. **A-B**. UMAP plots of integrated snRNA and snATAC data via weighted nearest neighbor (WNN) of Nkx2.1-lineage cells from P70 female WT and Het cortex, annotated by genotype (A), and PV+ and SST+ interneuron subtypes with relative proportions (B). **C.** Volcano plot depicting DEGs in P70 female Het PV+ (left) and SST+ (right) interneurons. **D-E.** UMAP plots of spatial transcripts labeled by age and genotype (D), and major cortical cell types (E). **F.** Spatial localization of major cortical cell types with singlet-assigned spots. **G.** Relative proportion of major cortical neuron types of female WT and Het mice. **H.** Scatter chart showing DEGs downregulated (blue) or upregulated (red) in distinct cortical cell types in P21 Het mice. **I.** clusterProfiler GO enrichment of top biological processes for DEGs of L2-3 IT cells between WT and Het mice. **J.** Spatial distribution of mRNA in P21 cortex of WT and Het mice visualized in cell segmented spatial transcriptomics data for *mt-Nd2*, *Sub1* and *Eif4a2* (top), with Eif4a2 immunostaining and quantification (bottom). n = 3 for each genotype. **K.** UMAP plots of P21 and P120 hippocampal spatial transcriptomics labeled by age and genotype (left) and cell types (right). **L.** Scatter chart showing DEGs downregulated (blue) or upregulated (red) in Het mice in distinct hippocampus cell types in P21 mice. **M.** clusterProfiler GO enrichment of top biological processes for DEGs of P21 CA1. **N.** Spatial distribution of mRNA in P21 hippocampus of WT and Het mice visualized in cell segmented spatial transcriptomics data. Two-sided Wilcoxon rank sum test with Bonferroni correction (C, H, L), one-sided Fisher’s exact with BH correction (I, M) and Welch’s 2-tailed t-test (J) were used. Scale bar = 100 μm.

We then investigated non-cell autonomous gene expression changes in P21 and P120 WT and female Het mice via spatial transcriptomics. Similar to cKO male cortices, cells from female P21 and P120 WT and Het mice co-clustered regardless of genotype (Figure 7D). We identified the same 23 cortical cell types, with no obvious changes in the proportion of any subtypes except for the expected moderate reduction of MGE-derived interneurons (Figures 7E-G & S15A). No significant difference in cortical thickness was found between WT and Het mice (Figure S15B). There were many more DEGs at P21 compared to P120, with the majority of DEGs downregulated in Het pyramidal cells at P21 whereas most DEGs were upregulated in Het pyramidal cells and astrocytes at P120 (Figures 7H & S15C). These DEGs were enriched for processes related to synapse structure or activity, calcium ion transport, neuron projection extension and dendritic spine morphogenesis (Figure 7I). Notably, mitochondrial genes (*mt-Nd2, mt-Nd3*) were broadly downregulated across cortical cell types at P21 (Figures 7H,J & S15A), indicating impaired energy metabolism in Het female mice. *Sub1*, a component of the transcriptional regulatory complex, was similarly decreased throughout the cortex of Het female mice (Figures 7H,J & S15A). *Eif4a2*, an ATP dependent RNA helicase whose dysfunction is associated with neurodevelopmental disorders featuring ID and seizures^83^, was markedly reduced in Het cortex, with Eif4a2 protein most strongly decreased in superficial cortical layers (Figures 7H,J & S15A). *Eif4a2* was also dysregulated in mouse model of polyalanine repeats at the *Arx* locus where protein translation is suppressed^82^.

As in the cortex, WT and Het hippocampal cell types showed substantial overlap between genotypes, although there was some age-based clustering in CA1 and DG principal cells (Figure 7K). CA1 and DG regions exhibited the greatest number of DEGs at both P21 and P120, with nearly all DEGs downregulated in Het females (Figures 7L & S15D). In P21 CA1, DEGs were enriched for protein transport and localization, ATP metabolism, and hormone secretion, whereas in the DG, DEGs were associated with transcription and translation, RNA splicing and lamellipodium organization (Figures 7M & S15E). Similar to the cortex, *Eif4a2*, *Eif1* and *Sub1* were downregulated in the hippocampus of Het female mice (Figures 7L,N & S15A). We did not observe significant changes in epilepsy-associated genes (*BDNF*, *Vgf*, *Cck*, etc.) as we did in the hippocampus of male cKO mice.

Overall, as expected there were many fewer DEGs throughout the cortex and hippocampus of Arx Het female mice compared to cKO males. Many DEGs were associated with fundamental transcriptional and translational processes, as well as mitochondrial organization and ATP synthesis, across both cortex and hippocampus.

### Loss of MGE-derived interneurons alters dorsal cortical neurogenesis, cellular organization and induces band heterotopias

Dorsal cortical progenitors progress through temporal specification guided by both intrinsic genetic programs and extrinsic cues^84,85^. Interneurons migrating through the cortex can influence pyramidal cell neurogenesis, radial migration and cortical lamination^86–88^. The decreased cortical thickness combined with proportional changes in the number of L2-3 IT and L5 IT pyramidal neurons in cKO male mice (Figure 5D-G) suggest that loss of migrating MGE-derived interneurons could influence on dorsal progenitor neurogenesis and cell fate.

We defined nine cell types from the spatial transcriptomics data in E13.5 and E15.5 Arx cKO and WT male mice, including progenitors (radial glial cells (RGCs), intermediate progenitor cells (IPCs)), postmitotic excitatory neurons (immature deep layer cortical projection neurons (Immature DL CPNs), deep layer cortical projection neurons (DL CPNs), upper layer cortical projection neurons (UL CPNs) and migratory excitatory neurons (Migratory ExNs)) and two streams of migrating interneurons (INs_SVZ stream and INs_MZ stream) (Figures 8A & Fig. S16A-B). The vast majority of DEGs were upregulated in cortical excitatory and inhibitory cells of E13.5 cKO male mice but were downregulated at E15.5 (Figures 8B & S16C). As expected, many markers of postmitotic interneurons (*Sst*, *Maf*, *Lhx6, Nxph1,* etc.) were significantly reduced in the cortex of E13.5 and/or E15.5 cKO male mice, while *Nkx2.1* and *Ebf3* were increased in cortical interneurons in cKO mice (Figure 8B-C & S16C).

**Figure 8.**
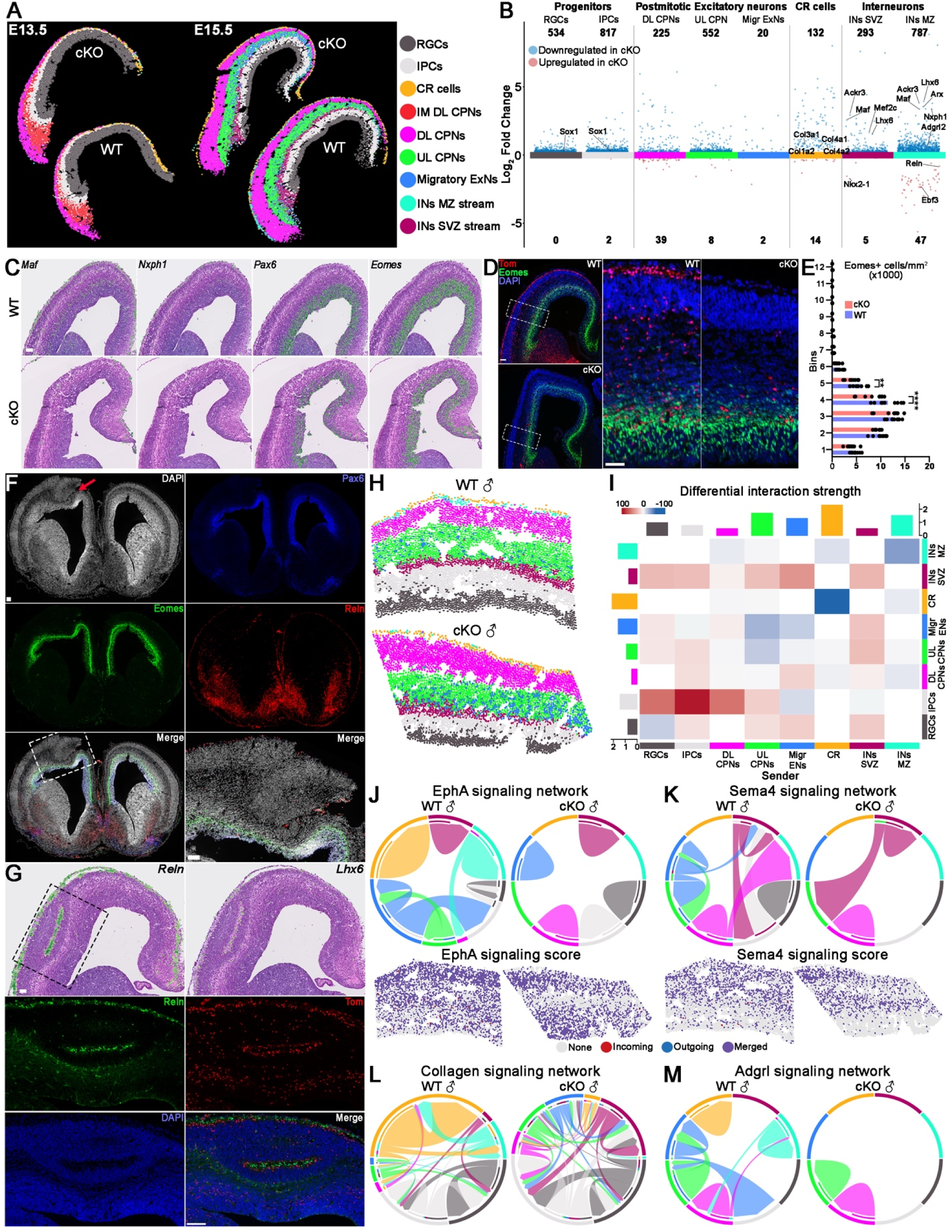
Loss of MGE-derived interneurons alters dorsal cortical neurogenesis and cell-cell signaling leading to cortical heterotopias. **A.** Spatial localization of E13.5 and E15.5 WT and cKO cell types from cell segmented spatial transcriptomics data. **B.** Scatter chart showing DEGs downregulated (blue) or upregulated (red) in cKO mice in distinct E15.5 cell types. **C.** Spatial distribution of mRNA in E15.5 cortex of WT and cKO mice visualized in cell segmented spatial transcriptomics data. **D-E:** Immunostaining (D) and quantification (E) of Eomes+ IPCs in E15.5 lateral cortex of WT and cKO mice. n = 3 brains/genotype, n = 9 slices/genotype. **F.** E15.5 cKO brain section stained for Pax6, Eomes and Reln and DAPI highlighting cortical layering abnormality. **G.** Spatial distribution of *Reln* and *Lhx6* mRNA in E15.5 cortex of WT and Het female mice visualized in cell segmented spatial transcriptomics data (top), and immunostained for Reln and tdTomato (bottom). **H.** Spatial localization of cortical cell types and area used for CellChat analysis from E15.5 WT and cKO male mice, color coding same as A. **I.** CellChat-derived heatmap depicting differential interaction strength between cell types from inferred cell-cell communications in E15.5 WT and cKO cortex. **J-M.** Chord diagram of EphA (J), Sema4 (K), collagen (L), and Adgrl (M) signaling networks between different cell types from inferred communication at the individual cell level in WT and cKO cortices, with spatial visualization of signaling score for EphA (J) and Sema4 (K). Two-sided Wilcoxon rank sum test with Bonferroni correction (B) and ordinary two-way ANOVA followed by Sidak’s multiple comparisons test (D) were used: ** = p ≤ .01, **** = p ≤ .0001. Scale bar = 100 μm.

At E13.5, several postmitotic excitatory neurons markers (*Neurod2, Neurod6, Bcl11b*, *Tbr1*) were significantly increased in cKO cortex (Figure S16C), suggesting premature cell cycle exit. Consistent with this, genes associated with cycling progenitor cells and neurogenesis (*Pax6*, *Eomes*, *Mki67, Sox1*) were significantly downregulated in the E15.5 cortex of cKO male mice (Figures 8B & S16C), and we confirmed a significant decrease of Eomes+ cells in the E15.5 cKO cortex (Figure 8D-E). Thus, loss of migratory MGE-derived interneurons in the dorsal cortex alters neurogenesis of excitatory projection neurons.

Patients with *ARX* variants can display lissencephaly and other cortical malformations^5,89,90^. We observed abnormal cortical structures resembling band heterotopias in both cKO male (Figure 8F) and het female mice (Figure 8G), and instances of oddly shaped pointed dorsal cortices in other brains (Figure S16D). This abnormal cortical formation appears to compress and disrupt the normal organization of Pax6+ and Eomes+ dorsal cortical progenitors (Figure 8F). Our spatial transcriptomics data revealed an ectopic layer of Reln+ Cajal-Retzius (CR) cells within this band heterotopia that is surrounded by Tom+ MGE-derived interneurons (Figure 8G). CR cells are required for normal inside-outside cortical patterning, and Reln is also expressed by interneurons where it plays a role in layer-specific distribution of cortical neurons^87^. Notably, *Reln* is upregulated in the IN_MZ stream in cKO mice (Figure 8B). These results demonstrate that loss of MGE-derived interneurons is sufficient to disrupt normal CR cell organization, likely leading to severe perturbation of dorsal cortical neurogenesis.

The non-cell-autonomous effects of *Arx* deletion in MGE-derived interneurons is likely due to dysregulated cell-cell communications. We inferred and analyzed cortical cell–cell communication networks from E15 spatial transcriptomic data using CellChat^91^. We performed CellChat analysis on ∼5000 dorsal cortical cells in E15.5 WT and cKO mice (Fig. 8H). The total number of inferred cell-cell communication networks were increased in cKO cortex, but the interaction strength was decreased (Figures 8I & S16E-F). There was a strong increase in both the number and strength of interactions between progenitor cells (RGCs and IPCs) with nearly all other cell types (Figures 8I & S16F). Conversely, interneurons in the MZ and CR cells had a strong reduction in interaction numbers and strength, with the strongest observed decrease in CR-CR communication (Figures 8I & S16F). This is consistent with our immunostaining results showing significant changes in these progenitor and CR cells.

The most altered signaling network in cKO mice involved two guidance factor families (EphA receptors, Sema4 ligands) and two families involved in cell adhesion (collagen and members of the adhesion GPCR L subfamily, ADGRL) (Figure 8J-M). EphA receptors and their ephrin ligands play critical role in neural proliferation, cell migration, synaptogenesis, and topographical patterning of the cortex^92^. EphA-based interactions between UL CPNs, CR and INs_SVZ stream were lost in cKO mice (Figure 8J). Class 4 semaphorins mediate receptor clustering in neuronal postsynaptic membranes, and deficiencies in Sema4 signaling is associated with defects in cell proliferation, exencephaly and neonatal lethality^93^. Sema4 interactions were completely lost between most cell types, but there is a predicted new sema4-based signaling network between UL CPNs and INs_SVZ cells in cKO mice (Figure 8K).

Collagen signaling is involved in guiding neuronal outgrowth, maturation and synaptogenesis^94^, and ADGRL signaling controls cell–cell and cell–matrix interactions during neural development^95^. Mutations in the adhesion GPCR gene *Adgrg1* (GPR56) and its ligand collagen type III alpha-1 (*Col3a1*) results in a malformed cerebral cortex that resembles cobblestone lissencephaly^96,97^, similar to what we see in Arx mutant mice. The collagen signaling network predominantly involves RGCs, IPCs and CR cells in WT mice, but collagen signaling is strongly downregulated in CR cells and upregulated in postmitotic cortical neurons in cKO mice (Figure 8L). This is consistent with numerous collagen genes (*Col3a1, Col4a1, Col4a2, Col1a2*) being downregulated specifically in CR cells in cKO mice (Figure 8B). ADGRL signaling was lost in all cell types except for CPNs in cKO mice (Figure 8M). Thus, reduction of MGE-derived interneurons in Arx mutant mice disrupts numerous signaling families causing altered cell-cell communication and cellular disorganization of CR cells and dorsal cortical progenitors, leading to profound defects in cortical neurogenesis.

## DISCUSSION

*ARX* variants are a leading cause of XLID, a complex set of disorders with a high comorbidity of ID and epilepsy that predominantly affects males, but female carriers also display a variety of phenotypes. In this study, we demonstrate how removal of *Arx* in the MGE leads to a near complete loss of forebrain MGE-derived interneurons and early lethality in cKO male mice, and reduced interneurons with increased seizure susceptibility and altered behavior in Het female mice. We constructed a transcriptomic atlas spanning embryogenesis through adulthood using single nuclei sequencing and spatial transcriptomics. These data revealed significant transcriptomic changes in the MGE leading to an *Arx* mutant-specific cell cluster that is consistent with a recent study in human forebrain progenitor primary cultures^26^. Most notably, the loss/reduction of MGE-derived interneurons in the embryonic cortex had a profound non-cell autonomous effect on gene expression, neurogenesis, cellular organization and cell-cell communication in dorsal cortical progenitors, ultimately leading to cortical abnormalities mimicking band heterotopias in both cKO male and Het female mice. These findings highlight that loss of *Arx* in the MGE is sufficient to induce both cell autonomous and non-cell autonomous transcriptomic changes that may explain the variation of symptoms and severity in patients *ARX* variants and XLID phenotypes.

Reduction of cortical MGE-derived interneurons was consistent across subtypes and closely followed Arx dosage, with a near complete loss of PV+ and SST+ cells in cKO male mice and ∼50% reduction in Het females, although there was some variability in subtype loss between different hippocampal regions. cKO male mice displayed infantile spasms and seizures, similar to removing *Arx* in all ganglionic eminences^17^, which likely led to most cKO males dying before weaning. Most Het female mice did not exhibit spontaneous seizure, and while Het females did show a significant increase in seizure susceptibility overall, ∼50% of Het mice had similar response to WT females. Het female mice also displayed significant but variable deficits in locomotor activity, anxiety behaviors and sensorimotor gating. These results are consistent with human studies where female carriers have a large range of mild to severe phenotypes^9^, even in monozygotic female twins with *ARX* variants^89^, implying mosaic XCI could in part lead to different phenotype severity. This study did not correlate XCI with behavior or cellular phenotypes, but Het female mice have a normal paternal X-chromosome and a mutant maternal X-chromosome based on our mating strategy.

Reduced MGE-derived interneurons are consistent with our embryonic transcriptome data where we found a complete loss of normal *Sst*+ and PV-fated *Maf*+ cell clusters in cKO male mice and an ∼50% reduction in Het female mice. These clusters were replaced with ectopic postmitotic cell clusters in cKO mice enriched for Lim homeodomain transcription factors (*Lmo1, Lmo3, Lmo4*) and dysregulation of genes associated cell migration (*Cxcr4*, *Cxcr7*, *Nrg1*); this population likely represents the bolus of cells whose dorsal migration was halted in the ventral telencephalon. *Sst* was also enriched in these ectopic clusters, though its role in these cells is unclear. *Lhx6* expression was maintained in these ectopic postmitotic cell clusters whereas *Lhx8*, which promotes forebrain cholinergic neuron fates^34^, was strongly downregulated. Lhx6 directly regulates Arx in the MGE, and Arx expression can partially rescue interneuron maturation in *Lhx6^−/−^* mice^98^, indicating Lhx6 acts upstream of Arx. In *Lhx6^−/−^*;*Lhx8^−/−^*double KO mice, *Lmo3* and *Nkx2.1* are both downregulated^35^, which is the opposite effect in Arx mutants. Shh expression in the MGE mantle is reduced in these double KO mice leading to abnormal progenitor differentiation^35^; Shh was also downregulated in Arx cKO mice. These findings highlight the intricate relationship between Arx, Nkx2.1, Lhx6, Lhx8 and Shh in regulating fate & maturation of MGE-derived neurons.

Loss of MGE-derived interneurons in *Arx* mutant mice led to striking non-cell autonomous cellular and transcriptional changes in the cortex and hippocampus. Neuroprotective genes were upregulated whereas immediate early genes (*Arc*, *cFos*, *Egr1*) were downregulated in pyramidal cells across all cortical layers in cKO male mice despite their increased seizure activity. This agrees with studies showing chronic network hyperactivity leads to long-term reduced expression of activity-dependent genes^99–102^. Many DEGs were associated dendrite development, cell growth and synaptogenesis, which is consistent with sustained hyperactivity leading to more immature neuronal states that may be a shared phenotype by many neuropsychiatric disorders^103,104^.

The hippocampus of cKO male mice displayed gliosis and other phenotypes indicative of TLE. Bdnf-Trk signaling is often upregulated in TLE^64^, especially in CA3 where BDNF may act as a central hub for network processing^65^. Bdnf mRNA and protein are both strongly upregulated in the CA3 of cKO mice. In general, the most notable changes in gene expression levels were observed in CA3 compared to other regions. Different cohorts of DEGs were identified between CA1 and CA3, with many CA3 DEGs associated with peptide and hormonal function that were not found in CA1 DEGs. Curiously, cfos protein was strongly upregulated in CA1 and CA3 pyramidal cells in the hippocampus, in contrast to the clear reduction of cfos in the cortex. Thus, loss of forebrain MGE-derived interneurons produces distinct genetic and cellular effects between the cortex and hippocampus, and between different regions of the hippocampus.

As expected, juvenile and adult Het female mice displayed fewer transcriptomic changes in interneurons and excitatory cells compared to cKO males. There was a strong bias for DEGs to be downregulated in cortical and hippocampal excitatory cells in Het female mice, while the distribution was more even or biased toward upregulation in cKO males. This could be due to decreased expression of transcription-associated (*Sub1*) and translation-associated (*Eif4a2*, *Eif*) in both cortical and hippocampal excitatory cells. In contrast to our data, *Eif4a2* was significantly upregulated in the cortex (but not hippocampus) in male Arx^(GCG)7/Y^ mice, a model of the elongated polyalanine *ARX* variant^82^. This highlights the need to explore transcriptional changes in different cell types and different Arx disease models to fully understand the broad range of phenotypes observed in both males and females.

Lissencephaly is a spectrum of severe brain malformations where normal brain folding fails to occur properly, with *ARX* variants being one of the leading genetic causes of this disease^90,105^. A related phenotype called band heterotopias (or double cortex) occurs when there is an ectopic band of neurons within or below the normal cortex, most commonly arising in females with *DCX* or *LIS1* variants^106^. We observe cortical malformations that resemble neocortical heterotopia and subcortical band heterotopia in both cKO male and Het female brains. In both cases we find an ectopic band of Reln+ CR cells, and in a E15.5 Het female brain, these CR cells are encircled by Tom+ MGE-derived interneurons. CellChat analysis predicts a strong downregulation of interaction strength for CR-CR and CR-IN_MZ communications, of which the collagen signaling network may play a prominent role. Thus, decreasing Arx expression in the MGE, and the corresponding loss of MGE-derived interneurons migrating into the cortex, is sufficient to induce cellular rearrangements, abnormal cell-cell signaling, reduced IPCs and ultimately cortical malformations. These findings demonstrate the critical role that migratory MGE-derived interneurons play in regulating dorsal cortical neurogenesis and highlight potential disease mechanisms in *ARX*-related neurodevelopmental disorders.

## METHODS

### Animals

All experimental procedures were conducted in accordance with the National Institutes of Health guidelines and were approved by the NICHD Animal Care and Use Committee (protocol #23-047 & 26-047). The following mouse lines were used in this study: *Arx-floxed* (gift from Drs. Eric Marsh & Jeffrey Golden)^38^; *Nkx2.1-Cre* (Jax# 008661)^27^, *Ai9* (Jax# 007909)^107^, *Sun1-sfGFP* (Jax# 030952)^108^. Mice were housed under standard conditions (12h light and 12h dark). The morning on which the vaginal plug was observed was denoted E0.5.

### Brain harvesting

Embryonic brain dissections: Pregnant dams were anesthetized with an i.p. injection of Euthasol (270 mg/kg, 50 μL injection per 30 g mouse), E13.5 embryos were removed and placed in ice-cold carbogenated artificial cerebral spinal fluid (ACSF, in mM: 87 NaCl, 26 NaHCO_3_, 2.5 KCl, 1.25 NaH_2_PO_4_, 0.5 CaCl_2_, 7 MgCl_2_, 10 glucose, 75 sucrose, saturated with 95% O_2_, 5% CO_2_, pH 7.4). MGE was dissected from individual brains in ice-cold ACSF, transferred to individual 0.5 mL Eppendorf tubes, then flash frozen on dry ice and stored at –80°C. For analyzing fixed tissue, E13.5 and E15.5 brains were removed and drop-fixed in 4% paraformaldehyde (PFA) in PBS overnight, washed in PBS, incubated in 30% sucrose in PBS at 4°C overnight, embedded in OCT and stored at –80°C.

Postnatal and adult brain dissections: P0-P120 mice were anesthetized with Euthasol. For harvesting brain regions, the somatosensory and motor cortex of P70 mice were dissected, transferred to 1.5 mL Eppendorf tubes, then flash frozen on dry ice and stored at –80°C. For analyzing fixed brains, P0-P120 mice were perfused with PBS followed by 4% PFA, brains were removed and post-fixed in 4% PFA overnight, washed in PBS, incubated in 30% sucrose in PBS at 4°C overnight, then embedded in OCT and stored at –80°C.

### Tissue staining and microscopy

E13.5 embryonic brains were cryosectioned at 14μm, mounted on slides, dried and stored at –20°C or –80°C. Slides were incubated in block solution (10% Normal Donkey Serum in PBS + 0.3% Triton X-100) for 1 hour RT, then incubated overnight with primary antibodies in block solution at 4°C and secondary antibodies for 2 hours RT. After washing, brain sections were mounted.

P0-P120 brains were cryosectioned at 30 μm, sections transferred to 96-well plates containing antifreeze solution (30% ethylene glycol, 30% glycerol, 40% PBS) and stored at –20°C. Prior to staining, brain sections were washed in PBS (3 x 10 minutes). Brain sections were incubated in block for 1 hour RT, then incubated for 24-48 hour in primary antibodies in block solution at 4°C. Then sections were washed in PBS (3 x 10 minutes) and incubated with secondary antibodies and DAPI for 2 hours at RT or overnight at 4°C. Brain sections were washed and mounted.

The following primary antibodies were used in this study: rat-anti SST (1:300, Millipore MAB354), goat-anti PV (1:1000, Swant PVG213), rabbit-anti nNos (1:500, Millipore MAB5380), rat-anti EOMES (1:500, Invitrogen 14-4875-82), rabbit-anti CUX1 (1:500, Proteintech 11733-1-AP), rabbit-anti c-Fos (1:500, Synaptic Systems 226008), rabbit-anti Pax6 (1:500, BioLegend 901301), mouse-anti Reelin (1:500, Sigma-Aldrich MAB5364), rabbit-anti BDNF (1:500, Invitrogen PA5-85730), rabbit-anti proCCK (1:1000, Frontier Institute MSFR105040), rabbit-anti Calb1 (1:200, Invitrogen 711443), mouse-anti Calb2 (Calretinin) (1:1000, Millipore MAB1568), mouse-anti GFAP (1:500, Sigma-Aldrich G3893), rabbit-anti Olig2 (1:500, Sigma-Aldrich AB9610), rabbit-anti eIF4A2 (1:500, Invitrogen PA5-79195), mouse-anti SYT2 (1:1000, Abcam ab154035), chick-anti VGAT (1:500, Synaptic Systems 131006), sheep-anti ARX (1:100, R&D systems AF7068). Species-specific fluorescent secondary antibodies (1:500) were used conjugated to AlexaFluor 488, 594, 647 and 790.

All images were captured on an Olympus VS200 scanner (VS200 ASW). Post-processing was performed with Adobe Photoshop and/or ImageJ2 (v2.9.0).

### HiPlex12 RNAscope

Brains were cryosectioned at 14 μm. RNAscope HiPlex12 reagents kit (Advanced Cell Diagnostics, #324409) were used according to the manufacturer’s instructions. All RNAscope HiPlex probes and reagents from Advanced Cell Diagnostics. The following probe sets were used in this study: Lmo1 (511211-T1), Lmo3 (497631-T2), Lmo4 (485491-T3), Lhx6 (422791-T4), Lhx8 (515101-T5), Ebf3 (576871-T6), Lmx1b (412931-T7), Nkx2-1 (434721-T8), Sst (404631-T9), Maf (412951-T10), Shh (314361-T11) and Nrg1 (418181-T12).

### Cell counting and analysis

All cell counts on Nkx2.1-Cre;Arx;Ai9 mice were performed manually using Photoshop and were blind to genotypes. Cortex: Cells were counted from 3-4 non-consecutive brain sections through the somatosensory cortex for each P21 brain. DAPI staining was used to divide the cortex into superficial (I-III) and deep (IV-VI) layers. Tom+, PV+ and SST+ cells were counted, and the density was calculated by cells/mm^2^. Any PV+ and SST+ cells that were Tom-were excluded from the counts. Hippocampus: Cells were counted from 4-6 non-consecutive sections spanning the anterior-to-middle P21 dorsal hippocampus. Tom+, PV+, SST+ and nNos+ cells counted, and the density was calculated by cell numbers/mm^2^. Small Tom+ cell bodies in CA2/3 that are primarily Olig2+ oligodendrocytes were excluded from these interneuron counts and counted as separate group. The number of Tom+/Calb1+ or Tom+/Calb1+/SST+ were counted from 4 sections through whole hippocampus, and 3 male WT and cKO brain were used. The thickness of the primary somatosensory cortex (S1) from P21 male mice (bregma: from 0.97 to 0.49 mm) and female mice (bregma: from –1.55 to –1.91 mm) were measured. Cux1 immunostaining was used to label layers II-IV and 3 mice per genotype, 3-4 sections measured per brain. cFos+ cells were counted from 2-4 sections through P21 the somatosensory cortex of male WT and cKO mice. The relative fluorescent intensity of CCK in layer 2-3 of somatosensory cortex was measured in three P21 male WT and cKO mice with 4 sections per mice. The E15.5 lateral cortex of male WT and cKO was divided into 12 bin and the quantification of EOMES+ cells were calculated from 3 sections for each brain.

### Pentylenetetrazole (PTZ) injection and recording

PTZ (Sigma-Aldrich, P6500) was prepared on the day of use with in sterile 0.9% (w/v) NaCl. PTZ injections and recording were performed on 4-month-old female mice. All mice receive an IP injection of PTZ at 35 mg/kg and were recorded immediately via video tracking system ANY-maze (Stoelting Company) for 60 min. Epileptic behaviors were scored using a modified Racine scoring system^109^: stage 0 = normal behavior, stage 1 = immobility and rigidity; stage 2 = head bobbing, facial, forelimb, or hindlimb myoclonus; stage 3 = continuous whole-body myoclonus, myoclonic jerks or tail held up stiffly; stage 4 = tonic seizure, continuous rearing and falling, stage 5 = clonic-tonic seizure, stage 6 = death. Total seizure scores were calculated at every 5-min block in the first 20 min. N (WT) = 9 and N (Het) = 11.

### Behavioral tests

All behavioral tests were performed on WT, Het and Hom littermates according to approved NICHD and/or NIMH standard operating procedures (SOPs) and cleaning protocols. Animal behavioral tests were carried out during the daylight cycle in NICHD animal behavioral room or at the NIMH Rodent Behavior Core. All mice were habituated to the testing environment for 30-60 minutes before behavior tests. All behavior assays were performed blind to genotype, and 1-4 independent cohorts were used for each behavioral test.

### Home-cage test

Individually housed juvenile mice were observed in their home cages placed in the Photobeam Activity System-Home Cage (PAS-HC, SD Instruments) to assess activity (ambulation, fine movements and rearing) that can be quantified by beam breaks. Mice were kept in the cage for 4 days with appropriate food and water. During this testing, mice were housed in a ‘quiet’ room to minimize outside interactions. Mice were returned to group housing after experiment. The total ambulatory movements, fine movements, rearing and general locomotor activity were analyzed and processed using the PAS-HC data collection transferred to Excel. N (WT) = 10 and N (Het) = 13.

### Open-field test

Juvenile mice were originally placed in the center of open field apparatus (51 cm x 51 cm x 30 cm Plexiglas box) and recorded for 20 minutes. The time spent in the center zone of open field areas were recorded and analyzed with the video tracking system ANY-maze (Stoelting Company). N (WT) = 10 and N (Het) = 12.

### Light-dark box test

The apparatus (40 cm x 40 cm x 30 cm) consists of dark chamber (40 cm x 15 cm x 15 cm) and light chamber compartment. There is a door between two chambers which allow mice to freely move between chambers. Bright white lights illuminated the box from above as mice were placed in the box for 10 minutes. For analysis, the session was divided into two 5-min bins (0–5 min and 6–10 min), and data from the second bin (6–10 min) were used for statistical comparisons between WT and Het to reduce the variability caused by acute novelty stress. The test was recorded by ANY-maze video trace camera to determine the amount of time each mouse spent in the light chamber. N (WT) = 12 and N (Het) = 12.

### Prepulse inhibition (PPI) of the acoustic startle reflex

Startle response and pre-pulse inhibition were performed via SR-LAB-Startle Response System (San Diego Instruments). Adult mice are placed in a startle apparatus with sound-attenuated chamber. Whole-body startle movements were detected by a piezoelectric accelerometer mounted beneath the cage, converted to electrical signals, and digitized and stored by a computer. A loudspeaker mounted to the side of the cage produced white noise acoustic stimuli. A continuous 64 dB background noise was present throughout the test. The test began with a 5-minute acclimation to the apparatus, followed by 12 pulse-alone trials. To measure PPI, 12 blocks containing 6 trials were presented: 1 pulse-alone trial, 4 prepulse-pulse trials, and 1 no-stimulus trial. Pulse-alone trials consisted of a 40 ms startle pulse at 120 dB; prepulse-pulse trials consisted of a 20 ms prepulse at 66, 68, 72, or 76 dB followed 100 ms later by the pulse; and no-stimulus trials consisted of background noise only. Trials will be presented in pseudorandom order, with an inter-trial interval of 10-20 s (mean of 15 s). Startle response will be determined as the peak amplitude within 100 ms of the startle pulse onset. The entire test lasted 30 minutes. N (WT) = 6 and N (Het) = 9.

### 2 Object Novel object recognition test

Novel object recognition test was conducted in an open field arena (40 cm x 40 cm white Plexiglas box). The protocol used a 3-day paradigm that includes habituation, training, and testing phase. During habituation (day 1), adult mice were placed into the center of an open field arena and allowed to explore for 10 minutes. For training (day 2), two identical objects were placed on either side of the center of the arena, and the mice explored the arena for 10 minutes. For testing (day 3), a familiar object and a novel object (dissimilar object) were placed in the same position as in the training day and mice explored the arena for 10 minutes. ANY-maze was used to record each trial. The amount of time each mouse spent with their nose oriented toward the object within 2.5 cm of the object edge was considered ‘investigating time’. The discrimination index (DI) was calculated as: DI = (time investigating novel − time investigating familiar)/(time investigating novel + time investigating familiar). N (WT) = 12 and N (Het) = 12.

### Free-walk test

FreeWalkScan 2.0 (CleverSys Inc.) was used to assess mice gait. Adult mice move freely in a 40 cm × 40 cm × 30 cm (length × width × height) chamber. A high-speed camera below a clear bottom plate captures mouse movement for 5 minutes with red light in dark room. Videos are analyzed using FreewalkScanTM 2.0 software for various characteristic of gait, including front base (distance between sequential footprints of front limbs), stride length (distance between two sequential footprints of same paw). N (WT) = 5 and N (Het) = 8.

### Multiome sequencing and analysis

Pregnant dams and P70 mice were terminally anesthetized with an i.p. injection of Euthasol (270 mg/kg, 50 μl injection per 30 g mouse). Tissue from male and female mice were processed separately for E13.5 and P70 samples. Frozen E13.5 and P70 tissue from *Nkx2.1-Cre*;*Arx;sun1-GFP* mice were mechanically lysed, washed, filtered and sorted to generate single nuclei suspensions for the 10x Genomics Multiome assay as previously described^30,110^. We combined tissue from 2 E13.5 embryo for each Multiome reaction. Nuclei were reconstituted in 1x Nuclei buffer at a density of 3,000 nuclei/μl. 5 μl of this solution was then used for the 10x Genomics Single Cell Multiome Assay per manufacturer’s protocol. Our single cell Multiome workflow ‘multiome-wf’ can be found at github (https://github.com/NICHD-BSPC/multiome-wf) with documentation located here (https://nichd-bspc.github.io/multiome-wf/).

Sequencing analysis: Gene expression and open chromatin state were profiled using 10X Chromium Single Cell Multiome ATAC + Gene Expression kit (10X Genomics, 1000285). Joint libraries were simultaneously created by following standard protocol. Sequencing was conducted with paired-end (50 x 50 bp) using an Illumina HiSeq 2500 or NovaSeq 6000. The raw sequencing data were processed using Cell Ranger ARC (v2.0) pipeline. The cellranger-arc mkfastq command was used to generate the demultiplexed FASTQ files from BCL files. The sequencing reads were aligned to custom built mouse (GRCm38/mm10) reference genome using cellranger-arc count. The cellranger-arc count command was used to generate gene-by-barcode (snRNA-seq) and peak-by-barcode (snATAC-seq) matrices. The cellranger-arc aggr command, without depth normalization (--normalize = none) was used to aggregate sample datasets into a single feature-barcode matrix file.

Embryonic Multiome-seq data analysis: For snRNA-seq data analysis, the aggregated feature-barcode matrix was used as input to Seurat (v5.2.1)^111^ in R (v4.4.2, https://cran.r-project.org). Low-quality cells were firstly removed based on the following QC metrics: unique molecular identifiers (UMIs) and mitochondrial counts. Upper outliers using standard deviation ‘sd’ and remove cells automatically, while lower outliers kept cells with UMI counts more than 100 and mitochondrial counts more than sd. The standard Seurat workflow with default parameters was performed unless otherwise noted: NormalizeData (LogNormalize), FindVariableFeatures (nfeatures = 3000), ScaleData, RunPCA, FindNeighbors, FindClusters and RunUMAP (Dims = 1:30) steps. For snATAC-seq data analysis, the aggregated peak-by-barcode matrix was used as input to Signac (v1.14.0) ^112^ in R. Low-quality cells were removed based on the following QC metrics: number of chromatin accessibility peaks and transcription start site enrichment score. Upper outliers using standard deviation ‘sd’ and remove cells automatically, while lower outliers kept cells with more than 1000 chromatin accessibility peaks and the transcription start site enrichment score greater than 2. After filtering out the low-quality cells, we proceeded with normalization, identified highly variable peaks and reduced dimensions using the function of RunTFIDF, FindTopFeatures (min.cutoff = “q0”) and RunSVD in Signac with default parameters. The regular non-linear dimension reduction and clustering were performed by RunUMAP (dims = 2:30), FindNeighbors (dims = 2:30) and FindClusters in Seurat. The gene activity scores were calculated using the GeneActivity function and added to Seurat object. Multimodal analysis: The shared cells across the two modalities (snRNA-seq and snATAC-seq datasets) were merged, and the RNA and ATAC dimensionality reduction was recomputed using default parameters. Multimodal data integration was performed by finding the Weighted Nearest Neighbor (WNN)^113^ using Seurat::FindMultiModalNeighbors. The dimensionality reduction was separately set to 1:30 (snRNA assay) and 2:30 (snATAC assay). The joint UMAP and clustering was performed using WNN graph with default parameters. Cell types of E13.5 MGE were identified with FindAllMarkers function within clusters and integrated with mature MGE cell type biomarkers. Notably, cells didn’t belong to MGE lineage were also removed for further analysis.

Adult Multiome-seq data analysis: For snRNA-seq data analysis, the aggregated feature-barcode matrix was used as input to Seurat (v5.2.1)^111^ in R (v4.4.2, https://cran.r-project.org). Cells with percent of ribosomal counts less than 2, percent of mitochondrial counts less than 1, RNA feature counts between 1000 and 7500, and UMIs counts less than 40000 were kept. The Seurat workflow was performed as previous described except with FindVariableFeatures (nfeatures = 2000). For snATAC-seq data analysis, the aggregated peak-by-barcode matrix was used as input to Signac (v1.14.0)^112^ in R. Cells with more than 1000 chromatin accessibility peaks, the transcription start site enrichment score greater than 2 and nucleosome signal less than 4 were kept. Multimodal analysis was performed as pervious described except with FindClusters (resolution = 1.2). Joint clusters with few cell numbers were removed.

Annotation analysis and merge datasets: All cell types were defined using the RNA expression only. Cell type annotation was based on clusters marker using FindAllMarkers function and public articles. Filtered RNA assay of Embryonic (E13.5) and adult (P70) datasets were merged and run default Seurat workflow, while filtered ATAC assay of both datasets were also merged and run previous described steps.

Pseudotime analysis: Single cell trajectory analysis of E13.5 snMultiome-seq datasets was performed using Monocle 3 (v1.3.7)^114^ (https://cole-trapnell-lab.github.io/monocle3/docs/trajectories/). Seurat objects were converted to a CellDataSet object using as.cell_data_set function and RNA counts were extracted. The reduce dimensionality of CellDataSet object was set to WNN.UMAP of Seurat object. Learning trajectory graph using learn_graph function. The root of trajectory was set to *Nes+* clusters and order cells using order_cells function with default parameters. Plotting trajectory colored by pseudotime with plot_cells function.

Visualization: Uniform Manifold Approximation and Projection (UMAP) coordinates and WNN clustering, computed by Seurat on multimodal integrated datasets, were visualized using DimPlot function. The expression of genes of interest was visualized using FeaturePlot, VlnPlot or DoHeatmap function. Packages ggplot2 (v3.5.1) and patchwork (v1.3.0) were used to visualize during Multiome datasets analysis process.

### Visium HD Spatial Gene Expression

Sample preparation and spatial transcriptomic sequencing: All brain tissues used in Visium HD were cryosectioned at 12 μm, which were taken from Fixed Frozen Tissue (FFT) blocks following the Visium HD FFT Preparation Handbook (CG000764) except one batch (one P120 female WT and one Het) that was taken from formalin fixed & paraffin embedded (FFPE) block following the Visium HD FFPE Preparation Handbook (CG000684). Hematoxylin (Sigma-Aldrich, GHS216) and Alcoholic Eosin (Sigma-Aldrich, HT110132) (H&E) staining and imaging were performed using preparation Handbook, then processed and sequenced following the Visium HD Spatial Gene Expression Reagent Kits User Guide (CG000685) with Visium HD Slides (1000670), Visium Mouse Transcriptome Probes v2 (1000667), Visium HD Reagents (1000668) and Visium HD Cassettes (1000669).

Space Ranger pipeline: Loupe Browser (v9.0.0) was used to align the CytAssist and H&E images, and Space Ranger (v 4.0) was used for analysis. Sequences were aligned with mm10 reference genome to gene expression counts using Space Ranger default setting. Space Ranger outputs Visium HD data binned to 2 μm, 8 μm and 16 μm resolution and cell segmented output. Unless otherwise specified, downstream analyses for all samples were performed on the 8 μm resolution datasets, which were recommended by 10x Genomics due to enriched UMI reads and closed single-cell scale compared to other resolution. Downstream analyses for E13.5 brain samples and E15.5 cortex were performed using cell segmented datasets to allow transcripts to be assigned at the cell level. Key metrics of 8 μm resolution (Number of binned squares under tissue, mean reads, mean UMIs and total genes detected) and cell segmentation metrics (number of cells, mean reads per cell and median genes per cell) are listed (Supplementary Table 1).

Spatial transcriptomic data processing: 8 μm resolution Visium HD spatial datasets were analyzed with Seurat (v5.4.0) following standard Visium HD workflow (https://satijalab.org/seurat/articles/visiumhd_analysis_vignette). Brain dataset was loaded with Load10X_Spatial function with bin.size = 8. Unless otherwise specified, LogNormalize, FindVariableFeatures and ScaleData were performed for “Spatial.008um” assay with default parameters. The sketch-based analysis approach was adopted with cells = 50,000 and perform clustering workflow, then sketch assay was projected to “Spatial.008um” assay. To identifying and segmenting spatial domain, Building Aggregates with a Neighborhood Kernel and Spatial Yardstick (BANKSY) was performed with default parameters. After dimensional reduction and clustering, we subset out cortex and hippocampus based on BANKSY clusters for further downstream analysis. To accurately annotate spatial data with scRNA-seq reference, Robust Cell Type Decomposition method was introduced. Here, cortex or hippocampus were separately integrated with ALLEN scRNA-seq dataset using RCTD with default workflows. Output from RCTD contained the weights for cell types in the reference in each spot and the class (‘singlet’, ‘doublet_certain’, ‘doublet_uncertain’ and ‘reject’) of each bin. Only the ‘singlet’ (one cell type) spots and a minute UMI threshold of 100 for a spot were used for further analysis. Upper limitation of UMI depended on datasets. When needed, cell segmentations Visium HD spatial datasets were analyzed with Seurat (v5.4.0) following standard workflow (https://satijalab.org/seurat/articles/visiumhd_analysis_cell_segmentations). Brain datasets were loaded with Load10X_Spatial with bin.size = segmented_outputs. Spatial transcripts of Visium HD datasets were visualized by using Seurat and Loupe Browser.

### Differential analysis

Multiome-seq datasets: Differentially expressed genes between genotypes were computed using the Wilcoxon rank sum test implemented in the Seurat FindMarkers function with Bonferroni correction (significance: log_2_FC > ± 0.2, adjusted P-value < 10^-6^) for the E13.5 MGE and P70 cortex datasets. Differentially expressed peaks between different genotypes were computed using the Wilcoxon rank sum test implemented in the Seurat FindMarkers function with Bonferroni correction (significance: log_2_FC > ± 0.2, adjusted P-value < 10^-6^). The EnhancedVolcano function from the EnhancedVolcano package (v1.24.0, https://github.com/kevinblighe/EnhancedVolcano) was used to visualize the DEGs between different conditions per population and parameters.

Spatial transcriptomic datasets: Unless otherwise specified, differentially expressed genes between genotypes were computed using the Wilcoxon rank sum test implemented in the Seurat FindMarkers function with Bonferroni correction (significance: log_2_FC > ± 0.2, adjusted P-value < 0.05). When compared E15.5 cortex DEGs, parameters were set to adjusted P-value < 10^-50^ and log_2_FC > ± 0.1 for significant determination. When compared E15.5 subpallium DEGs, parameters were set to adjusted P-value < 10^-6^ and log_2_FC > ± 0.2 for significant determination.

### Cell-cell communication analysis

Cell-cell communication was analyzed in E15.5 male cortex using CellChat ^91^ version 3 (Spatial CellChat) following tutorial (SpatialCellChat analysis of spatial transcriptomics data). E15.5 segment Visium HD datasets of cortex from WT and cKO were input and only part of dorsal cortex (around 5, 000 cells) was kept and used to create individual spatial CellChat object. All CellChatDB database except for “Non-protein Signaling” was used for cell-cell communication analysis. The communication probability was computed with 250 μm interaction range and 10 μm contact range. Cell-cell communication at signaling pathway level and network centrality scores were computed with default setting. Spatial CellChat objects from WT and cKO were integrated and comparison analysis was performed following tutorial (Comparison analysis of multiple datasets using CellChat).

### Enrichment analysis

Gene ontology pathway analysis was performed on each group’s (e.g., genotype and sex) DEGs with adjusted p values <0.05 using the enrichGO function from the clusterProfiler package (v4.14.0)^115^. The parameters were set as following: OrgDb = org.Mm.eg.db (v3.20.0), keyType = “SYMBOL”, ont = “BP”, universe = “All detected genes in specific population”. ClusterProfiler::simplify function was also used to reduce the redundancy among enriched terms.

### Data & Code availability

snMultiome sequencing and spatial transcriptomics data generated in this study will be deposited in the Gene Expression Omnibus (GEO) and made publicly available upon publication of the manuscript. The code for snMultiome and spatial transcriptomics analysis will be publicly available on Github upon publication of the manuscript. Additional source data, statistics, etc. used in this study will be available at Synapse.org upon publication of this manuscript. For any inquiries about data accessibility and analyses, please.

## ACKNOWLEDGEMENTS

We thank all members of the Section on Cellular and Molecular Neurodevelopment, as well as Drs. Todd MacFarlan & Pedro Rocha for discussion and comments on this project and manuscript. We thank Drs. Eric Marsh & Jeffrey Golden for the *Arx-floxed* mice. We thank the NICHD Molecular Genomics Core, specifically Vivek Mahadevan, Tianwei Li and James Iben; the NINDS and NICHD animal facility for mouse husbandry assistance; the NICHD Bioinformatics and Scientific Programming Core (BSPC), specifically Apratim Mitra and Ryan Dale; the NICHD Microscopy and Imaging Core. This work utilized the computational resources of the NIH HPC Biowulf cluster (http://hpc.nih.gov). This work was supported, in part, by the NIMH IRP Rodent Behavioral Core (MH002952). This project was funded by NICHD intramural projects HD008962 (T.J.P.); NICHD Intramural Research Fellowship (J.L.); NICHD Career Development Award (J.L.). This research was supported by the Intramural Research Program of the National Institutes of Health (NIH). The contributions of the NIH author(s) are considered Works of the United States Government. The findings and conclusions presented in this paper are those of the author(s) and do not necessarily reflect the views of the NIH or the U.S. Department of Health and Human Services.

## AUTHOR CONTRIBUTIONS STATEMENT

Conceptualization – J.L. and T.J.P.; Investigation – J.L., V.G., A.M., A.T., and Y.Z.; Formal analysis – J.L., V.G., A.M., and A.T.; Supervision –T.J.P.; Funding acquisition–J.L. and T.J.P.; Writing, original draft – J.L. and T.J.P.; Writing, review & editing – all authors.

## COMPETING INTERESTS STATEMENT

The authors declare no competing interests.

### Declaration of generative AI and AI-assisted technologies in the writing process

During the preparation of this work, the authors used Claude to revise grammar and refine the text. After using this tool, the authors reviewed and edited the content as needed and take full responsibility.

**Figure S1.**
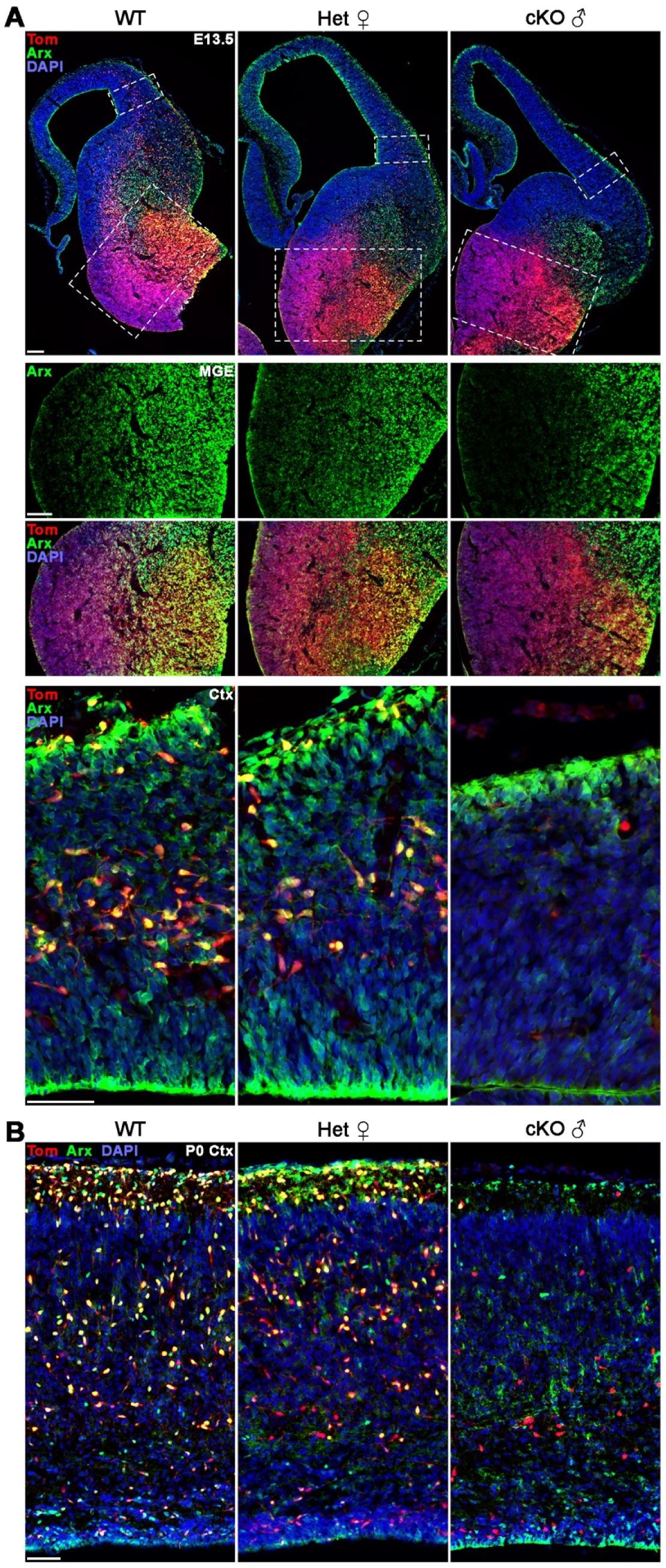
Loss of Arx protein in MGE-derived cells in cKO male mice. **A.** Arx protein is reduced in the MGE of E13.5 Het female and cKO male mice. Arx is expressed in migratory MGE-derived interneurons in the cortex of E13.5 WT and Het female mice, but lost in cKO male mice. White dotted boxes in top row indicate higher magnification images of MGE and lateral cortex in lower images. Scale bars = 100 μm for hemisphere and MGE, 50 μm for cortex. **B.** Tom+ MGE-derived cortical interneurons express Arx protein in P0 WT and Het female mice, but these cells are Arx-negative in cKO male mice. Scale bar = 50 μm.

**Figure S2.**
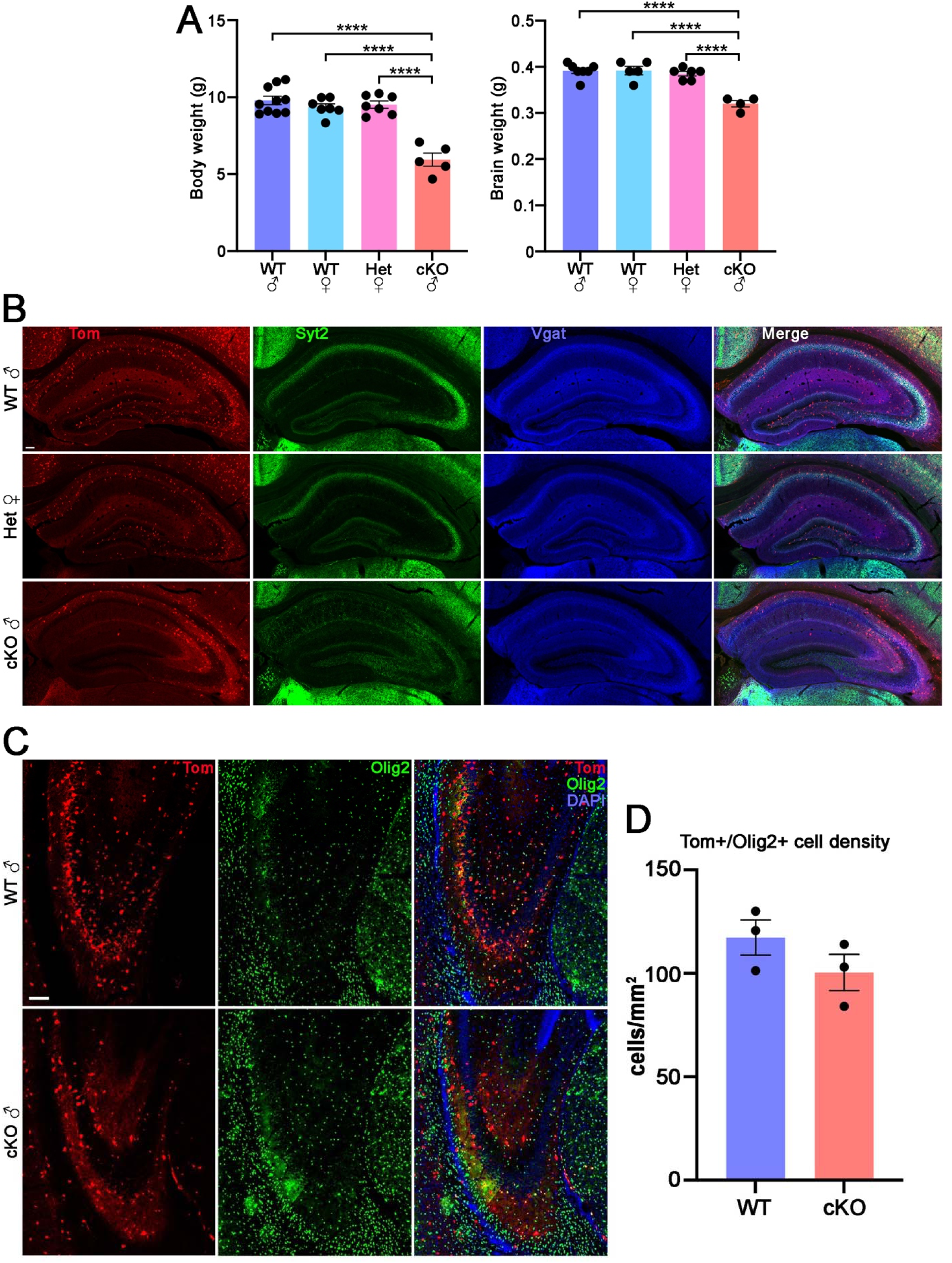
Decreased body weight, brain weight and synaptotagmin-2 expression in Arx mutant mice. **A.** Quantification of body weight (left) and brain weight (right) from P21 WT, Het and cKO mice. n = 10 WT male, 7 WT female, 7 Het female and 5 cKO male for body weight; n = 7 WT male, 5 WT female, 6 Het female and 4 cKO male for brain weight. **B.** Reduction of synaptotagmin-2 (Syt2) in the hippocampus of P21 Arx mutant mice. **C-D.** Images highlighting Tom+/Olig2+ oligodendrocytes in CA3 in P21 male WT and cKO mice (C), and quantification of these cells (D). n = 3 for each genotype. Ordinary one-way ANOVA followed by Tukey’s multiple comparison tests (A) and Welch’s 2-tailed t-test (D) were used: **** = p ≤ .0001. Scale bar = 100 μm.

**Figure S3.**
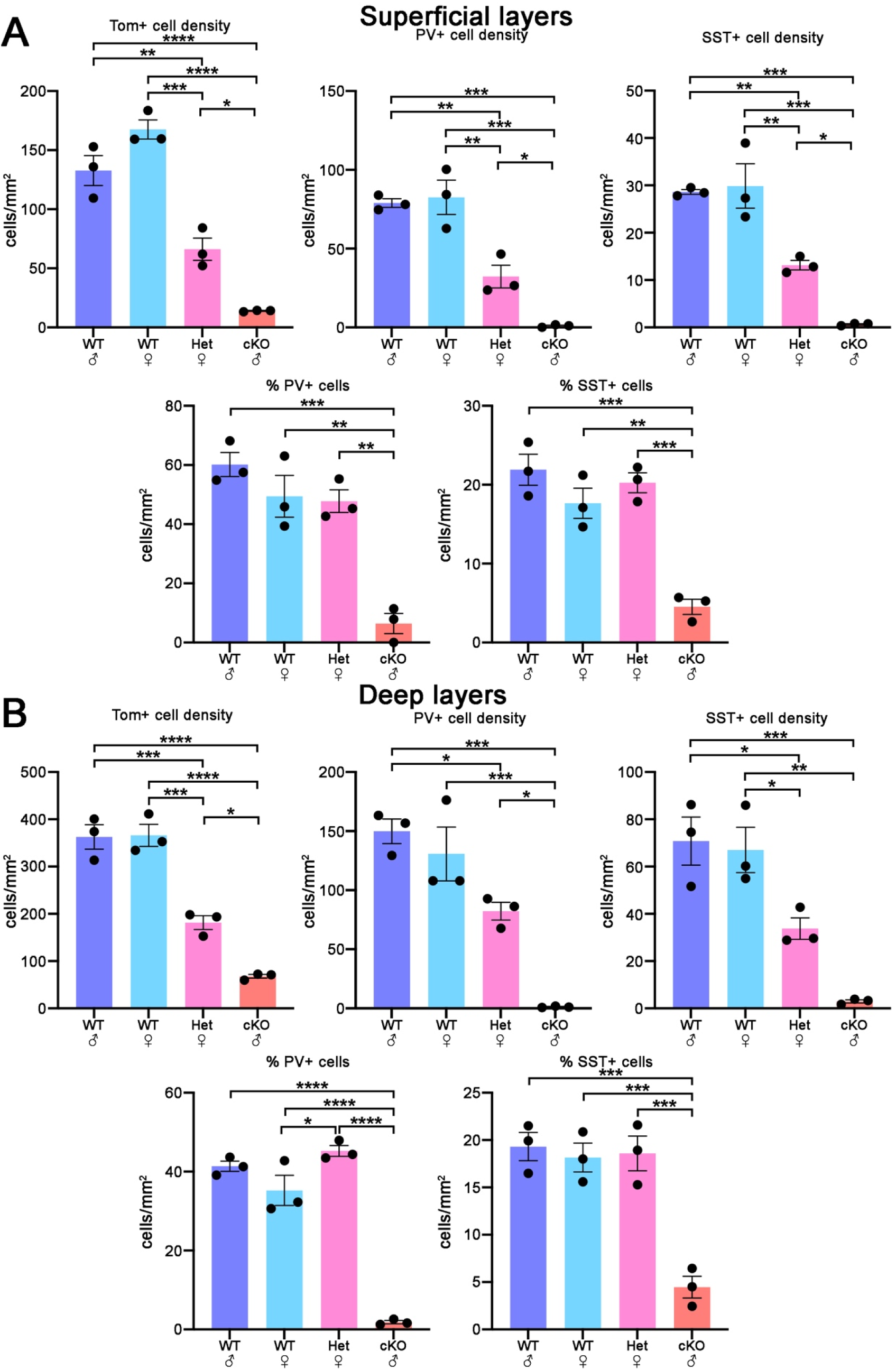
Decreased MGE-derived cortical interneurons in the superficial and deep layers of Arx mutant mice. **A-B**. Graphs depicting the density of Tom+, PV+ and SST+ cells (top) and the percent of Tom+ cells expressing PV or SST (bottom) in the superficial layers I-III (A) and deep layers IV-VI (B) of P21 WT, Het and cKO mice. n = 3 for each condition. All stats are one-way ANOVA followed by Tukey’s multiple comparison tests: * = p ≤ .05, ** = p ≤ .01, *** = p ≤ .001, **** = p ≤ .0001.

**Figure S4.**
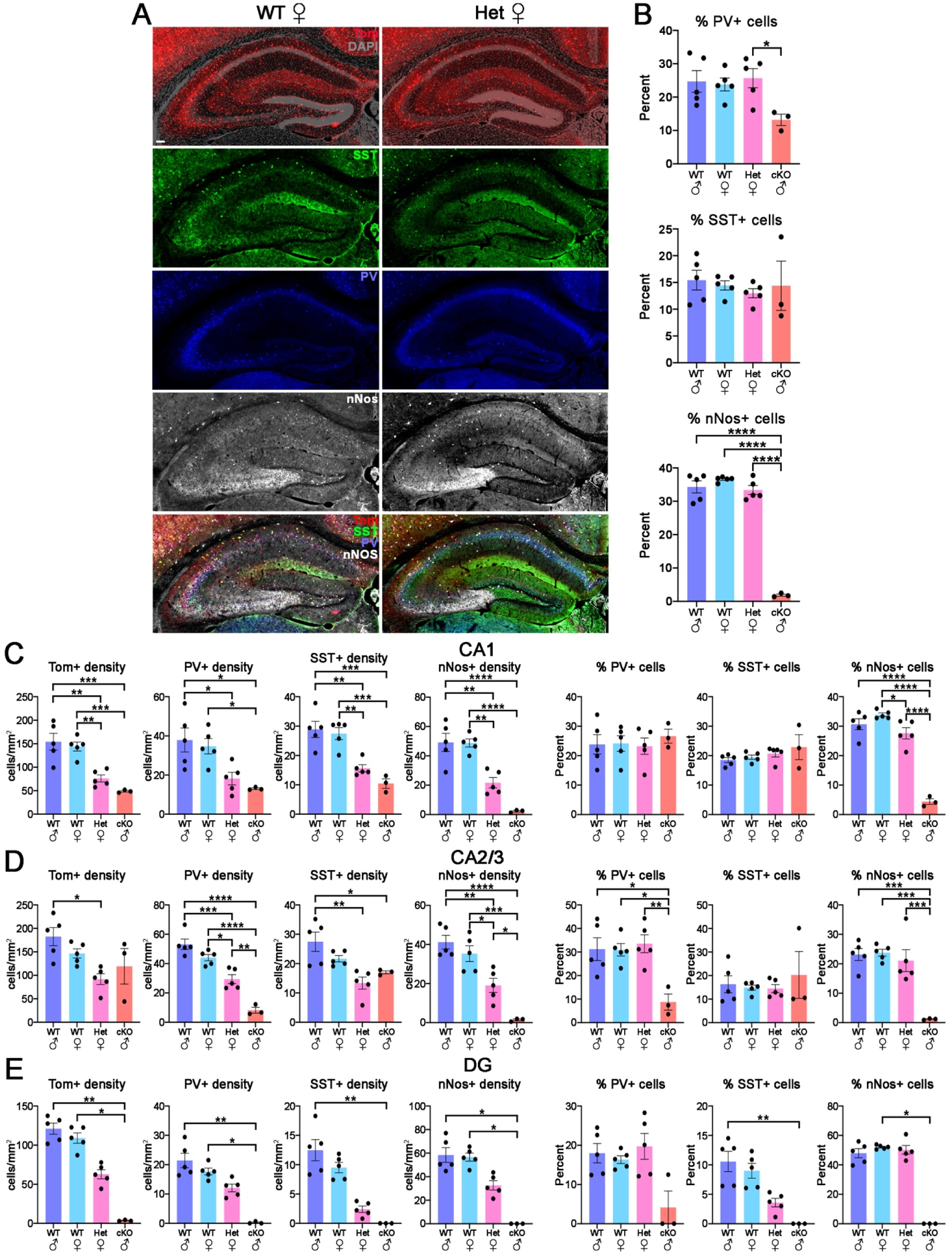
Decreased MGE-derived hippocampus interneurons in Arx mutant mice. **A.** P21 WT and Het female hippocampus mice stained for PV, SST and nNos. Scale bar = 100 μm. **B.** Percent of hippocampal Tom+ cells expressing PV, SST or nNos in the hippocampus. **C-E.** Quantification of cell density of Tom+ PV+, SST+ and nNos+ cells (left) and percent of Tom+ cells expressing PV, SST or nNos (right) in the CA1 (C), CA2-3 (D) and DG (E). n = 5 WT male, 5 WT female, 5 Het female and 3 cKO male mice. Ordinary one-way ANOVA followed by Tukey’s multiple comparison tests or Holm-Sidak’s multiple comparisons test (B, C, D) and Kruskal-Wallis test followed by Dunn’s multiple comparisons test (E) based on normal distribution were used: * = p ≤ .05, ** = p ≤ .01, *** = p ≤ .001, **** = p ≤ .0001.

**Figure S5.**
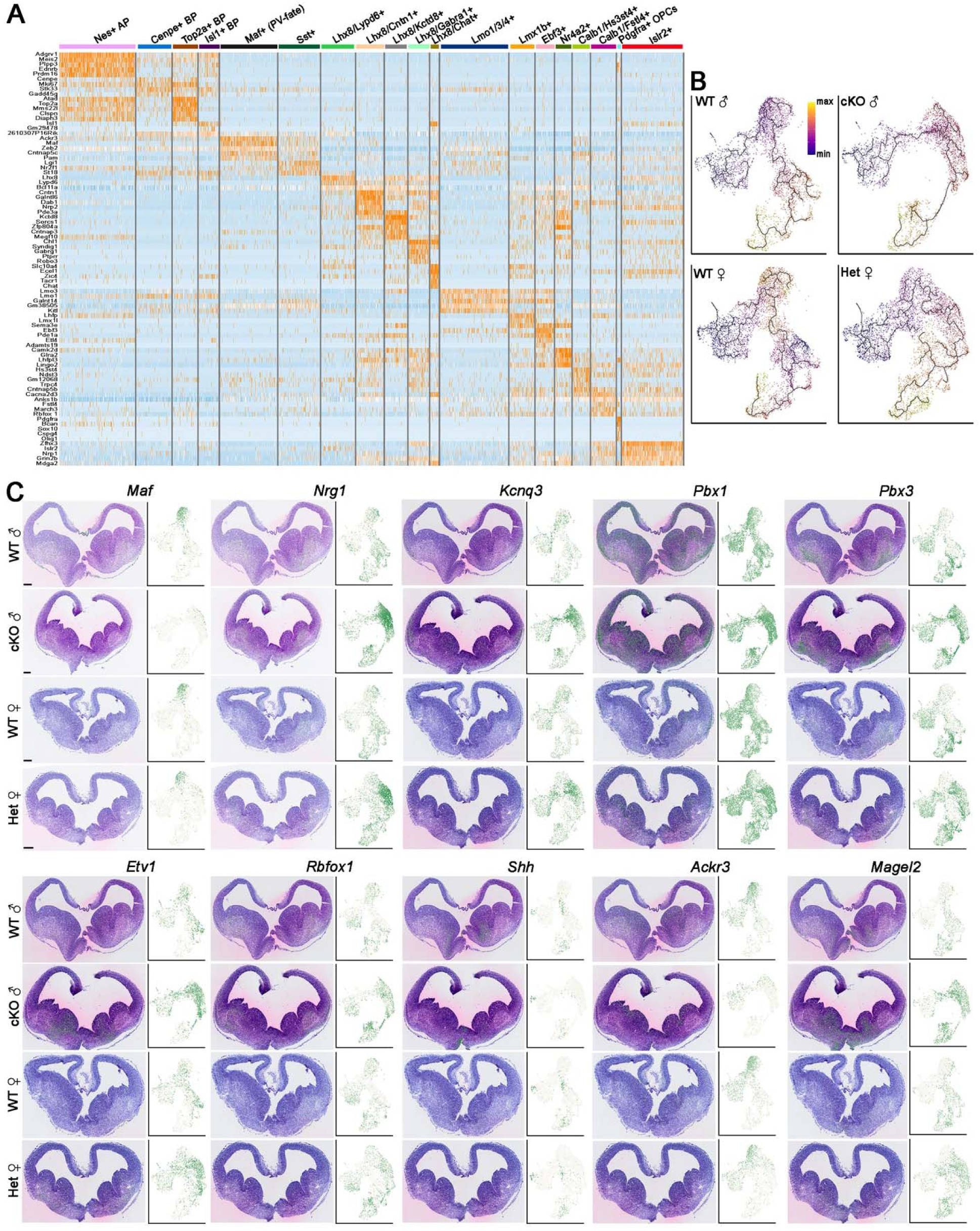
Genetically defined E13.5 cell types and spatial gene expression patterns. **A.** Heatmap depicting genes enriched in cell clusters of E13.5 MGE from snRNA-seq datasets. **B.** Integrated RNA and ATAC UMAP plots of E13.5 MGE cells showing pseudotime developmental trajectory using Monocle 3. **C.** mRNA expression of DEGs in E13.5 brains visualized in spatial transcriptomics data (left) and UMAP plots from snMultiome data (right). Scale bar = 200 μm.

**Figure S6.**
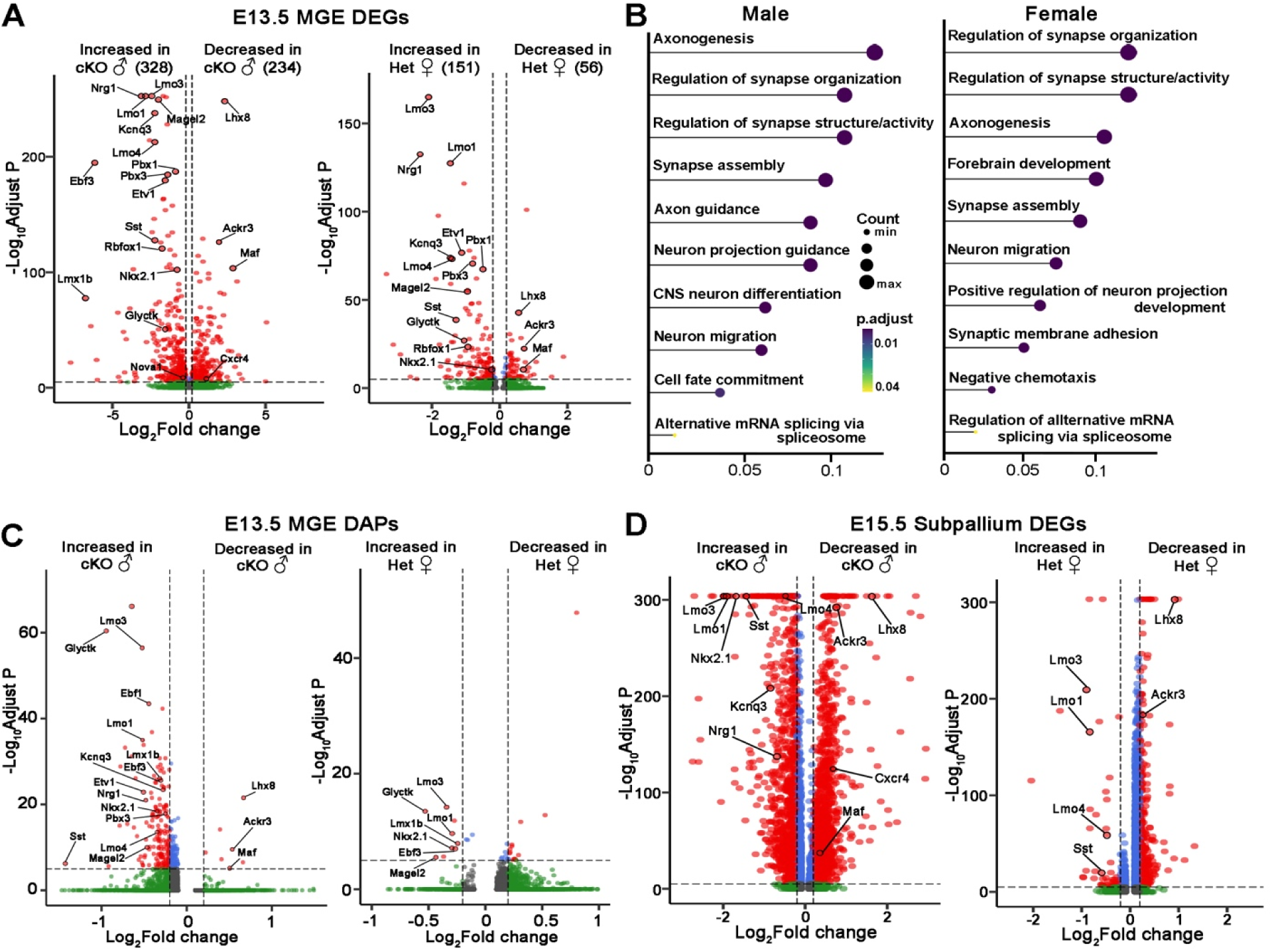
Differentially expression genes and differentially accessible peaks in embryonic Arx mutant brains. **A.** Volcano plot depicting DEGs from snRNA data in E13.5 cKO male (left) and Het female (right) MGE compared to WT. **B.** clusterProfiler GO enrichment top hits of biological processes for DEGs of E13.5 MGE in male (left) and female (right) mice. **C.** Volcano plots depicting differentially accessible peaks from snATAC data in E13.5 cKO (left) and Het (right) MGE compared to WT. **D.** Volcano plot depicting DEGs from spatial transcriptomics data in E15.5 cKO (left) and Het (right) subpallium compared to WT. Two-sided Wilcoxon rank sum test with Bonferroni correction (A, C, D) and one-sided Fisher’s exact with BH correction (B) were used.

**Figure S7.**
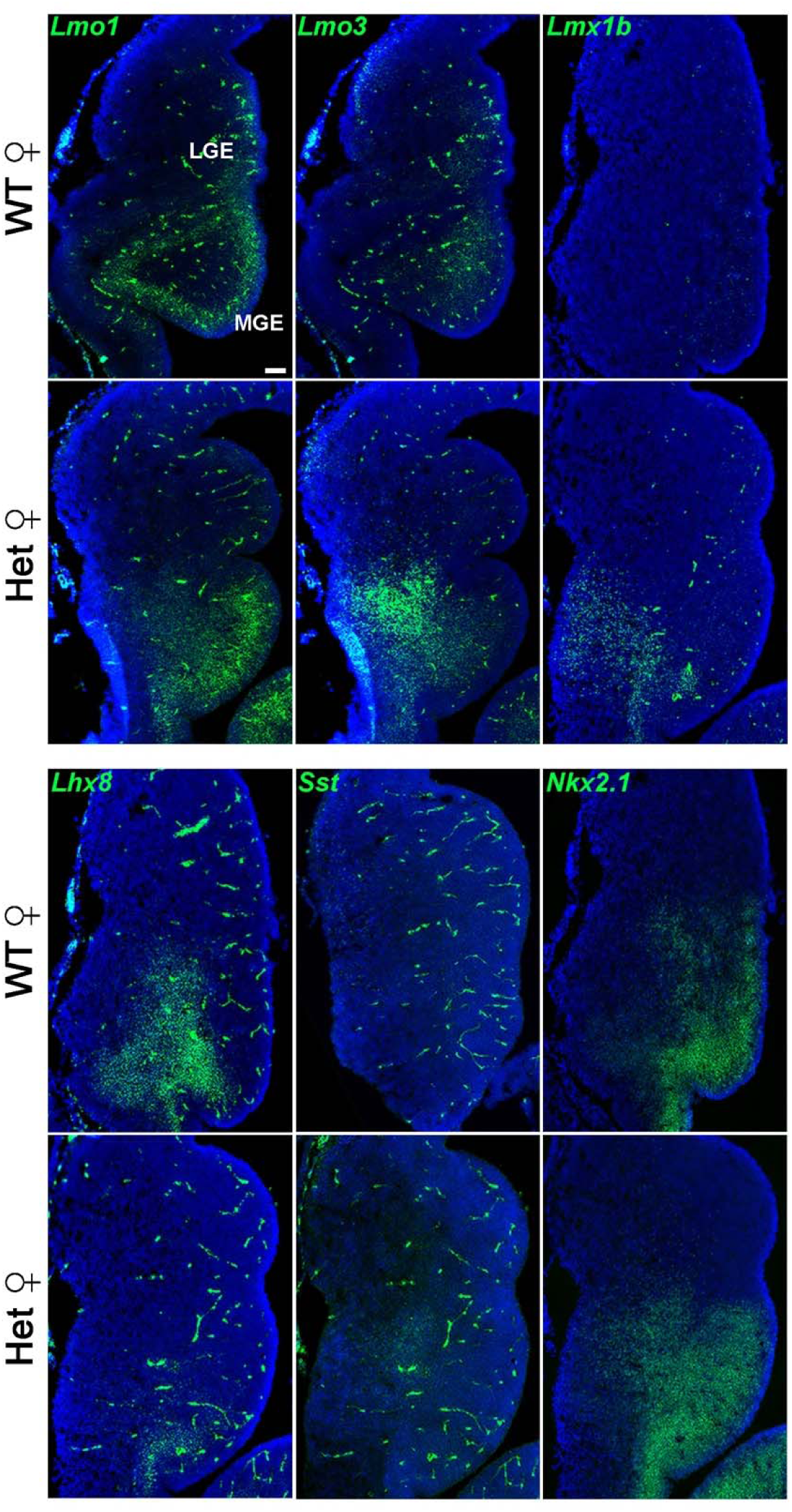
FISH of DEGs in E13.5 WT and Het female MGE. FISH confirmed an increase of *Lmo1, Lmo3, Lmx1b, Sst* and *Nkx2-1,* and a reduction of *Lhx8* in E13.5 female Het MGE compared to WT. Scale bar = 100 μm.

**Figure S8.**
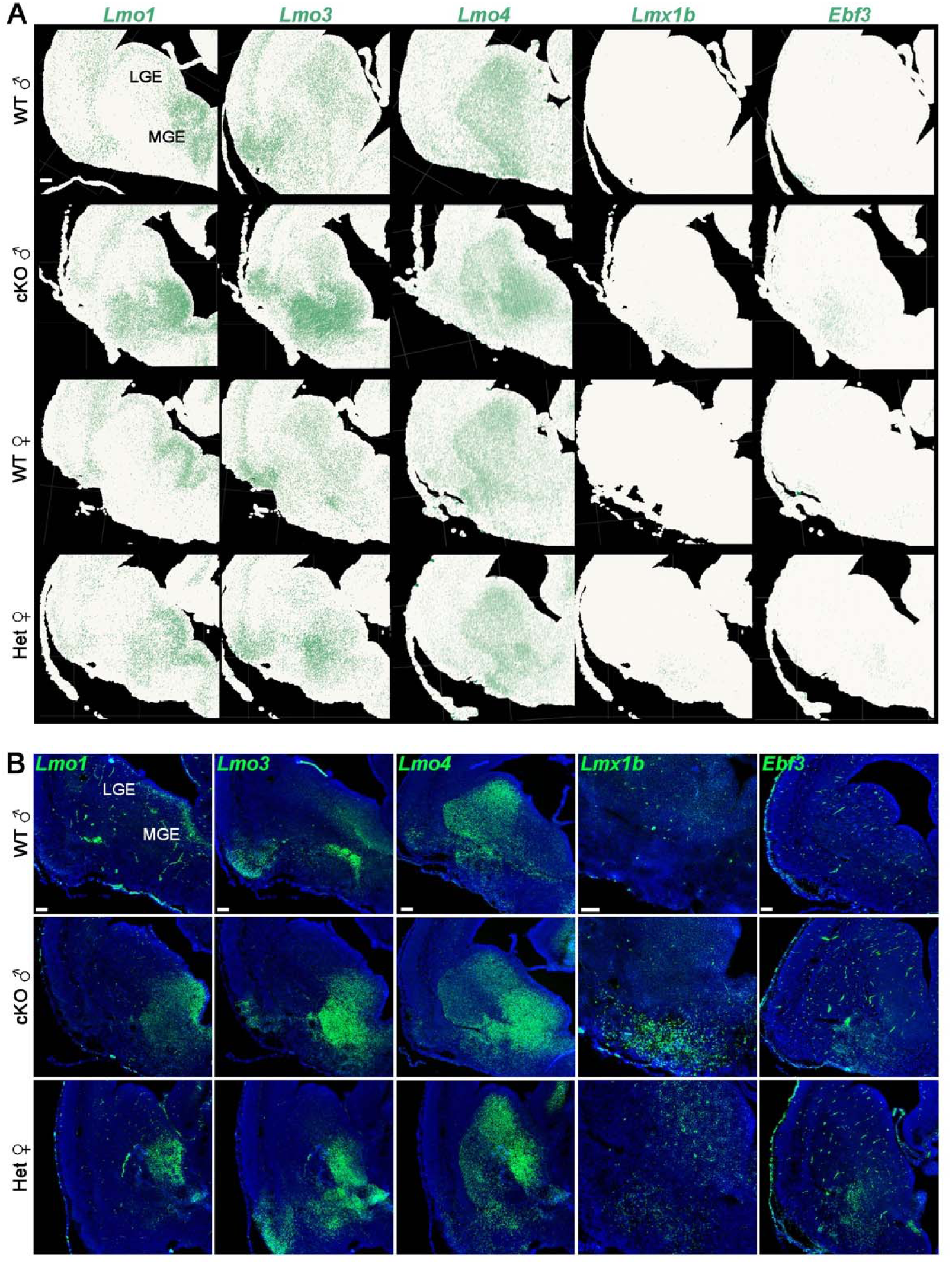
Spatial distribution of upregulated DEGs in E15.5 Arx mutant mice. **A-B**. Spatial distribution of transcripts in the E15.5 mouse brain from spatial transcriptomics data (A) and FISH visualization (B) for *Lmo1, Lmo3, Lmo4, Lmx1b* and *Ebf3.* Scale bar = 100 μm.

**Figure S9.**
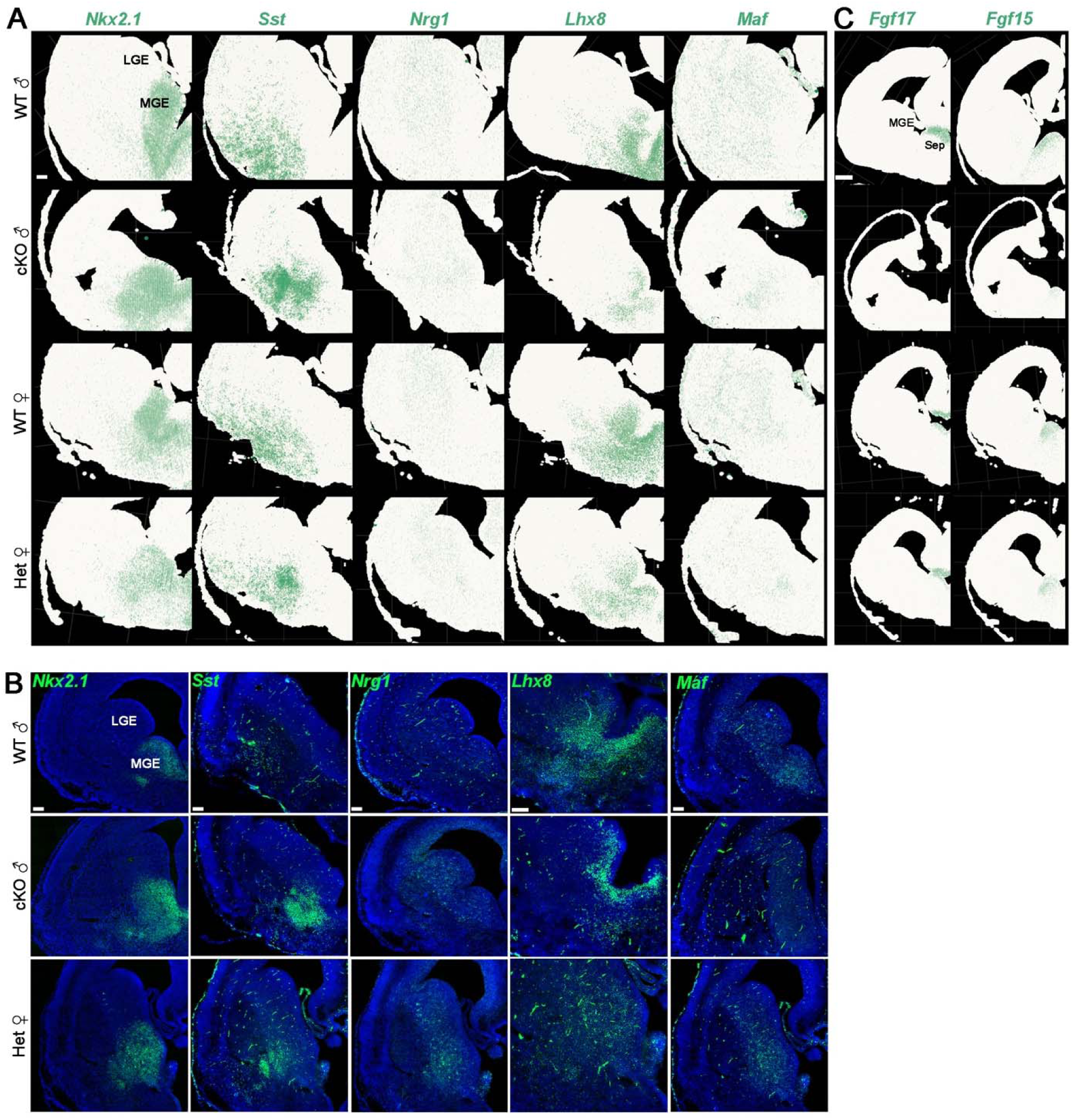
Spatial distribution of DEGs in E15.5 subpallium. **A-B**. Spatial distribution of transcripts in the E15.5 mouse brain from spatial transcriptomics data (A) and FISH visualization (B) for *Nkx2.1, Sst, Nrg1, Lhx8* and *Maf.* Scale bar = 100 μm. **C.** Spatial distribution of *Fgf17* and *Fgf15* in the E15.5 mouse brain from spatial transcriptomics data. Scale bar = 250 μm.

**Figure S10.**
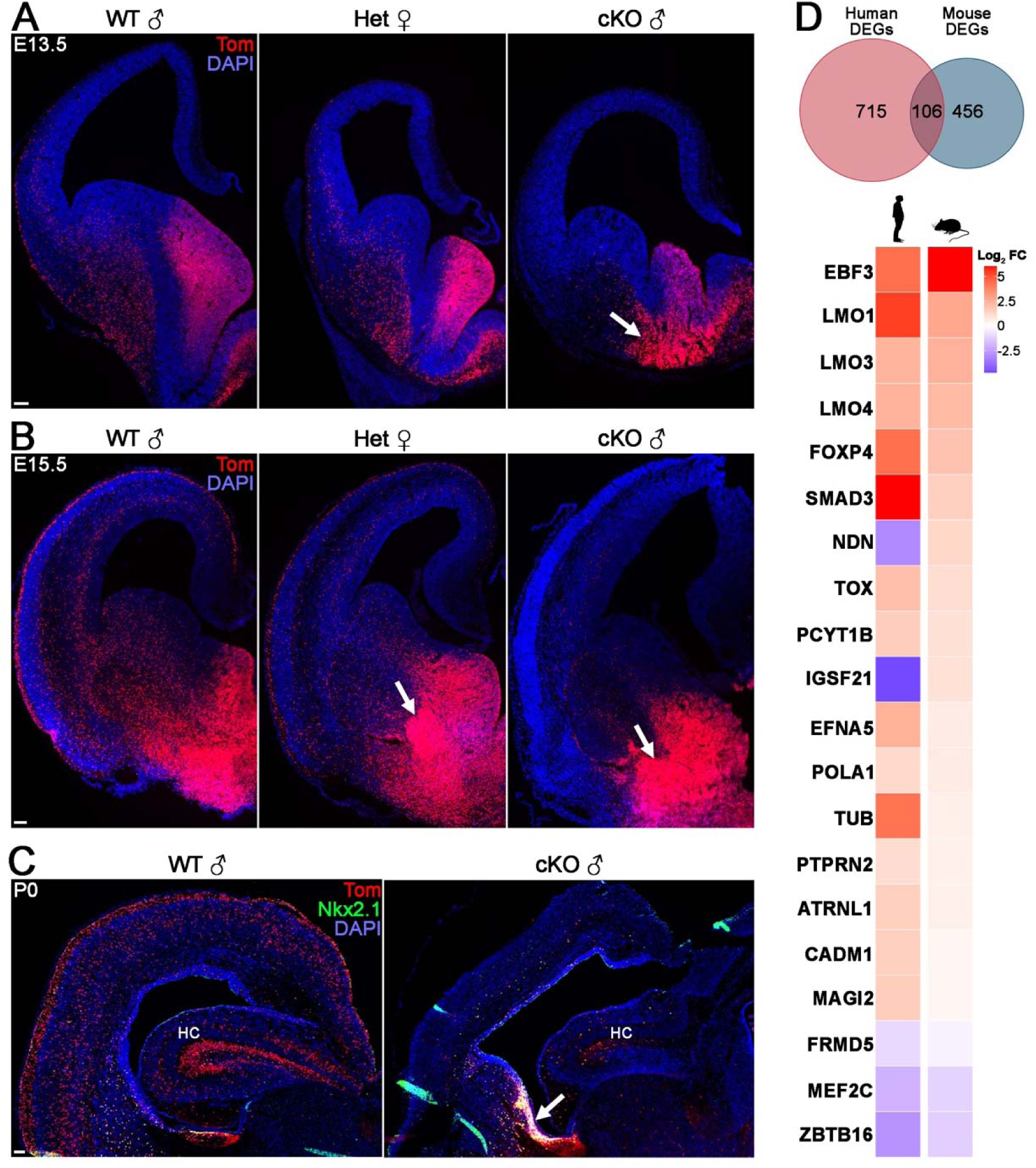
Migration defect from MGE-derived interneurons in Arx mutant mice. **A-B**. Tom+ MGE-derived interneurons get stuck in the MGE mantle in E13.5 (A) and E15.5 (B) Arx cKO mice, and Tom+ cells remain confined to the ventral forebrain periventricular zone at P0 (C). **D.** Venn diagram (top) and heatmap (bottom) showing the top hits of shared dysregulated genes upon Arx loss in both human neural progenitor cultures and E13.5 mouse MGE. Scale bar = 100 μm.

**Figure S11.**
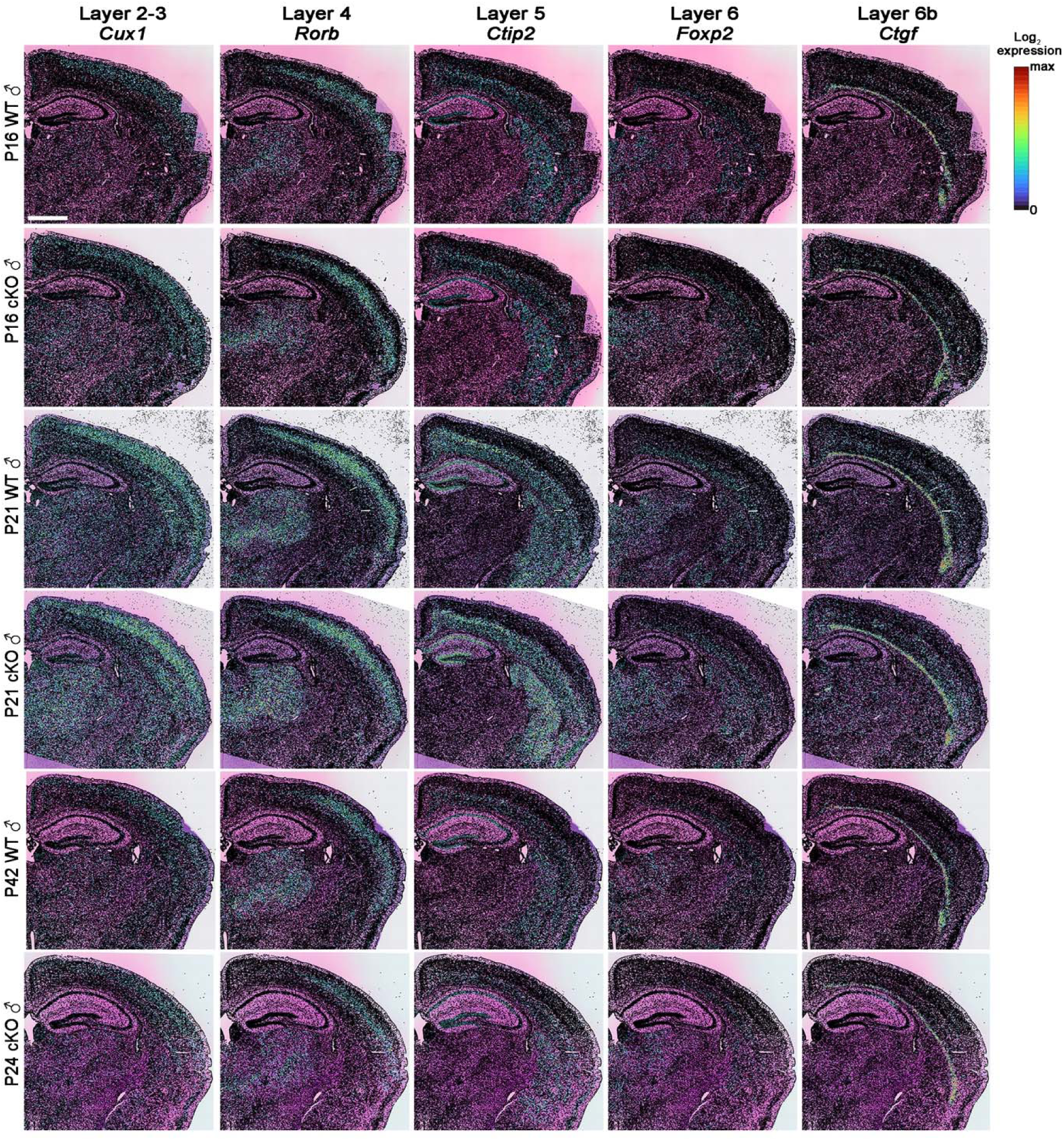
Spatial distribution of cortical layer markers in WT and cKO male postnatal brains. Spatial distribution of mRNA in P16, P21 and P42 WT and cKO male mice for *Cux1* (layer II-III), *Rorb* (layer IV), *Ctip2* (layer V), *Foxp2* (layer VI), and *Ctgf* (layer VIb) visualized in cell segmented spatial transcriptomics data. Scale bar = 1 mm.

**Figure S12.**
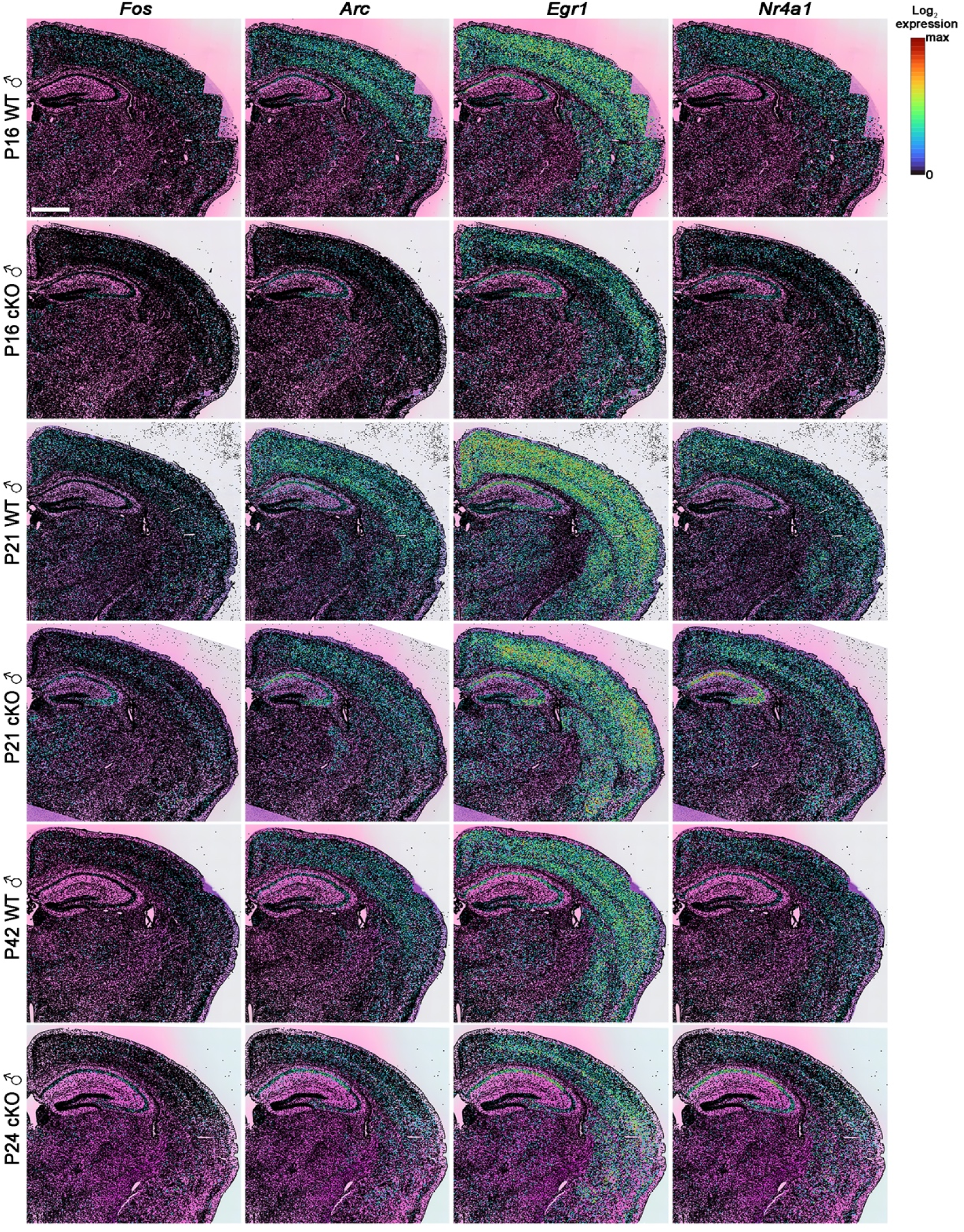
Spatial distribution of activity related genes in WT and cKO male postnatal brains. Spatial distribution of mRNA in P16, P21 and P42 WT and cKO male mice for *Fos*, *Arc*, *Egr1* and *Nr4a1* visualized in cell segmented spatial transcriptomics data. Scale bar = 1 mm.

**Figure S13.**
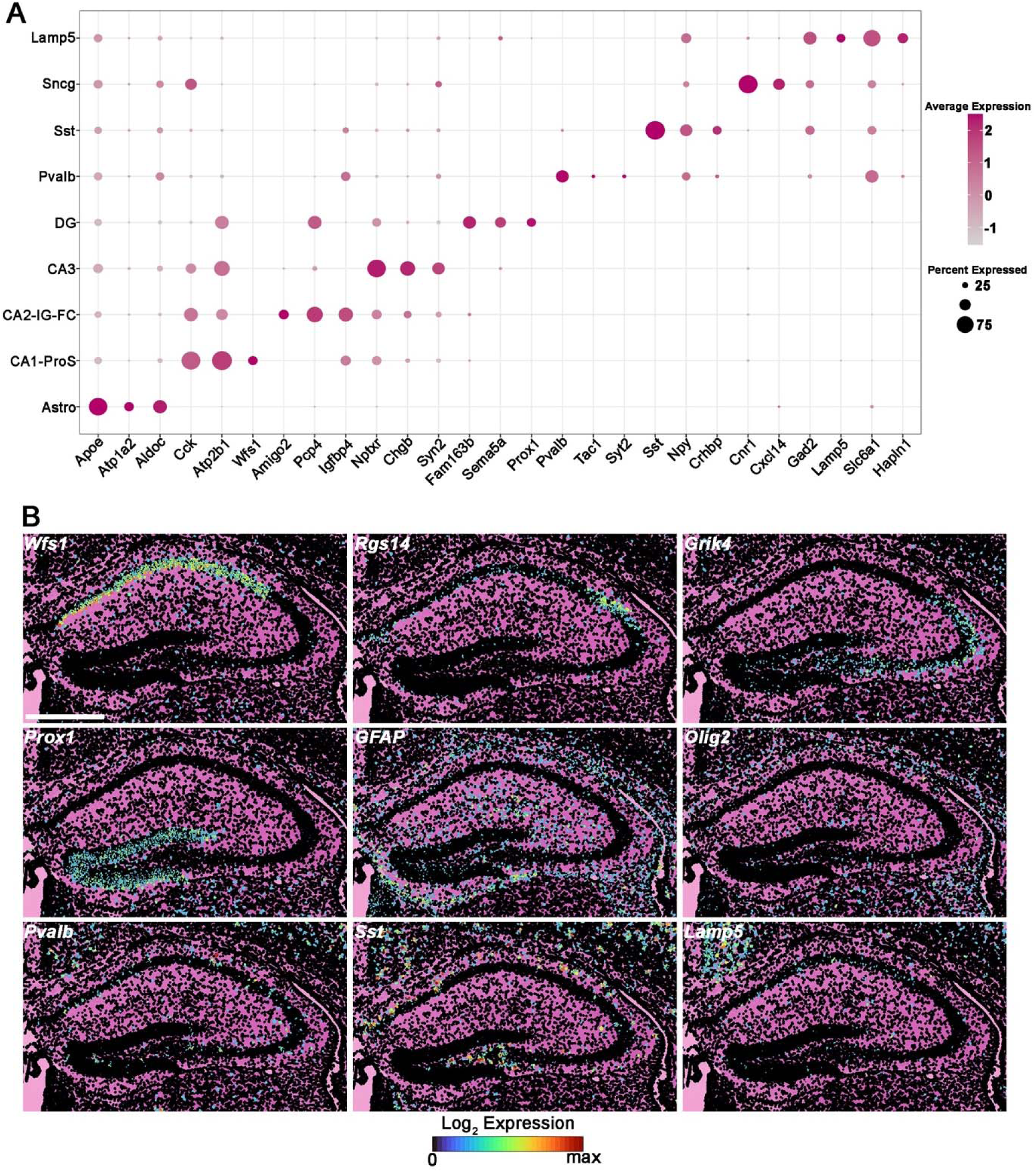
Genes defining hippocampal subdomains and cell types. **A.** Dot plot for depicting genes enriched in hippocampal subdomains and cell types. **B.** Spatial distribution of mRNA in P16 WT male hippocampus visualized in cell segmented spatial transcriptomics data. Scale bar = 0.5 mm.

**Figure S14.**
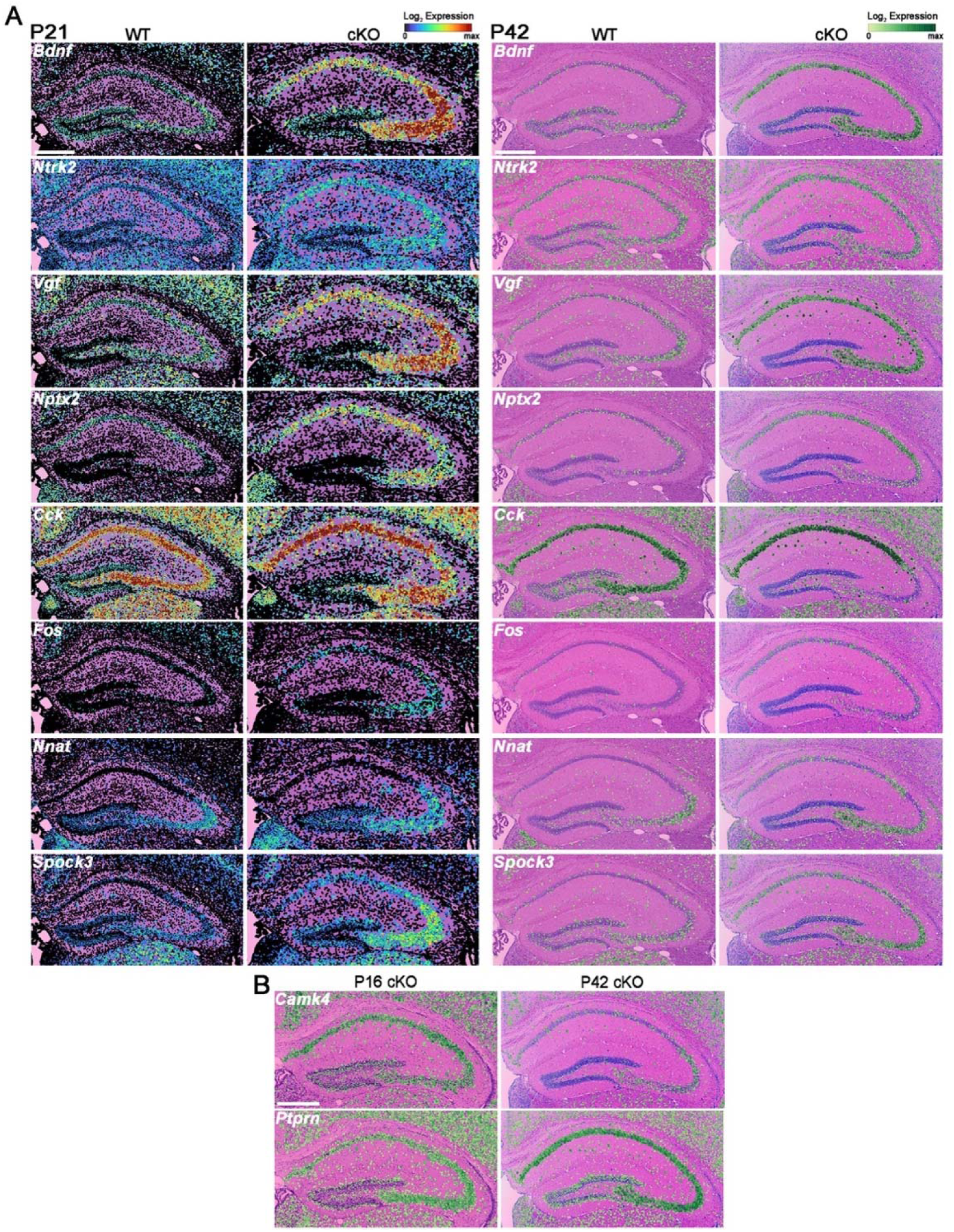
Spatial distribution of seizure related genes transcripts in WT and cKO male hippocampus. **A.** Spatial distribution of mRNA in P21 (left) and P42 (right) WT and cKO male hippocampus visualized in cell segmented spatial transcriptomics data. **B.** Spatial distribution of mRNA in P16 (left) and P42 (right) cKO male hippocampus. Scale bar = 0.5 mm.

**Figure S15.**
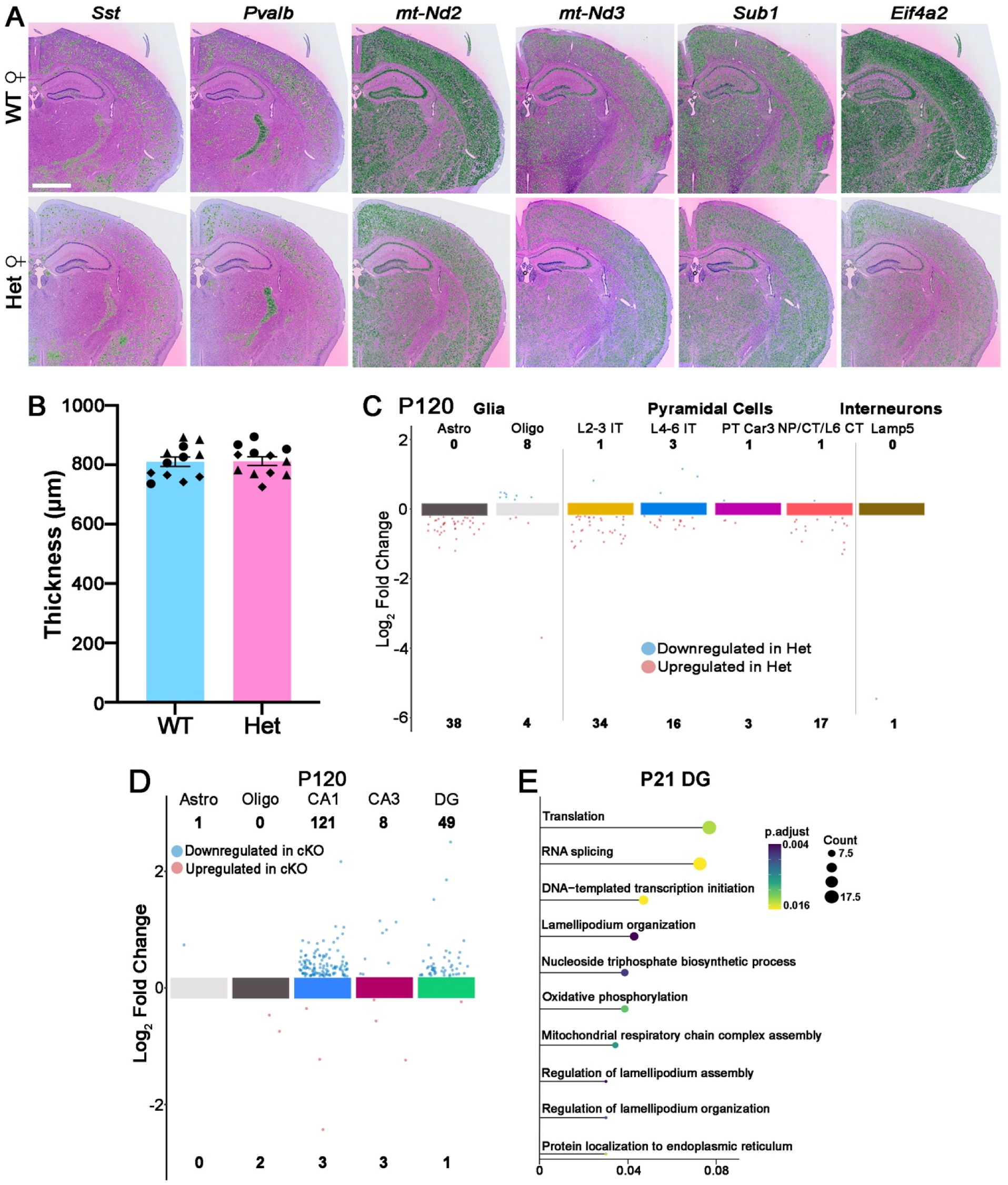
Altered gene expression in Het female mice. **A.** Spatial distribution of DEGs in P21 WT and Het female mice visualized in cell segmented spatial transcriptomics data. **B.** Cortical thickness in P21 female mice through S1. n = 3 brains/genotype, 12 slices counted for each genotype, each brain labeled with a different shape. **C.** Scatter chart showing DEGs downregulated (blue) or upregulated (red) in distinct cortical cell types in P120 Het mice. **D.** Scatter chart showing DEGs downregulated (blue) or upregulated (red) in Het mice in P120 hippocampus subdomains and cell types. **E.** clusterProfiler GO enrichment of top biological processes for DEGs of P21 DG cells between WT and Het mice. Welch’s 2-tailed t-test (B), two-sided Wilcoxon rank sum test with Bonferroni correction (C, D), and one-sided Fisher’s exact with BH correction (E) were used. Scale bar = 1 mm.

**Figure S16.**
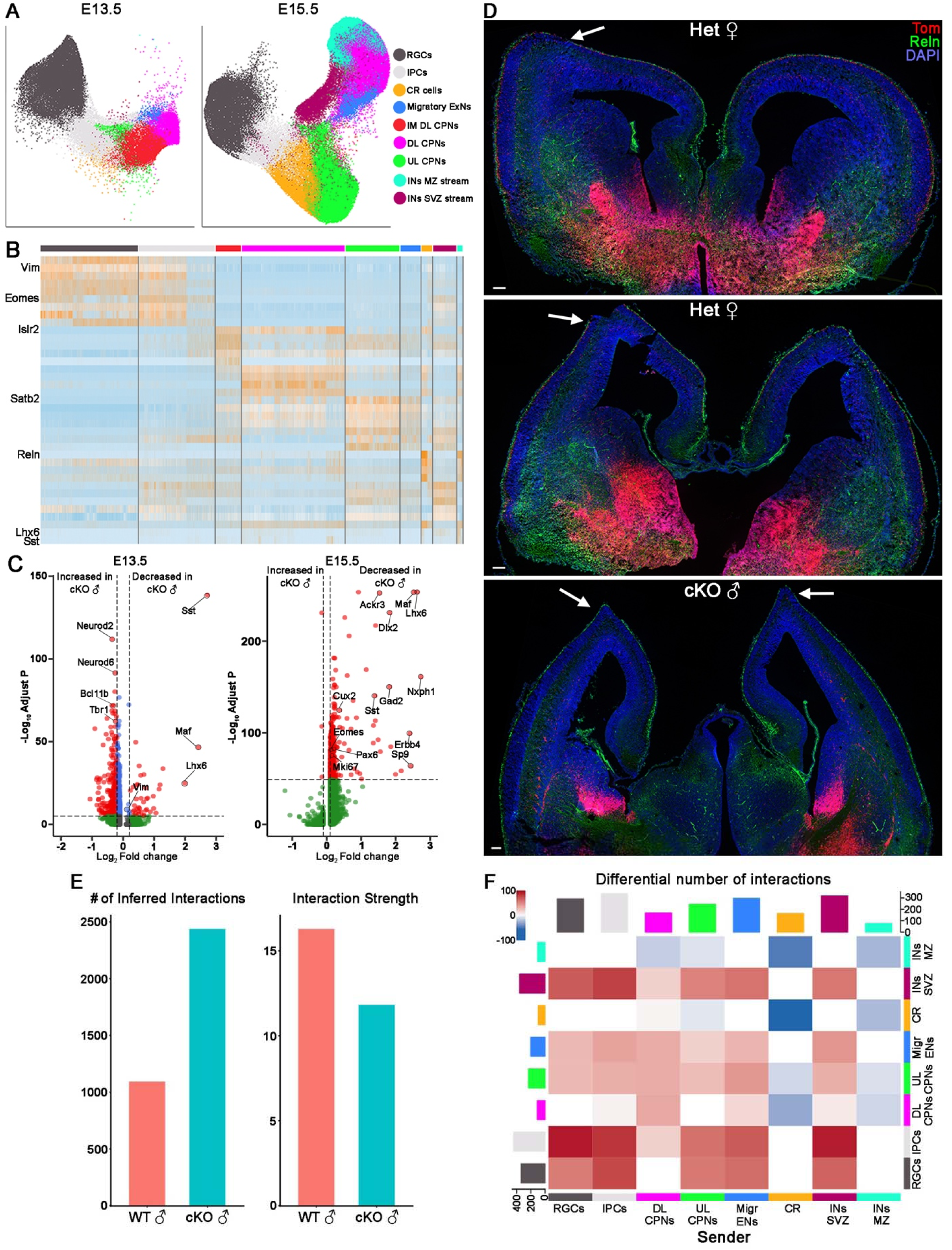
Altered gene expression and cortical abnormalities in embryonic cKO mutants. **A.** UMAP plots of E13.5 and E15.5 cortex spatial transcripts labeled by cell types. **B.** Heatmap depicting genes enriched in embryonic cortical cell types. **C.** Volcano plot depicting E13.5 (left) and E15.5 (right) cortical DEGs between WT and cKO mice. **D.** E15.5 Het female and cKO male brain sections stained for tdTomato and Reln highlighting cortical abnormalities. Scale bar = 100 μm. **E.** Graphs depicting number of inferred interactions (left) and interaction strength (right) between E15.5 male WT and cKO cortex. **F.** CellChat-derived heatmap depicting differential number of interactions between cell types from inferred cell-cell communications in E15.5 WT and cKO cortex.

## REFERENCES

1. Neri, G., Schwartz, C.E., Lubs, H.A., and Stevenson, R.E. (2018). X-linked intellectual disability update 2017. Am J Med Genet A 176, 1375–1388. 10.1002/ajmg.a.38710.

2. Tejada, M.I., and Ibarluzea, N. (2020). Non-syndromic X linked intellectual disability: Current knowledge in light of the recent advances in molecular and functional studies. Clin Genet 97, 677–687. 10.1111/cge.13698.

3. Patel, D.R., Cabral, M.D., Ho, A., and Merrick, J. (2020). A clinical primer on intellectual disability. Transl Pediatr 9, S23–S35. 10.21037/tp.2020.02.02.

4. Bernardo, P., Cuccurullo, C., Rubino, M., De Vita, G., Terrone, G., Bilo, L., and Coppola, A. (2024). X-Linked Epilepsies: A Narrative Review. Int J Mol Sci 25. 10.3390/ijms25074110.

5. Sherr, E.H. (2003). The ARX story (epilepsy, mental retardation, autism, and cerebral malformations): one gene leads to many phenotypes. Curr Opin Pediatr 15, 567–571.

6. Gecz, J., Cloosterman, D., and Partington, M. (2006). ARX: a gene for all seasons. Curr Opin Genet Dev 16, 308–316. 10.1016/j.gde.2006.04.003.

7. Friocourt, G., and Parnavelas, J.G. (2010). Mutations in ARX Result in Several Defects Involving GABAergic Neurons. Front Cell Neurosci 4, 4. 10.3389/fncel.2010.00004.

8. Shoubridge, C., Fullston, T., and Gecz, J. (2010). ARX spectrum disorders: making inroads into the molecular pathology. Hum Mutat 31, 889–900. 10.1002/humu.21288.

9. Gras, M., Heide, S., Keren, B., Valence, S., Garel, C., Whalen, S., Jansen, A.C., Keymolen, K., Stouffs, K., Jennesson, M., et al. (2024). Further characterisation of ARX-related disorders in females due to inherited or de novo variants. J Med Genet 61, 103–108. 10.1136/jmg-2023-109203.

10. Szelenyi, E.R., Fisenne, D., Knox, J.E., Harris, J.A., Gornet, J.A., Palaniswamy, R., Kim, Y., Venkataraju, K.U., and Osten, P. (2024). Distributed X chromosome inactivation in brain circuitry is associated with X-linked disease penetrance of behavior. Cell Rep 43, 114068. 10.1016/j.celrep.2024.114068.

11. Kitamura, K., Yanazawa, M., Sugiyama, N., Miura, H., Iizuka-Kogo, A., Kusaka, M., Omichi, K., Suzuki, R., Kato-Fukui, Y., Kamiirisa, K., et al. (2002). Mutation of ARX causes abnormal development of forebrain and testes in mice and X-linked lissencephaly with abnormal genitalia in humans. Nat Genet 32, 359–369.

12. Curie, A., Friocourt, G., des Portes, V., Roy, A., Nazir, T., Brun, A., Cheylus, A., Marcorelles, P., Retzepi, K., Maleki, N., et al. (2018). Basal ganglia involvement in ARX patients: The reason for ARX patients very specific grasping? Neuroimage Clin 19, 454–465. 10.1016/j.nicl.2018.04.001.

13. Colombo, E., Collombat, P., Colasante, G., Bianchi, M., Long, J., Mansouri, A., Rubenstein, J.L., and Broccoli, V. (2007). Inactivation of Arx, the murine ortholog of the X-linked lissencephaly with ambiguous genitalia gene, leads to severe disorganization of the ventral telencephalon with impaired neuronal migration and differentiation. J Neurosci 27, 4786–4798. 10.1523/JNEUROSCI.0417-07.2007.

14. Lim, Y., Akula, S.K., Myers, A.K., Chen, C., Rafael, K.A., Ibach, M.G., Trevathan, E., Walsh, C.A., Golden, J.A., and Cho, G. (2026). ARX mutation-associated interneuron defects provide insights into mechanisms underlying developmental epilepsies. Brain. 10.1093/brain/awag036.

15. Marsh, E.D., Nasrallah, M.P., Walsh, C., Murray, K.A., Nicole Sunnen, C., McCoy, A., and Golden, J.A. (2016). Developmental interneuron subtype deficits after targeted loss of Arx. BMC Neurosci 17, 35. 10.1186/s12868-016-0265-8.

16. Marcorelles, P., Laquerriere, A., Adde-Michel, C., Marret, S., Saugier-Veber, P., Beldjord, C., and Friocourt, G. (2010). Evidence for tangential migration disturbances in human lissencephaly resulting from a defect in LIS1, DCX and ARX genes. Acta Neuropathol 120, 503–515. 10.1007/s00401-010-0692-z.

17. Marsh, E., Fulp, C., Gomez, E., Nasrallah, I., Minarcik, J., Sudi, J., Christian, S.L., Mancini, G., Labosky, P., Dobyns, W., et al. (2009). Targeted loss of Arx results in a developmental epilepsy mouse model and recapitulates the human phenotype in heterozygous females. Brain 132, 1563–1576. 10.1093/brain/awp107.

18. Joseph, D.J., Von Deimling, M., Hasegawa, Y., Cristancho, A.G., Ahrens-Nicklas, R.C., Rogers, S.L., Risbud, R., McCoy, A.J., and Marsh, E.D. (2021). Postnatal Arx transcriptional activity regulates functional properties of PV interneurons. iScience 24, 101999. 10.1016/j.isci.2020.101999.

19. Kitamura, K., Itou, Y., Yanazawa, M., Ohsawa, M., Suzuki-Migishima, R., Umeki, Y., Hohjoh, H., Yanagawa, Y., Shinba, T., Itoh, M., et al. (2009). Three human ARX mutations cause the lissencephaly-like and mental retardation with epilepsy-like pleiotropic phenotypes in mice. Hum Mol Genet 18, 3708–3724. 10.1093/hmg/ddp318.

20. Price, M.G., Yoo, J.W., Burgess, D.L., Deng, F., Hrachovy, R.A., Frost, J.D., Jr., and Noebels, J.L. (2009). A triplet repeat expansion genetic mouse model of infantile spasms syndrome, Arx(GCG)10+7, with interneuronopathy, spasms in infancy, persistent seizures, and adult cognitive and behavioral impairment. J Neurosci 29, 8752–8763. 10.1523/JNEUROSCI.0915-09.2009.

21. Lee, K., Mattiske, T., Kitamura, K., Gecz, J., and Shoubridge, C. (2014). Reduced polyalanine-expanded Arx mutant protein in developing mouse subpallium alters Lmo1 transcriptional regulation. Hum Mol Genet 23, 1084–1094. 10.1093/hmg/ddt503.

22. Jackson, M.R., Lee, K., Mattiske, T., Jaehne, E.J., Ozturk, E., Baune, B.T., O’Brien, T.J., Jones, N., and Shoubridge, C. (2017). Extensive phenotyping of two ARX polyalanine expansion mutation mouse models that span clinical spectrum of intellectual disability and epilepsy. Neurobiol Dis 105, 245–256. 10.1016/j.nbd.2017.05.012.

23. Dubos, A., Meziane, H., Iacono, G., Curie, A., Riet, F., Martin, C., Loaec, N., Birling, M.C., Selloum, M., Normand, E., et al. (2018). A new mouse model of ARX dup24 recapitulates the patients’ behavioral and fine motor alterations. Hum Mol Genet 27, 2138–2153. 10.1093/hmg/ddy122.

24. Siehr, M.S., Massey, C.A., and Noebels, J.L. (2020). Arx expansion mutation perturbs cortical development by augmenting apoptosis without activating innate immunity in a mouse model of X-linked infantile spasms syndrome. Dis Model Mech 13. 10.1242/dmm.042515.

25. Nieto-Estevez, V., Varma, P., Mirsadeghi, S., Caballero, J., Gamero-Alameda, S., Hosseini, A., Silvosa, M.J., Thodeson, D.M., Goswami, S., Lybrand, Z.R., et al. (2026). Dual developmental effects of ARX poly-alanine mutations on human cortical excitatory and inhibitory neurons. Cell Rep 45, 116746. 10.1016/j.celrep.2025.116746.

26. Ding, J.W., Kim, C.N., Ostrowski, M.S., Abeykoon, Y., Pavlovic, B.J., Wallace, J.L., Schaefer, N.K., Nowakowski, T.J., and Pollen, A.A. (2026). Dissecting gene regulatory networks governing human cortical cell fate. Nature. 10.1038/s41586-025-09997-7.

27. Xu, Q., Tam, M., and Anderson, S.A. (2008). Fate mapping Nkx2.1-lineage cells in the mouse telencephalon. J Comp Neurol 506, 16–29.

28. Sommeijer, J.P., and Levelt, C.N. (2012). Synaptotagmin-2 is a reliable marker for parvalbumin positive inhibitory boutons in the mouse visual cortex. PLoS One 7, e35323. 10.1371/journal.pone.0035323.

29. Rhodes, C.T., Asokumar, D., Sohn, M., Naskar, S., Elisha, L., Stevenson, P., Lee, D.R., Zhang, Y., Rocha, P.P., Dale, R.K., et al. (2024). Loss of Ezh2 in the medial ganglionic eminence alters interneuron fate, cell morphology and gene expression profiles. Front Cell Neurosci 18, 1334244. 10.3389/fncel.2024.1334244.

30. Li, J., Tanzillo, A.F., Pizzirusso, G., Caccavano, A., Chittajallu, R., Sohn, M., Abebe, D., Zhang, Y., Pelkey, K.A., Dale, R.K., et al. (2026). Reducing methylation of histone 3.3 lysine 4 in the medial ganglionic eminence and hypothalamus recapitulates neurodevelopmental disorder phenotypes. Nat Commun 17. 10.1038/s41467-026-69248-9.

31. Ioannidou, C., Marsicano, G., and Busquets-Garcia, A. (2018). Assessing Prepulse Inhibition of Startle in Mice. Bio Protoc 8, e2789. 10.21769/BioProtoc.2789.

32. Swerdlow, N.R., Weber, M., Qu, Y., Light, G.A., and Braff, D.L. (2008). Realistic expectations of prepulse inhibition in translational models for schizophrenia research. Psychopharmacology (Berl) 199, 331–388. 10.1007/s00213-008-1072-4.

33. Pai, E.L., Chen, J., Fazel Darbandi, S., Cho, F.S., Chen, J., Lindtner, S., Chu, J.S., Paz, J.T., Vogt, D., Paredes, M.F., and Rubenstein, J.L. (2020). Maf and Mafb control mouse pallial interneuron fate and maturation through neuropsychiatric disease gene regulation. Elife 9. 10.7554/eLife.54903.

34. Fragkouli, A., van Wijk, N.V., Lopes, R., Kessaris, N., and Pachnis, V. (2009). LIM homeodomain transcription factor-dependent specification of bipotential MGE progenitors into cholinergic and GABAergic striatal interneurons. Development 136, 3841–3851. 10.1242/dev.038083.

35. Flandin, P., Zhao, Y., Vogt, D., Jeong, J., Long, J., Potter, G., Westphal, H., and Rubenstein, J.L. (2011). Lhx6 and Lhx8 coordinately induce neuronal expression of Shh that controls the generation of interneuron progenitors. Neuron 70, 939–950. 10.1016/j.neuron.2011.04.020.

36. Chen, Y.J., Friedman, B.A., Ha, C., Durinck, S., Liu, J., Rubenstein, J.L., Seshagiri, S., and Modrusan, Z. (2017). Single-cell RNA sequencing identifies distinct mouse medial ganglionic eminence cell types. Sci Rep 7, 45656. 10.1038/srep45656.

37. McKenzie, O., Ponte, I., Mangelsdorf, M., Finnis, M., Colasante, G., Shoubridge, C., Stifani, S., Gecz, J., and Broccoli, V. (2007). Aristaless-related homeobox gene, the gene responsible for West syndrome and related disorders, is a Groucho/transducin-like enhancer of split dependent transcriptional repressor. Neuroscience 146, 236–247. 10.1016/j.neuroscience.2007.01.038.

38. Fulp, C.T., Cho, G., Marsh, E.D., Nasrallah, I.M., Labosky, P.A., and Golden, J.A. (2008). Identification of Arx transcriptional targets in the developing basal forebrain. Hum Mol Genet 17, 3740–3760. 10.1093/hmg/ddn271.

39. Colasante, G., Sessa, A., Crispi, S., Calogero, R., Mansouri, A., Collombat, P., and Broccoli, V. (2009). Arx acts as a regional key selector gene in the ventral telencephalon mainly through its transcriptional repression activity. Dev Biol 334, 59–71. 10.1016/j.ydbio.2009.07.014.

40. Abe, H., Okazawa, M., and Nakanishi, S. (2011). The Etv1/Er81 transcription factor orchestrates activity-dependent gene regulation in the terminal maturation program of cerebellar granule cells. Proc Natl Acad Sci U S A 108, 12497–12502. 10.1073/pnas.1109940108.

41. Wang, Y., Li, G., Stanco, A., Long, J.E., Crawford, D., Potter, G.B., Pleasure, S.J., Behrens, T., and Rubenstein, J.L. (2011). CXCR4 and CXCR7 have distinct functions in regulating interneuron migration. Neuron 69, 61–76. 10.1016/j.neuron.2010.12.005.

42. Golonzhka, O., Nord, A., Tang, P.L.F., Lindtner, S., Ypsilanti, A.R., Ferretti, E., Visel, A., Selleri, L., and Rubenstein, J.L.R. (2015). Pbx Regulates Patterning of the Cerebral Cortex in Progenitors and Postmitotic Neurons. Neuron 88, 1192–1207. 10.1016/j.neuron.2015.10.045.

43. Hanley, O., Zewdu, R., Cohen, L.J., Jung, H., Lacombe, J., Philippidou, P., Lee, D.H., Selleri, L., and Dasen, J.S. (2016). Parallel Pbx-Dependent Pathways Govern the Coalescence and Fate of Motor Columns. Neuron 91, 1005–1020. 10.1016/j.neuron.2016.07.043.

44. Sass, J.O., Fischer, K., Wang, R., Christensen, E., Scholl-Burgi, S., Chang, R., Kapelari, K., and Walter, M. (2010). D-glyceric aciduria is caused by genetic deficiency of D-glycerate kinase (GLYCTK). Hum Mutat 31, 1280–1285. 10.1002/humu.21375.

45. Singh, V., and Auerbach, D.S. (2024). Neurocardiac pathologies associated with potassium channelopathies. Epilepsia 65, 2537–2552. 10.1111/epi.18066.

46. Schubert, T., and Schaaf, C.P. (2025). MAGEL2 (patho-)physiology and Schaaf-Yang syndrome. Dev Med Child Neurol 67, 35–48. 10.1111/dmcn.16018.

47. Mei, L., and Xiong, W.-C. (2008). Neuregulin 1 in neural development, synaptic plasticity and schizophrenia. Nature reviews Neuroscience 9, 437–452. 10.1038/nrn2392.

48. Sanchez-Alcaniz, J.A., Haege, S., Mueller, W., Pla, R., Mackay, F., Schulz, S., Lopez-Bendito, G., Stumm, R., and Marin, O. (2011). Cxcr7 controls neuronal migration by regulating chemokine responsiveness. Neuron 69, 77–90. 10.1016/j.neuron.2010.12.006.

49. Cholfin, J.A., and Rubenstein, J.L. (2008). Frontal cortex subdivision patterning is coordinately regulated by Fgf8, Fgf17, and Emx2. J Comp Neurol 509, 144–155. 10.1002/cne.21709.

50. Singhal, V., Chou, N., Lee, J., Yue, Y., Liu, J., Chock, W.K., Lin, L., Chang, Y.C., Teo, E.M.L., Aow, J., et al. (2024). BANKSY unifies cell typing and tissue domain segmentation for scalable spatial omics data analysis. Nat Genet 56, 431–441. 10.1038/s41588-024-01664-3.

51. Cable, D.M., Murray, E., Zou, L.S., Goeva, A., Macosko, E.Z., Chen, F., and Irizarry, R.A. (2022). Robust decomposition of cell type mixtures in spatial transcriptomics. Nat Biotechnol 40, 517–526. 10.1038/s41587-021-00830-w.

52. Yao, Z., van Velthoven, C.T.J., Nguyen, T.N., Goldy, J., Sedeno-Cortes, A.E., Baftizadeh, F., Bertagnolli, D., Casper, T., Chiang, M., Crichton, K., et al. (2021). A taxonomy of transcriptomic cell types across the isocortex and hippocampal formation. Cell 184, 3222–3241 e3226. 10.1016/j.cell.2021.04.021.

53. West, A.K., Hidalgo, J., Eddins, D., Levin, E.D., and Aschner, M. (2008). Metallothionein in the central nervous system: Roles in protection, regeneration and cognition. Neurotoxicology 29, 489–503. 10.1016/j.neuro.2007.12.006.

54. Ralhan, I., Do, A.D., Bae, J.Y., Feringa, F.M., Cai, W., Chang, J., Chik, K., Lee, N.Y.J., Gerry, C.J., van der Kant, R., et al. (2026). Protective ApoE variants support neuronal function by effluxing oxidized phospholipids. Neuron 114, 661–678 e610. 10.1016/j.neuron.2025.10.040.

55. Pelkey, K.A., Calvigioni, D., Fang, C., Vargish, G., Ekins, T., Auville, K., Wester, J.C., Lai, M., Mackenzie-Gray Scott, C., Yuan, X., et al. (2020). Paradoxical network excitation by glutamate release from VGluT3(+) GABAergic interneurons. Elife 9. 10.7554/eLife.51996.

56. Dudok, B., Klein, P.M., Hwaun, E., Lee, B.R., Yao, Z., Fong, O., Bowler, J.C., Terada, S., Sparks, F.T., Szabo, G.G., et al. (2021). Alternating sources of perisomatic inhibition during behavior. Neuron 109, 997–1012 e1019. 10.1016/j.neuron.2021.01.003.

57. Oh, S., Park, H., and Zhang, X. (2021). Hybrid Clustering of Single-Cell Gene Expression and Spatial Information via Integrated NMF and K-Means. Front Genet 12, 763263. 10.3389/fgene.2021.763263.

58. Liu, L., Wang, J., Liu, X., Wang, J., Chen, L., Zhu, H., Mai, J., Hu, T., and Liu, S. (2024). Prenatal prevalence and postnatal manifestations of 16p11.2 deletions: A new insights into neurodevelopmental disorders based on clinical investigations combined with multi-omics analysis. Clin Chim Acta 552, 117671. 10.1016/j.cca.2023.117671.

59. Baulac, M. (2015). MTLE with hippocampal sclerosis in adult as a syndrome. Rev Neurol (Paris) 171, 259–266. 10.1016/j.neurol.2015.02.004.

60. Selten, M., Bernard, C., Mukherjee, D., Hamid, F., Hanusz-Godoy, A., Oozeer, F., Zimmer, C., and Marin, O. (2025). Regulation of PV interneuron plasticity by neuropeptide-encoding genes. Nature 643, 173–181. 10.1038/s41586-025-08933-z.

61. Ganfornina, M.D., Do Carmo, S., Martinez, E., Tolivia, J., Navarro, A., Rassart, E., and Sanchez, D. (2010). ApoD, a glia-derived apolipoprotein, is required for peripheral nerve functional integrity and a timely response to injury. Glia 58, 1320–1334. 10.1002/glia.21010.

62. Bohrer, C., Pfurr, S., Mammadzada, K., Schildge, S., Plappert, L., Hils, M., Pous, L., Rauch, K.S., Dumit, V.I., Pfeifer, D., et al. (2015). The balance of Id3 and E47 determines neural stem/precursor cell differentiation into astrocytes. EMBO J 34, 2804–2819. 10.15252/embj.201591118.

63. Chen, Z.P., Wang, S., Zhao, X., Fang, W., Wang, Z., Ye, H., Wang, M.J., Ke, L., Huang, T., Lv, P., et al. (2023). Lipid-accumulated reactive astrocytes promote disease progression in epilepsy. Nat Neurosci 26, 542–554. 10.1038/s41593-023-01288-6.

64. Scharfman, H.E. (2005). Brain-derived neurotrophic factor and epilepsy--a missing link? Epilepsy Curr 5, 83–88. 10.1111/j.1535-7511.2005.05312.x.

65. Busch, R.M., Yehia, L., Blumcke, I., Hu, B., Prayson, R., Hermann, B.P., Najm, I.M., and Eng, C. (2022). Molecular and subregion mechanisms of episodic memory phenotypes in temporal lobe epilepsy. Brain Commun 4, fcac285. 10.1093/braincomms/fcac285.

66. Bozdagi, O., Rich, E., Tronel, S., Sadahiro, M., Patterson, K., Shapiro, M.L., Alberini, C.M., Huntley, G.W., and Salton, S.R. (2008). The neurotrophin-inducible gene Vgf regulates hippocampal function and behavior through a brain-derived neurotrophic factor-dependent mechanism. J Neurosci 28, 9857–9869. 10.1523/JNEUROSCI.3145-08.2008.

67. Hermey, G., Plath, N., Hubner, C.A., Kuhl, D., Schaller, H.C., and Hermans-Borgmeyer, I. (2004). The three sorCS genes are differentially expressed and regulated by synaptic activity. J Neurochem 88, 1470–1476. 10.1046/j.1471-4159.2004.02286.x.

68. Gomez de San Jose, N., Massa, F., Halbgebauer, S., Oeckl, P., Steinacker, P., and Otto, M. (2022). Neuronal pentraxins as biomarkers of synaptic activity: from physiological functions to pathological changes in neurodegeneration. J Neural Transm (Vienna) 129, 207–230. 10.1007/s00702-021-02411-2.

69. Singec, I., Knoth, R., Ditter, M., Hagemeyer, C.E., Rosenbrock, H., Frotscher, M., and Volk, B. (2002). Synaptic vesicle protein synaptoporin is differently expressed by subpopulations of mouse hippocampal neurons. J Comp Neurol 452, 139–153. 10.1002/cne.10371.

70. Lee, K.J., Queenan, B.N., Rozeboom, A.M., Bellmore, R., Lim, S.T., Vicini, S., and Pak, D.T. (2013). Mossy fiber-CA3 synapses mediate homeostatic plasticity in mature hippocampal neurons. Neuron 77, 99–114. 10.1016/j.neuron.2012.10.033.

71. Lee, S.Y., and Soltesz, I. (2011). Cholecystokinin: a multi-functional molecular switch of neuronal circuits. Dev Neurobiol 71, 83–91. 10.1002/dneu.20815.

72. Avanzini, G., and Franceschetti, S. (2003). Cellular biology of epileptogenesis. Lancet Neurol 2, 33–42. 10.1016/s1474-4422(03)00265-5.

73. Ruttimann, E., Vacher, C.M., Gassmann, M., Kaupmann, K., Van der Putten, H., and Bettler, B. (2004). Altered hippocampal expression of calbindin-D-28k and calretinin in GABA(B(1))-deficient mice. Biochem Pharmacol 68, 1613–1620. 10.1016/j.bcp.2004.07.019.

74. Oyang, E.L., Davidson, B.C., Lee, W., and Poon, M.M. (2011). Functional characterization of the dendritically localized mRNA neuronatin in hippocampal neurons. PLoS One 6, e24879. 10.1371/journal.pone.0024879.

75. Hartmann, U., Hulsmann, H., Seul, J., Roll, S., Midani, H., Breloy, I., Hechler, D., Muller, R., and Paulsson, M. (2013). Testican-3: a brain-specific proteoglycan member of the BM-40/SPARC/osteonectin family. J Neurochem 125, 399–409. 10.1111/jnc.12212.

76. Wang, Y., Yang, H., Li, N., Wang, L., Guo, C., Ma, W., Liu, S., Peng, C., Chen, J., Song, H., et al. (2024). A Novel Ubiquitin Ligase Adaptor PTPRN Suppresses Seizure Susceptibility through Endocytosis of Na(V)1.2 Sodium Channels. Adv Sci (Weinh) 11, e2400560. 10.1002/advs.202400560.

77. Wu, H., Suo, G., Li, T., Zheng, Y., Li, H., Shen, F., Wang, Y., Ni, H., and Wu, Y. (2022). CaMKIV mediates spine growth deficiency of hippocampal neurons by regulation of EGR3/BDNF signal axis in congenital hypothyroidism. Cell Death Discov 8, 482. 10.1038/s41420-022-01270-4.

78. Nakaya, N., Sultana, A., Lee, H.S., and Tomarev, S.I. (2012). Olfactomedin 1 interacts with the Nogo A receptor complex to regulate axon growth. J Biol Chem 287, 37171–37184. 10.1074/jbc.M112.389916.

79. Rak, M., Benit, P., Chretien, D., Bouchereau, J., Schiff, M., El-Khoury, R., Tzagoloff, A., and Rustin, P. (2016). Mitochondrial cytochrome c oxidase deficiency. Clin Sci (Lond) 130, 393–407. 10.1042/CS20150707.

80. Wu, S.J., Dai, M., Yang, S.P., McCann, C., Qiu, Y., Kumar, V., Marrero, G.J., Tsyporin, J., Huang, S., Shin, D., et al. (2026). Pyramidal neurons proportionately alter cortical interneuron subtypes. Nature. 10.1038/s41586-025-09996-8.

81. Curmi, P.A., Gavet, O., Charbaut, E., Ozon, S., Lachkar-Colmerauer, S., Manceau, V., Siavoshian, S., Maucuer, A., and Sobel, A. (1999). Stathmin and its phosphoprotein family: general properties, biochemical and functional interaction with tubulin. Cell Struct Funct 24, 345–357. 10.1247/csf.24.345.

82. Drongitis, D., Caterino, M., Verrillo, L., Santonicola, P., Costanzo, M., Poeta, L., Attianese, B., Barra, A., Terrone, G., Lioi, M.B., et al. (2022). Deregulation of microtubule organization and RNA metabolism in Arx models for lissencephaly and developmental epileptic encephalopathy. Hum Mol Genet 31, 1884–1908. 10.1093/hmg/ddac028.

83. Paul, M.S., Duncan, A.R., Genetti, C.A., Pan, H., Jackson, A., Grant, P.E., Shi, J., Pinelli, M., Brunetti-Pierri, N., Garza-Flores, A., et al. (2023). Rare EIF4A2 variants are associated with a neurodevelopmental disorder characterized by intellectual disability, hypotonia, and epilepsy. Am J Hum Genet 110, 120–145. 10.1016/j.ajhg.2022.11.011.

84. Villalba, A., Gotz, M., and Borrell, V. (2021). The regulation of cortical neurogenesis. Curr Top Dev Biol 142, 1–66. 10.1016/bs.ctdb.2020.10.003.

85. Bright, A.R., Kotlyarenko, Y., Neuhaus, F., Rodrigues, D., Feng, C., Peters, C., Vitali, I., Donmez, E., Myoga, M.H., Dvoretskova, E., and Mayer, C. (2025). Temporal control of progenitor competence shapes maturation in GABAergic neuron development in mice. Nat Neurosci. 10.1038/s41593-025-01999-y.

86. Silva, C.G., Peyre, E., Adhikari, M.H., Tielens, S., Tanco, S., Van Damme, P., Magno, L., Krusy, N., Agirman, G., Magiera, M.M., et al. (2018). Cell-Intrinsic Control of Interneuron Migration Drives Cortical Morphogenesis. Cell 172, 1063–1078 e1019. 10.1016/j.cell.2018.01.031.

87. Vilchez-Acosta, A., Manso, Y., Cardenas, A., Elias-Tersa, A., Martinez-Losa, M., Pascual, M., Alvarez-Dolado, M., Nairn, A.C., Borrell, V., and Soriano, E. (2022). Specific contribution of Reelin expressed by Cajal-Retzius cells or GABAergic interneurons to cortical lamination. Proc Natl Acad Sci U S A 119, e2120079119. 10.1073/pnas.2120079119.

88. Reichard, J., Wolff, P., Xie, S., Zuo, K., Fullio, C.L., Du, J., Graff, S., Linde, J., Yildiz, C.B., Pitschelatow, G., et al. (2025). DNMT1-mediated regulation of somatostatin-positive interneuron migration impacts cortical architecture and function. Nat Commun 16, 6834. 10.1038/s41467-025-62114-0.

89. Rodgers, J., Calvert, S., Shoubridge, C., and McGaughran, J. (2021). A novel ARX loss of function variant in female monozygotic twins is associated with chorea. Eur J Med Genet 64, 104315. 10.1016/j.ejmg.2021.104315.

90. Sciuto, L., Fichera, V., Zanghì, A., Vecchio, M., Falsaperla, R., Galioto, S., Palmucci, S., Belfiore, G., Di Napoli, C., Polizzi, A., and Praticò, A.D. (2024). Lissencephaly, Pachygyrias, Band Heterotopias, RELN Pathway, and Mutations (Incomplete Neuron Migration). J Pediatr Neurol 22, 377–386. 10.1055/s-0044-1786790.

91. Jin, S., Plikus, M.V., and Nie, Q. (2025). CellChat for systematic analysis of cell-cell communication from single-cell transcriptomics. Nat Protoc 20, 180–219. 10.1038/s41596-024-01045-4.

92. Reber, M., Hindges, R., and Lemke, G. (2007). Eph receptors and ephrin ligands in axon guidance. Adv Exp Med Biol 621, 32–49. 10.1007/978-0-387-76715-4_3.

93. Gurrapu, S., and Tamagnone, L. (2016). Transmembrane semaphorins: Multimodal signaling cues in development and cancer. Cell Adh Migr 10, 675–691. 10.1080/19336918.2016.1197479.

94. Wareham, L.K., Baratta, R.O., Del Buono, B.J., Schlumpf, E., and Calkins, D.J. (2024). Collagen in the central nervous system: contributions to neurodegeneration and promise as a therapeutic target. Mol Neurodegener 19, 11. 10.1186/s13024-024-00704-0.

95. Langenhan, T., Piao, X., and Monk, K.R. (2016). Adhesion G protein-coupled receptors in nervous system development and disease. Nat Rev Neurosci 17, 550–561. 10.1038/nrn.2016.86.

96. Li, S., Jin, Z., Koirala, S., Bu, L., Xu, L., Hynes, R.O., Walsh, C.A., Corfas, G., and Piao, X. (2008). GPR56 regulates pial basement membrane integrity and cortical lamination. J Neurosci 28, 5817–5826. 10.1523/JNEUROSCI.0853-08.2008.

97. Luo, R., Jeong, S.J., Jin, Z., Strokes, N., Li, S., and Piao, X. (2011). G protein-coupled receptor 56 and collagen III, a receptor-ligand pair, regulates cortical development and lamination. Proc Natl Acad Sci U S A 108, 12925–12930. 10.1073/pnas.1104821108.

98. Vogt, D., Hunt, R.F., Mandal, S., Sandberg, M., Silberberg, S.N., Nagasawa, T., Yang, Z., Baraban, S.C., and Rubenstein, J.L. (2014). Lhx6 directly regulates Arx and CXCR7 to determine cortical interneuron fate and laminar position. Neuron 82, 350–364. 10.1016/j.neuron.2014.02.030.

99. Winston, S.M., Hayward, M.D., Nestler, E.J., and Duman, R.S. (1990). Chronic electroconvulsive seizures down-regulate expression of the immediate-early genes c-fos and c-jun in rat cerebral cortex. J Neurochem 54, 1920–1925. 10.1111/j.1471-4159.1990.tb04892.x.

100. Calais, J.B., Valvassori, S.S., Resende, W.R., Feier, G., Athie, M.C., Ribeiro, S., Gattaz, W.F., Quevedo, J., and Ojopi, E.B. (2013). Long-term decrease in immediate early gene expression after electroconvulsive seizures. J Neural Transm (Vienna) 120, 259–266. 10.1007/s00702-012-0861-4.

101. Imoto, Y., Segi-Nishida, E., Suzuki, H., and Kobayashi, K. (2017). Rapid and stable changes in maturation-related phenotypes of the adult hippocampal neurons by electroconvulsive treatment. Mol Brain 10, 8. 10.1186/s13041-017-0288-9.

102. Corbett, B.F., You, J.C., Zhang, X., Pyfer, M.S., Tosi, U., Iascone, D.M., Petrof, I., Hazra, A., Fu, C.H., Stephens, G.S., et al. (2017). DeltaFosB Regulates Gene Expression and Cognitive Dysfunction in a Mouse Model of Alzheimer’s Disease. Cell Rep 20, 344–355. 10.1016/j.celrep.2017.06.040.

103. Murano, T., Hagihara, H., Tajinda, K., Matsumoto, M., and Miyakawa, T. (2019). Transcriptomic immaturity inducible by neural hyperexcitation is shared by multiple neuropsychiatric disorders. Commun Biol 2, 32. 10.1038/s42003-018-0277-2.

104. Murano, T., Hagihara, H., Tajinda, K., Takao, K., Takamiya, Y., Katoh, K., Robison, A.J., Matsumoto, M., Namihira, M., and Miyakawa, T. (2026). Repetitive neuronal activation regulates cellular maturation state via nuclear reprogramming. Nat Commun 17. 10.1038/s41467-026-74202-w.

105. Di Donato, N., Timms, A.E., Aldinger, K.A., Mirzaa, G.M., Bennett, J.T., Collins, S., Olds, C., Mei, D., Chiari, S., Carvill, G., et al. (2018). Analysis of 17 genes detects mutations in 81% of 811 patients with lissencephaly. Genet Med 20, 1354–1364. 10.1038/gim.2018.8.

106. Haverfield, E.V., Whited, A.J., Petras, K.S., Dobyns, W.B., and Das, S. (2009). Intragenic deletions and duplications of the LIS1 and DCX genes: a major disease-causing mechanism in lissencephaly and subcortical band heterotopia. Eur J Hum Genet 17, 911–918. 10.1038/ejhg.2008.213.

107. Madisen, L., Zwingman, T.A., Sunkin, S.M., Oh, S.W., Zariwala, H.A., Gu, H., Ng, L.L., Palmiter, R.D., Hawrylycz, M.J., Jones, A.R., et al. (2010). A robust and high-throughput Cre reporting and characterization system for the whole mouse brain. Nat Neurosci 13, 133–140. 10.1038/nn.2467.

108. Mo, A., Mukamel, E.A., Davis, F.P., Luo, C., Henry, G.L., Picard, S., Urich, M.A., Nery, J.R., Sejnowski, T.J., Lister, R., et al. (2015). Epigenomic Signatures of Neuronal Diversity in the Mammalian Brain. Neuron 86, 1369–1384. 10.1016/j.neuron.2015.05.018.

109. Shimada, T., and Yamagata, K. (2018). Pentylenetetrazole-Induced Kindling Mouse Model. J Vis Exp. 10.3791/56573.

110. Lee, D.R., Zhang, Y., Rhodes, C.T., and Petros, T.J. (2023). Generation of single-cell and single-nuclei suspensions from embryonic and adult mouse brains. STAR Protoc 4, 101944. 10.1016/j.xpro.2022.101944.

111. Satija, R., Farrell, J.A., Gennert, D., Schier, A.F., and Regev, A. (2015). Spatial reconstruction of single-cell gene expression data. Nat Biotechnol 33, 495–502. 10.1038/nbt.3192.

112. Stuart, T., Srivastava, A., Madad, S., Lareau, C.A., and Satija, R. (2021). Single-cell chromatin state analysis with Signac. Nat Methods 18, 1333–1341. 10.1038/s41592-021-01282-5.

113. Hao, Y., Hao, S., Andersen-Nissen, E., Mauck, W.M., 3rd, Zheng, S., Butler, A., Lee, M.J., Wilk, A.J., Darby, C., Zager, M., et al. (2021). Integrated analysis of multimodal single-cell data. Cell 184, 3573–3587 e3529. 10.1016/j.cell.2021.04.048.

114. Cao, J., Spielmann, M., Qiu, X., Huang, X., Ibrahim, D.M., Hill, A.J., Zhang, F., Mundlos, S., Christiansen, L., Steemers, F.J., et al. (2019). The single-cell transcriptional landscape of mammalian organogenesis. Nature 566, 496–502. 10.1038/s41586-019-0969-x.

115. Xu, S., Hu, E., Cai, Y., Xie, Z., Luo, X., Zhan, L., Tang, W., Wang, Q., Liu, B., Wang, R., et al. (2024). Using clusterProfiler to characterize multiomics data. Nat Protoc 19, 3292–3320. 10.1038/s41596-024-01020-z.

